# Beyond RECIST: mathematical modeling and Bayesian inference reveal the importance of immune suppressive parameters in metastatic breast cancer

**DOI:** 10.1101/2025.08.11.669777

**Authors:** Jesse Kreger, Edgar Gonzalez, Xiaojun Wu, Jonathan A. Martinez, Evanthia T. Roussos Torres, Adam L. MacLean

## Abstract

Successful immunotherapies must overcome patient- and organ-specific tumor heterogeneities to mount an effective response. Yet tumor dynamics remain poorly characterized in organ-specific contexts and associated immune environments. To quantify heterogeneous tumor responses, we developed methods to fit mathematical models of the tumor-immune dynamics to patients undergoing combination therapy for metastatic breast cancer: checkpoint inhibition via nivolumab + ipilimumab combined with entinostat, as measured by RECIST criteria. In a subset of patients additional immune dynamics were quantified by spatial proteomics. Bayesian parameter inference revealed that only immune-modulatory parameters controlled response; parameters controlling cytotoxicity were uninformative. Through posterior parameter sampling and simulation, we created virtual tumor cohorts, enabling extrapolation beyond the data to predict probabilities of response in metastatic lesions where no data exist. We validated predictions from our virtual tumor population using held-out data characterizing off-target lesions from the patient cohort. Profile-wise likelihood analysis revealed that scans in the week immediately following treatment hold particularly high value in identifying the tumor dynamics. Overall, we demonstrate how through modeling & inference cohort size limitations can be overcome through the creation of virtual tumor populations, giving insight into the site-specific mechanisms of disease progression and response.

## Introduction

Modulating the host immune system to help it mount an effective antitumor response offers great potential. Immune checkpoint inhibitors (ICIs) seek to activate a T cell response against tumor cells by blocking T cell checkpoints including CTLA-4 or PD-1^1^, and have become standard of care for many patients. While ICI treatment alone has proven successful in certain cases for the treatment of solid tumors, e.g. lung or melanoma ^2,3^, single-agent ICI therapy has to-date been ineffective for the treatment of breast cancers confined to the breast or with distant metastases ^4,5^. In recent preclinical and clinical studies, we have observed that modulating the epigenome via histone deacetylase inhibitors (HDACi) such as entinostat may be sufficient to overcome barriers to effective ICI therapy for breast cancer^6,7,8,9,10^. Encouraging data from a phase 1b trial for a metastatic breast cancer cohort (NCI-9844: nivolumab + ipilimumab + entinostat) revealed a 25% response rate ^11^, but the small cohort size hampers thorough analyses of the mechanisms of response. Compounding the small cohort size, sources of heterogeneity emerging at multiple biological scales preclude robust evaluation of site-specific tumor dynamics in responders vs non-responders. These sources include breast cancer-specific subtype heterogeneity, patient heterogeneity, effects due to different sites of metastatic lesions, and heterogeneity due to the myriad, complex signals that comprise the tumor microenvironment (TME) at the site of any lesion ^12,13,14,15^. Generally, phase Ib trials can also include cohorts with different cancers present: the original cohort of the NCI-9844 trial included seven different cancer types.

Mathematical oncology and cancer systems immunology approaches ^16,17^ offer means to explain and predict phenomena underlying tumor progression and responses to treatment^18,19,20,21,22,12,23,24^. In cases where data are sparse, these are particularly valuable as means to extrapolate beyond the data to predict patient response to treatment. In Kreger et al. ^18^ we developed a differential equation model which, fitted to clinical trial tumor data ^25^, demonstrated the importance of myeloid-derived suppressor cell (MDSC) dynamics in controlling nascent metastatic tumor growth and predicted interventions that could promote shifts toward less immune-suppressed states.

In the current work we develop a new model informed by our previous work^18^ that enables the investigation of individual patient tumor dynamics in response to treatment with ICIs + HDACi^11^. We fit models to temporal data characterizing changes in tumor size as quantified by imaging via RECIST (‘Response Evaluation Criteria in Solid Tumors’) to study not only the dynamics of individual patients, but also the differences in dynamics between metastatic lesions within a patient. In doing so we can quantify both the inter- and the intra-patient tumor response het-erogeneity. By fitting patient-specific or lesion-specific models (variants of virtual patients ^13,26^ or digital twins ^27,28,29^), we develop a systematic framework way to simulate in light of new information (e.g. different immune parameters or a new metastatic site). Through the *in silico* simulation of tumor dynamics and the resulting predicted patient responses (using RECIST criteria) we are able to go *beyond* RECIST^30,31^ and provide means to understand the effects of new treatments on tumor-immune dynamics in cases where data are sparse and limited.

The remainder of this paper is organized as follows. We develop a flexible pipeline for the inference of patient- and site-specific tumor-immune parameters in Julia ^32^, fit to individual tumors from NCI-9844^11^. The fitted posterior parameter distributions characterizing tumor growth or response are used to identify the relative importance of the TME: we discover that only immunosuppressive (and not cytotoxic) parameters are predictive of response. We then develop methods to create virtual patient cohorts by sampling from posterior distributions, in a patient-specific manner informed by spatial proteomics, which facilitate analysis of *in silico* tumor populations across different site and treatment groups. From these we can predict responses unseen in the clinical cohort available. Finally, via analysis of the fitted models using a profile likelihood approach we discover that parameters are identifiable with limited additional cost given sufficient measurements of the tumor size in the time period immediately following treatment initiation.

## Results

### Individual breast cancer metastatic responses vary widely following triple combination therapy

We studied the tumor-immune parameters controlling responses to a new triple combination therapy (immune checkpoint inhibitors (ICIs) nivolumab and ipilimumab + entinostat) in site-specific breast cancer metastases from individual patient lesions. The phase 1b cohort NCI-9844^11^ consists of 55 target lesions from 20 patients (Fig. 1A-B and Table 1). The mean number of tumors per patient is 2.75 (Fig S1); each tumor has a minimum of two measurements. Quantified by RECIST (Response Evaluation Criteria in Solid Tumors), the mean number of measurements per tumor was 3.75 (Fig. S2) for lesions spanning 11 general sites (Figs. S3-S5). Of the 20 patients, 10 were hormone receptor-positive (HR^+^) and 10 were triple-negative (TNBC). Characterization of the individual dynamics of each of the 55 tumors is available in the Supplementary Material. As per RECIST, the same lesions are followed over time and were designated as target lesions or non-target lesions by study radiologists. Tumor sizes were also determined by study radiologists per RECIST guidelines and reported per trial protocol ^11^.

**Table 1:** Clinical characteristics of patient cohort from Roussos Torres et al.^11^. ORR: Overall response rate; OS: Overall Survival; IMC: Imaging mass cytometry.

| ID | Subtype | Best Response | ORR (RECIST) | OS (months) | IMC Available | # Tumors Followed | Fraction of Tumors that Responded |
| --- | --- | --- | --- | --- | --- | --- | --- |
| R-1 | TNBC | SD | Non-responder | 50.92 | - | 4 | 0.75 |
| R-2 | TNBC | PR | Responder | 19.58 | Yes | 3 | 1.0 |
| R-4 | HR<sup>+</sup> | PD | Non-responder | 13.34 | - | 3 | 0.33 |
| R-5 | TNBC | CR | Responder | 52.14 | Yes | 4 | 1.0 |
| R-6 | TNBC | SD | Non-responder | 2.83 | Yes | 3 | 0.0 |
| R-7 | TNBC | SD | Non-responder | 11.20 | - | 3 | 0.66 |
| R-8 | TNBC | PD | Non-responder | 8.08 | Yes | 3 | 0.66 |
| R-9 | HR<sup>+</sup> | PD | Non-responder | 1.71 | - | 3 | 0.33 |
| R-10 | HR<sup>+</sup> | SD | Non-responder | 46.85 | Yes | 3 | 0.66 |
| R-12 | HR<sup>+</sup> | PD | Non-responder | 3.45 | - | 2 | 0.0 |
| R-13 | HR<sup>+</sup> | SD | Non-responder | 4.80 | Yes | 2 | 0.0 |
| R-14 | TNBC | PR | Responder | 24.38 | - | 1 | 1.0 |
| R-15 | TNBC | SD | Non-responder | 16.76 | - | 1 | 0.0 |
| R-16 | TNBC | PR | Responder | 24.02 | Yes | 4 | 1.0 |
| R-17 | TNBC | PD | Non-responder | 11.99 | Yes | 3 | 0.33 |
| R-18 | HR<sup>+</sup> | PR | Responder | 67.35 | Yes | 4 | 1.0 |
| R-19 | HR<sup>+</sup> | PD | Non-responder | 11.66 | - | 4 | 0.5 |
| R-20 | HR<sup>+</sup> | SD | Non-responder | 7.33 | Yes | 1 | 1.0 |
| R-23 | HR<sup>+</sup> | SD | Non-responder | 5.03 | - | 2 | 0.0 |
| R-24 | HR<sup>+</sup> | SD | Non-responder | 6.74 | - | 2 | 0.0 |

**Figure 1:**
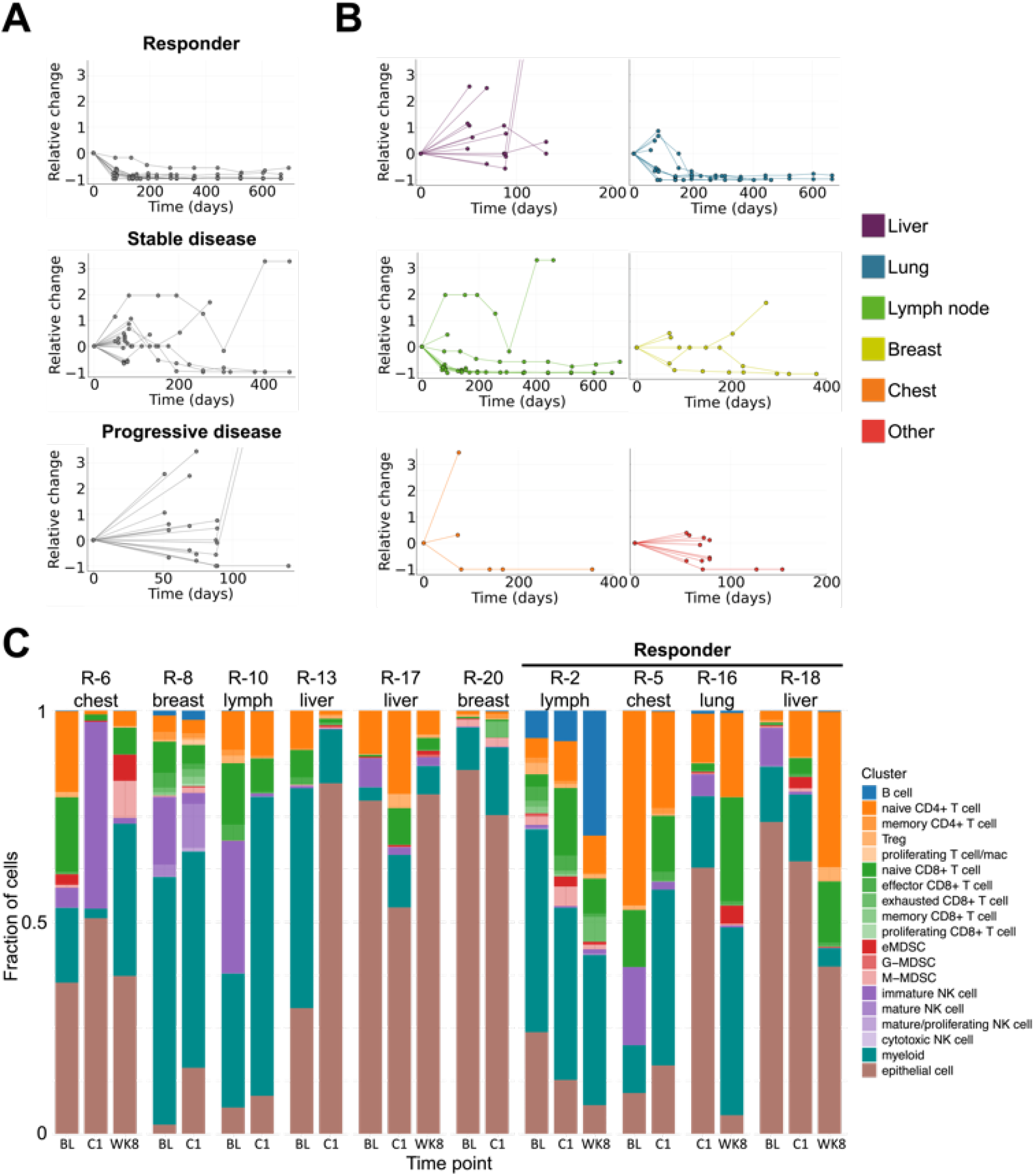
Target lesion tumor measurements and spatial proteomics detail response/non-response to combination treatment. **A.** Relative change in tumor volume for 55 individual tumors (from 20 patients ^11^) plotted by patient RECIST response classification. Five patients were classified as (partial/complete) responders; 15 patients were classified as non-responders (stable/progressive disease). **B**. Relative change in tumor volume for individual tumors plotted by site. **C**. Proportions of immune cell types characterized via spatial proteomics (imaging mass cytometry) in tumor tissues from 10 patients at baseline, C1D1 (post entinostat), and week eight (post combination therapy).

Patients are classified as a responder or non-responder based on RECIST: a standardized set of rules that measures changes in size from cross-sectional images of tumors over time to define response to treatment^33^. RECIST comprises four categories of response: complete response (CR), the disappearance of all tumors; partial response (PR), minimum of 30% reduction in tumor size; stable disease (SD), no significant change in tumor size; and progressive disease (PD), a 20% or greater increase in tumor size or the appearance of new lesions. These categories are determined by measuring target and non-target lesions and provide crucial endpoints in clinical trials for evaluating drug effectiveness. In the NCI-9844 cohort there were five responders (1 CR + 4 PR), nine patients with stable disease, and six patients with progressive disease. By immune-related RECIST (irRECIST), an updated set of criteria that takes into account the impact of immune infiltration on tumor size (pseudoprogression)^33^ there were five responders and eight patients with stable disease (two patients with SD by RECIST were classified as PD by irRECIST and one patient swapped from PD to SD).

To inform mathematical models of site-specific responses to therapy, we classified each individual tumor as “increasing” or “decreasing” based on its change relative to baseline (see Methods), as measured by RECIST (Fig. 1A-B). Of the 55 tumors followed, 32 were classified as decreasing and 23 as increasing. Patient tumor dynamics varied by site: multiple patients harboring tumors both that shrank and that grew over time. However, RECIST response does not take this variation into account: per RECIST, response is determined by the change in tumor volume of the combination of all measured tumors (target lesions). Comparison between the standard per-patient RECIST approach to quantifying response (via a sum of the largest diameters for each time point) and our lesion-specific approach based on relative change shows high concordance overall; our relative change approach has higher power to detect differences than the standard approach (see Supplementary Fig. S6 for comparison and Figs. for individual patient dynamics). One important note on the tumor dynamics is that there is a sampling bias towards responders, who typically stay in the trial for longer periods, resulting in a greater number of measurements per tumor over a longer duration than for non-responders (Fig. 1A). We observed strong associations for tumor site with response (Fig. 1B). Whereas most lung and lymph node tumors responded and shrank following therapy (Fig. 1B, blue and green), liver tumors exhibited poorer responses overall. This is in accordance with the literature, where it has been shown that ICIs are typically ineffective against liver metastases ^34,35,36,37,38^.

### Spatial proteomics reveals patient-specific signatures and tissue-specific immune composition shifts with response

Characterization of immune TME dynamic shifts with treatment provides informative correlative analyses, and are often available from clinical trials, especially those studying novel immunotherapies. Tools that enable integration of these dynamics with tumor response modeling could thus fill an unmet need and fuel new innovation for TME modeling. For 10 of the 20 patients followed (Table 1), we measured tumor-immune composition using spatial proteomics obtained through the analysis of patient biopsies via imaging mass cytometry (IMC)^10^. Using a custom panel of metal-isotope tagged antibodies for epithelial, stromal, and immune populations as well as functional markers (see Gonzalez et al. ^10^ and Methods) we quantified 30 immune cell subtypes across 10 major cell types. Of greatest relevance to this study among the immune compartment are the lymphocytes, mature myeloid cells, and MDSCs (Fig 1C). For each patient, multiple regions within the same site were analyzed at each of 2-3 time points: at baseline (BL), after entinostat treatment (C1D1), and after the addition of anti-PD-1 and anti-CTLA-4 (WK8).

A higher proportion of B cells was measured in responders, along with evidence of mature tertiary lymphoid structures and follicular structures in lymph node biopsies of responders ^10^. We also found (via co-occurrence analysis) that interactions between CD8+ T cells and macrophages decreased at week 8 compared to baseline only in responders ^10^. Analysis of the temporal tumor-immune compositions revealed overall stable or increasing proportions of epithelial cancer cells among non-responders, whereas responders exhibited decreasing epithelial cell proportions in 3/4 patients (Fig 1C). In several non-responders (4/6), the proportion of MDSCs increased over time; whereas in only one responder was an increase observed (Fig. 1C and Fig. S27 for the immune-only composition dynamics).

### Bayesian parameter inference captures distinct tumor-immune dynamics across patients and sites

In an effort to integrate the phenotypic characterization of tumor biopsies obtained by IMC with RECIST response measurements, we developed a pipeline for parameter inference in a patient- and site-specific manner with the goal of determining which tumor-immune parameters could explain (or drive) site-specific tumor responses (Fig. 2A). Our previous work ^18^ developed a model of tumor-immune dynamics, focusing on the role MDSCs during nascent metastatic growth without treatment. The previous model was developed to address they hypothesis that immune suppression by MDSCs affected de novo metastatic growth. Our new model addresses the hypothesis that the immune parameters differ by site of metastatic disease and as such influence response to treatment in a site-specific manner. To capture the TME dynamics observed in biopsies from patients in NCI-9844 we developed patient-specific models with initial conditions and parameters fitted from data. The delay in MDSC recruitment to the TME was set to zero since time has passed since each lesion was established (τ = 0), the cancer cell populations were initialized a the tumor steady state-representative of already established metastases, and tumors were simulated under a parameter regime that matched TME cell populations identifed via knowledge guided subclustering from scRNAseq of mouse models of primary breast and breast to lung metastatic TMEs ^10^.

**Figure 2:**
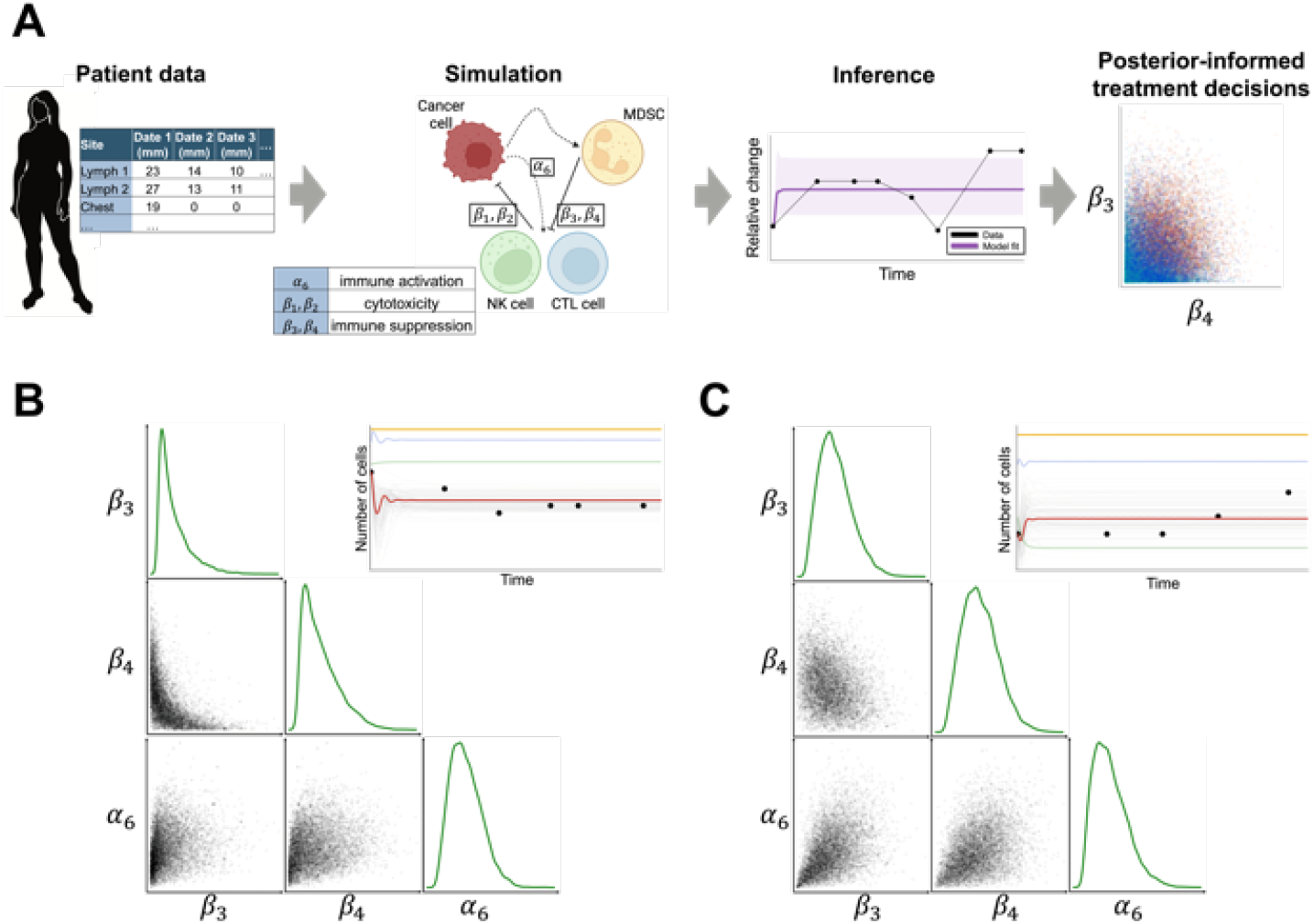
Bayesian inference pipeline to infer patient and tumor-specific immune responses to treatment. **A.** Outline of modeling and analysis pipeline. See Methods and Table 2 for further details. **B**. Example posterior parameter distributions and model fits for a tumor responsive to combination treatment.**C**. Example posterior parameter distributions and model fits for a tumor not responsive to combination treatment.

The model is defined below. *x*_T_, *x*_MDSC_, *x*_NK_, and *x*_CTL_ are the population sizes of cancer cells, myeloid-derived suppressor cells (MDSCs), natural killer (NK) cells, and CD8+ cytotoxic T cells (CTL) respectively, at time *t*. We use the general notation *δx*_*i*_ to denote the rate of change of *x*, thus:

**Table 2:** Description of model parameters and values.

| Notation | Description | Nascent tumor<sup>[18]</sup> | Steady state | Change |
| --- | --- | --- | --- | --- |
| $x_T(t), t \leq 0$ | initial condition for tumor cells | 1 or 2 | steady state | yes |
| $x_{\text{MDSC}}(0)$ | initial condition for MDSCs | $\alpha_2/\zeta_2$ | steady state | yes |
| $x_{\text{NK}}(0)$ | initial condition for NK cells | $\frac{\zeta_2\alpha_4}{\alpha_2\beta_3+\zeta_2\zeta_3}$ | steady state | yes |
| $x_{\text{CTL}}(0)$ | initial condition for CTL cells | 0 | steady state | yes |
| $\tau$ | delay parameter for MDSCs | varies | 0 | - |
| $\alpha_1$ | tumor growth rate | $10^{-1}$ | $10^{-1}$ | - |
| $\eta$ | tumor maximum size | $10^7$ | $10^7$ | - |
| $\beta_1$ | tumor cells inhibition rate by NK cells | $3.5 \times 10^{-6}$ | $6 \times 10^{-4}$ | yes |
| $\beta_2$ | tumor cells inhibition rate by CTL cells | $1.1 \times 10^{-7}$ | $7 \times 10^{-4}$ | yes |
| $\zeta_1$ | tumor cell death rate | 0 | 0 | - |
| $\alpha_2$ | MDSCs circulating rate | $10^2$ | $5 \times 10^2$ | yes |
| $\alpha_3$ | MDSCs expansion coefficient | $10^8$ | $10^8$ | - |
| $\zeta_2$ | MDSCs death rate | 0.2 | 0.2 | - |
| $\alpha_4$ | NK cells circulating rate | $1.4 \times 10^4$ | $5 \times 10^1$ | yes |
| $\alpha_5$ | NK cells expansion coefficient | $2.5 \times 10^{-2}$ | $2.5 \times 10^{-2}$ | - |
| $\beta_3$ | NK cells inhibition rate by MDSCs | $4 \times 10^{-5}$ | $4 \times 10^{-5}$ | - |
| $\zeta_3$ | NK cells death rate | $4.12 \times 10^{-2}$ | $4.12 \times 10^{-2}$ | - |
| $\alpha_6$ | CTL stimulation by tumor-NK cell interaction | $10^3$ | $4 \times 10^{-3}$ | yes |
| $\alpha_7$ | CTL expansion coefficient | $10^{-1}$ | $10^{-1}$ | - |
| $\beta_4$ | CTL inhibition rate by MDSCs | $10^{-4}$ | $10^{-4}$ | - |
| $\zeta_4$ | CTL death rate | $2 \times 10^{-2}$ | $2 \times 10^{-2}$ | - |
| $\gamma_1$ | steepness of MDSC production | $10^{10}$ | $10^{10}$ | - |
| $\gamma_2$ | steepness of NK production | $2.02 \times 10^7$ | $2.02 \times 10^7$ | - |
| $\gamma_3$ | steepness of CTL production | $2.02 \times 10^7$ | $2.02 \times 10^7$ | - |

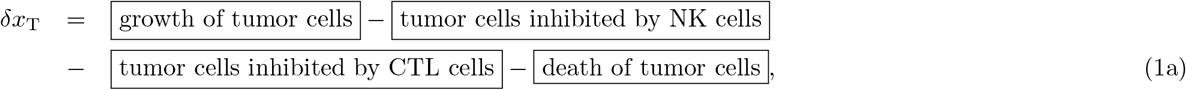

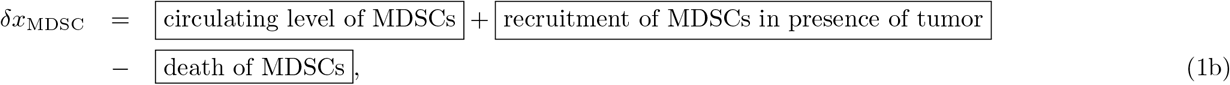

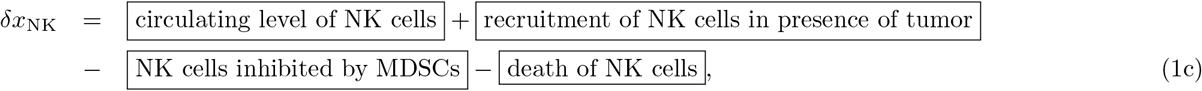

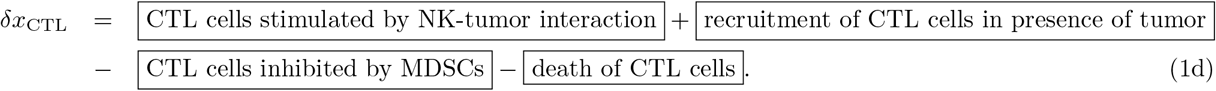

In different scenarios we consider the dynamics of an ordinary differential equation implementation of the model: 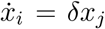, with *i, j* ∈ [T, MDSC, NK, CTL]; and a stochastic differential equation implementation:

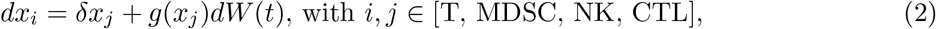

where *g*(*x*) denotes the noise model and *dW* (*t*) denotes an increment of a Weiner process (full details given in Methods). All parameters are defined along with values used in Table 2, and reflect processes specific to each cell type as guided by read outs of relative gene expression pathways from our own scRNAseq data and some from processes reported for each cell type in the literature.

Each individual lesion measured by RECIST was then fit to the model via Bayesian parameter inference (fits for all 55 tumors are given in SI Figs. S28-S82). To maximize information gained, we normalized all tumor size measurements and fit the model to the relative change in the tumor population size (see Methods), enabling direct comparison of data to the mathematical model. Parameter posterior distributions for a representative responding tumor and non-responding tumor are shown in Fig. 2B-C.

### Immune suppressive but not cytotoxic parameters control simulated responses to therapy

To fit individual lesions we selected a five-dimensional free parameter space consisting of (*β*_1_, *β*_2_, *β*_3_, *β*_4_, and *α*_6_) from the 24-parameter mathematical model (Table 2). Fig. 3A shows a table of these five parameters, each representing an important cell-cell interaction within the TME. This subset was selected based on each parameter’s sensitivity to fluctuation in tumor size ^18^ as well as the extent to which individual TME interactions are impacted by treatment. These latter effects were determined using our extensive preclinical data ^10^, in which TME cell subtypes were rigorously characterized by single-cell sequencing of cells isolated from lung metastases growing in mice treated with the same drug combination as used in patients from our clinical trial ^11^. Of the 39 cell subtypes present, interactions among the four most pertinent (tumor cells, MDSCs, NK cells, and CD8+ T-cells) were incorporated. Changes in cell signaling, interactions and infiltration inferred from these data guided the choice of the five-parameter subset analyzed here.

**Figure 3:**
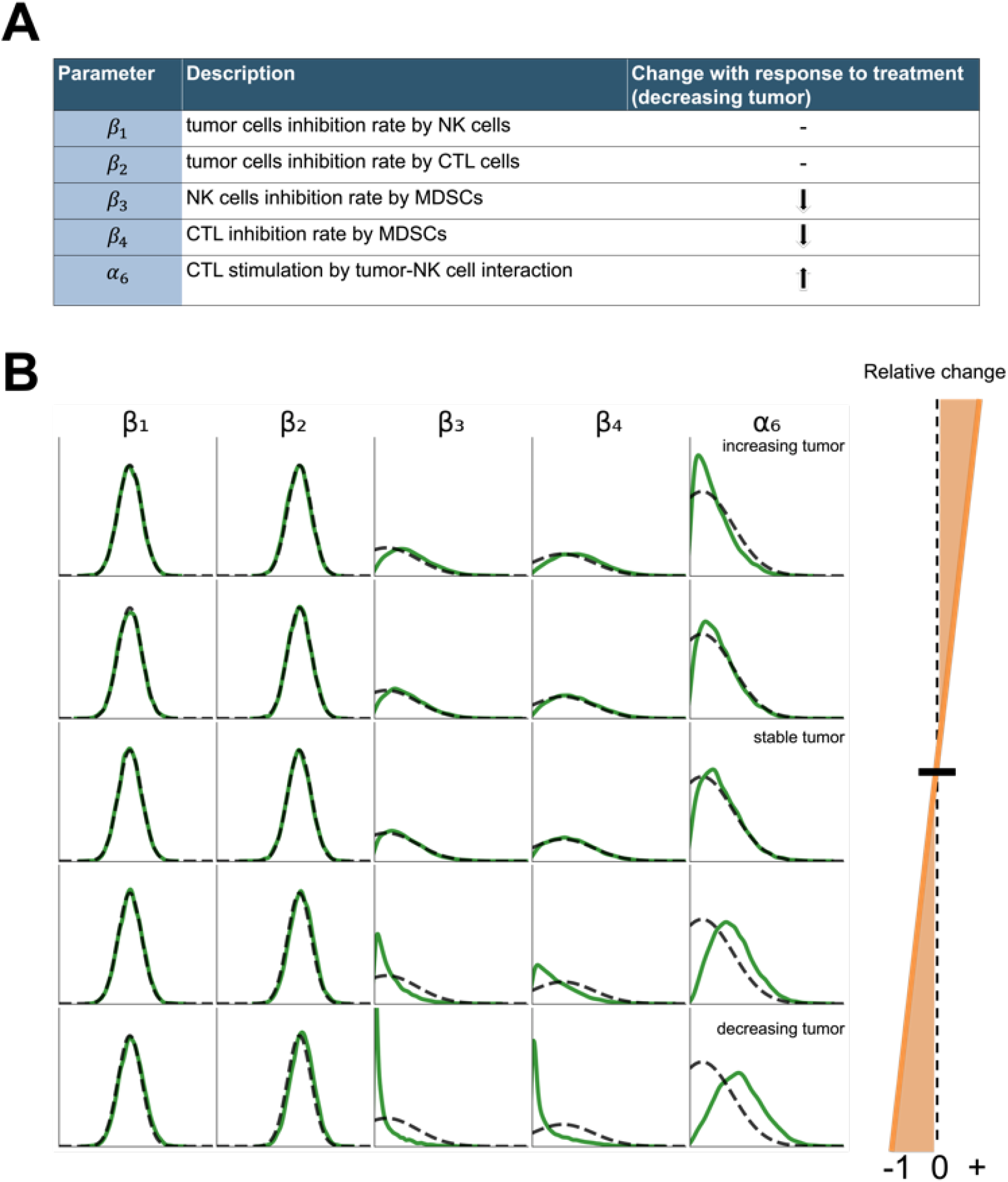
Immune suppression and CTL stimulation control tumor growth. **A.** Parameter table and notation. **B**. Posterior distributions (solid green) and prior distributions (dashed gray) are shown for five tumors. *β*_3_ and *β*_4_ decrease as tumor response improves, whereas *α* _6_ increases for responsive tumors. *β*_1_ and *β*_2_ show no change (posterior distribution compared to prior distribution) for both responsive and non-responsive tumors.

Analysis of all fitted tumors (of which five representative tumor model fits are shown; Fig. 3B) revealed that marginal posterior distributions (green lines) for two of the five parameters (*β*_1_, *β*_2_) did not deviate from the prior (dashed lines; Fig. 3B). These parameters — tumor cell killing by NK cells or CTLs — represent cytotoxicity thus revealing that these parameters are uninformative with regards to determining tumor response during combination treatment (they are “sloppy”^39^). As such, they are not included below in further fitting or analysis.

The parameters that characterized treatment-induced changes on immune cell populations that indirectly affected the tumor cells were the most informative of response. These three parameters are: *β*_3_ (NK cell inhibition by MDSCs), *β*_4_ (CTL inhibition by MDSC), and *α*_6_ (CTL stimulation by tumor-NK interaction). Each of these strongly influenced tumor response dynamics, as seen by the differences in posterior distributions compared to the priors. *β*_3_ and *β*_4_ decreased with tumors that shrank, consistent with the reduction in immune suppression necessary to mount an anti-tumor response. *α*_6_ increased with tumors that shrank, consistent with its definition as an activator of the anti-tumor response (thus positively correlated with anti-tumor response). We observed that the posterior parameters for models fitted to stable tumors differed little from their priors. This is expected given the initial conditions: tumors are simulated from their steady state.

To quantify the relative importance of the three parameters (*β*_3_, *β*_4_, *α*_6_) in their ability to predict tumor responses, we trained decision trees on the posterior distributions, using them to predict which combinations of parameters best determine response (i.e. classifies tumor as decreasing or increasing). We found that all three of the informative parameters (i.e. after excluding *β*_1_ and *β*_2_) were similarly predictive; they all contained information regarding tumor response (Supplementary Table S1). We also trained decision trees on posterior distributions to test their ability to predict the site of a tumor. Liver tumors generally respond poorly to treatment compared to lung or lymph node tumors (Fig. 1B), so we expect diminished immune-suppressive effects in the lung and the lymph node relative to the liver. For both liver vs. lung and liver vs. lymph node (Supplementary Tables S2 and S3) we found that MDSC suppression parameters (*β*_3_ and *β*_4_) were the most important in controlling response by site (higher predictive power in discriminating liver vs other tumors). Again, highlighting the importance of immune suppression in controlling differences in metastatic responses by site.

Overall, the three most important parameters (*β*_3_, *β*_4_, and *α*_6_) were those that describe differences in immune activation or suppression through tumor-immune interactions as they pertain to each specific site of disease. When we test the ability of the model to predict which parameters drive the responses in each site, we learn that it is the primarily the immune suppressive parameters that constrain responses to different degrees in the liver, lung, and lymph node.

### Virtual tumor simulation enables accurate patient-specific response predictions at new metastatic sites

One of the potential uses of this approach is to determine whether or not a specific treatment will control tumor growth at a site of disease not previously identified on a scan. This is of clinical importance because often times a change in therapy is initiated at the time of progression of known lesions and soon after the therapy is started a new area of metastasis is identified. If the new therapy was started less than 1-2 months prior, clinicians are often faced with determining whether the new lesion is likely to respond to the current therapy or not, i.e. wait another month before the current therapy is deemed a failure vs. change the therapy now. We set out to see if the model can predict response at ‘virtual’ sites of disease that are not yet reported in a patient. The implications of this would be that a clinician may be able to know with higher certainty whether a change in therapy is indicated now or if the current therapy may be effective in time.

Moreover, this approach can be used to overcome the intrinsic limitations of small and heterogenous clinical cohorts in trial, e.g. we may want to know how a new therapy affects metastases in the liver across the whole patient population even when only a proportion of the patients had known liver tumors. The ability to ask whether an investigational therapy can control disease at certain disease sites and not others without the need for every patient to have disease at all sites could boost the power of clinical trials without having to enroll more patients. To simulate virtual tumors, we sampled and simulated from the posterior distributions that captured site-specific dynamics of response (see Methods).

As a preliminary example, we simulated a virtual liver tumor in patient R-18 (who did not present with liver metastasis). This virtual liver tumor was simulated through parameter sampling based on site and patient response (Fig. 4A-C). We simulated tumor responses based on focal site (liver; purple), on the focal patient (R-18; green), or based on a combination of both, i.e. the complete virtual tumor simulation for liver metastasis in patient R-18 (Fig. 4D). Simulations for R-18 tumors are similar to a responsive tumor (see Fig. 2B); as expected since patient R-18 is a responder. Similarly, simulations for all liver tumors are similar to a non-responsive tumor (Fig. 2C); as expected since we know that most liver tumors in our cohort present as aggressive and non-responsive (Fig. 4A). The simulation of virtual liver tumors in patient R-18 showed non-progression (i.e. SD or PR) in approximately 81% of the tumors. This is a worse prognosis relative to the overall patient R-18 virtual population, but it remains a much better prognosis than for liver tumors more generally. This test case illustrates that if we were to later observe liver tumors in patient R-18, we would expect that approximately 81% of those tumors would shrink. This indicates that if patient R-18 had liver mets, tumors at this site would not respond as well to the treatment as compared to other sites of her metastatic disease, however, her overall prognosis may still reflect response. Should we have had this information during the trial, and this patient developed liver metastases we may decide to keep her on the trial and monitor closely as she would be predicted to respond even if she developed liver mets.

**Figure 4:**
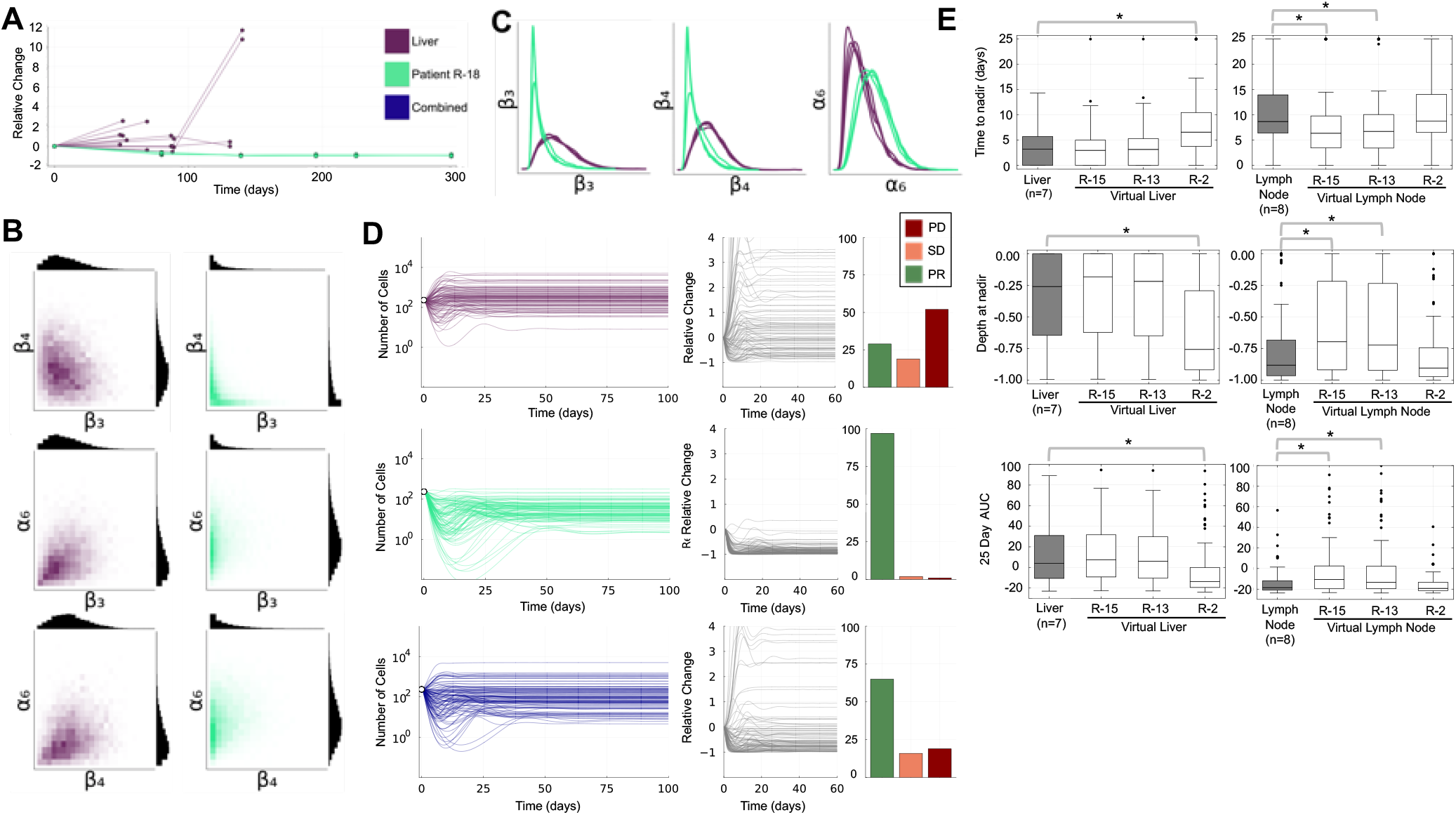
Virtual tumor populations to simulate patient-and site-specific responses. **A.** Tumor RECIST data from liver tumors from all patients (purple) and from tumors from patient R-18 (green). **B-C**. Joint and marginal posterior parameter distributions corresponding to models fitted to all liver tumors or to all tumors from patient R-18. **D**. Virtual tumor simulations (10^2^ for each sample) for the liver, for patient R-18, and for liver tumors in patient R-18 (blue). Bar plots (right) give the fraction of virtual tumor responses that respond by RECIST (PR; green), SD (orange), or PD (red). **E**. Summary statistic for the early tumor dynamics for virtual liver and lymph node tumors simulated for three patients and compared to all liver or all lymph node tumors (data). Early time AUC: area under the relative change tumor curve in the range [0,25] days. Distributions are computed from 10^2^ tumor simulations.

To assess the validity of our approach and test whether virtual tumor simulations agree with unseen data, we generated virtual liver and lymph node tumors for three additional patients from the cohort. As means of evaluation, we introduced summary statistics calculated from the early tumor dynamics to compare responses between patients and site. We quantified early tumor dynamics by three summary statistics of individual simulation trajectories: time to nadir, depth at nadir, and AUC at 25 days post-treatment (Fig. 4E). Patients R-2 and R-13 were selected as a canonical responder and non-responder, respectively (i.e. they serve as positive controls). R-15 was selected as the test case, since for this patient we have off-target lesion data which was not used at any point in the previous inference. Analysis of early tumor dynamics for R-2 confirms — in comparison to liver tumors — that significantly higher times-to-nadir and significantly lower depths at nadir and AUCs correspond to a clinical responder. Analysis of early tumor dynamics for R-13 confirms — in comparison to lymph node tumors — that significantly shorter times-to-nadir and significantly higher depths at nadir and AUCs corresponds to a clinical non-responder. R-15 matches the profile of R-13: differences in summary statistics indicative of progressive disease. This is confirmed by the off-target lesions followed for R-15. Multiple off-target lesions measured for R-15 were initially stable, but eventually grew and this patient progressed, in agreement with our prediction from virtual lymph node and liver tumor simulations.

These results highlight that statistics based on the early tumor dynamics can distinguish responses by site, and that virtual tumor simulations provide valuable insight into unseen sites of disease.

### Personalized response predictions based on predicted tumor-immune dynamics

In Gonzalez et al. ^10^, tumor responses to combination therapy required the simultaneous modulation of multiple TME interactions. Here, we investigated how by incorporating personalized measurements of immune parameters ^38^ we could improve simulated responses to virtual tumors. We incorporated immune-specific parameters into virtual tumors by identifying clinical changes in immune composition that could be translated into parameter changes in the model.

We observed changes in the CTL population in the IMC data (Fig. 1C). Three of the patients (R-18, R-2, R-6) had significant populations of effector CD8+ T cells, memory CD8+ T cells, and proliferating CD8+ cells (Fig. S27). For these patients, we simulated virtual tumors with a CTL stimulation modulation, for which we scaled CTL stimulation (*α*_6_) and set 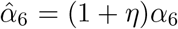, where *η* is the relative change in the proportion of CD8+ T cells at week 8 relative to baseline.

Analysis of responses in patients without (Fig. 5A) or with CTL stimulation (Fig. 5B) identified dramatic differences. When the proportion of CTLs increases in the TME (patient R-18, *η >* 0), virtual tumors were much more likely to respond to the treatment. Conversely, when the proportion of CTLs decreased during treatment (patients R-2 and R-6, *η <* 0), simulations the virtual tumors with the CTL modulation respond worse to treatment.

**Figure 5:**
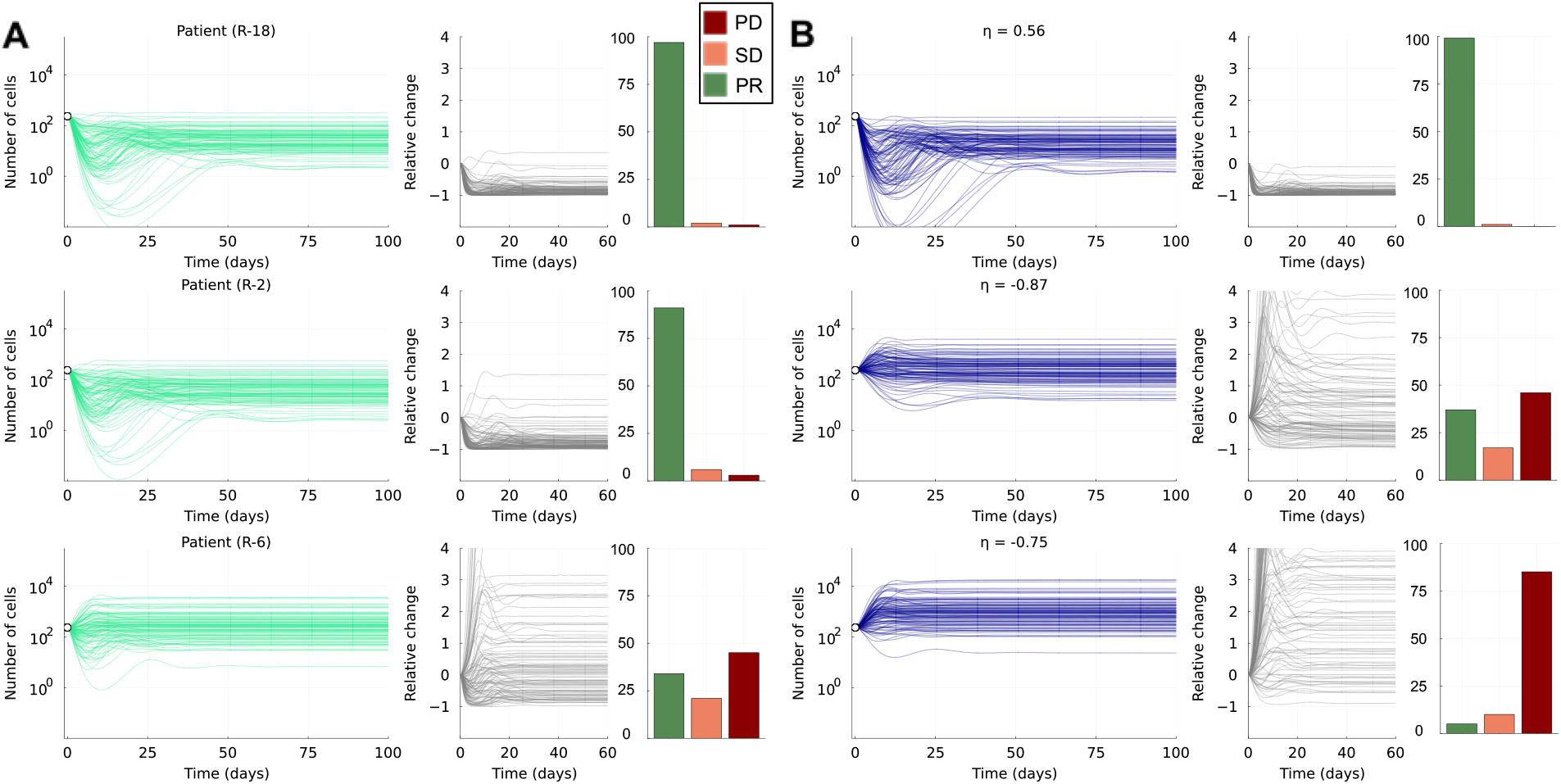
Simulation of patient-specific immune dynamics reinforces tumor responses to treatment. **A.** 10^2^ tumor simulations for patient R-18, patient R-2, and patient R-6. The fraction of tumors that are PR (green), SD (orange), and PD (red) are also shown. **B**. Same as A but tumor simulations include an *α*_6_ (CTL stimulation by tumor-NK cell interaction) modulation as quantified by IMC data.

### Immune parameters are identifiable even without measurement of the immune cell population dynamics

Temporal inference is ameliorated by denser sampling in time, however in a clinical setting this is often hard or impossible. Both costs and patient considerations limit the number of possible scans that can inform RECIST. Serial biopsies are hard to perform in all patients, but especially in those with metastatic disease in locations that are technically challenging to sample and in patients who are not feeling well thus less amenable to serial biopsies. Compounding this is the lack of temporal data for immune cell populations. As a result, parameters of the model (even after reducing the dimension of parameter space from five to three) are likely to be structurally and practically non-identifiable ^41^. To test this and assess if use of our model could be used to supplement the need for serial sampling via imaging or biopsies, which is still done in clinical trials despite listed limitations, we assessed parameter identifiability via a profile likelihood approach ^42,40^. The models considered in Simpson and Maclaren ^40^ are analytically solvable; in lieu of such solutions, here we adapt the likelihood function for use with numerical solutions of our four-state ODE system. This facilitates analysis of parameter identifiability both considering our clinical data and considering data generated *in silico* to quantify conditions under which parameters can become identifiable. In other words, what does our input have to be in terms of usage of patient data to be able to accurately identify parameters and make predictions.

In the case of tumors both from responders (Fig. 6A) and non-responders (Fig. S84), we found (as we expected) that the three parameters of the model were non-identifiable. In order to assess conditions under which parameters of the model become identifiable, we simulate virtual tumors with differing sampling densities (Methods). In the case where a tumor measurement was made every day for 300 days, all three model parameters are identifiable (Fig. 6B). I.e. small 95% asymptotic confidence intervals (blue) are observed for each parameter around its maximum likelihood estimate (MLE; green). While this sampling frequency would be infeasible at large scale, it is striking that all parameters are identifiable since the data contain no explicit measurements of immune population dynamics, only the tumor; yet the model parameters do not directly influence the tumor dynamics. The three model parameters (*β*_3_: NK cell inhibition by MDSCs; *β*_4_: CTL inhibition by MDSCs; and *α*_6_: the CTL stimulation rate due to tumor-NK cell interactions) are identifiable by fitting the four-population model to data characterizing only the tumor population. We also simulated more realistic scenarios where data is limited, and asked: what is the best strategy if a total of 11 measurements are possible over 300 days. With regularly spaced measurements every 30 days, all three parameters are non-identifiable (Fig. 6C). However, if measurements are made every day for the first week post-treatment and then evenly spaced from day 8 to 300, the identifiability improves dramatically: *α*_6_ becomes identifiable (Fig. 6D). This demonstrates that quantifying the tumor dynamics immediately after treatment is highly valuable and can yield constrained, identifiable parameters with modest increases in cost.

**Figure 6:**
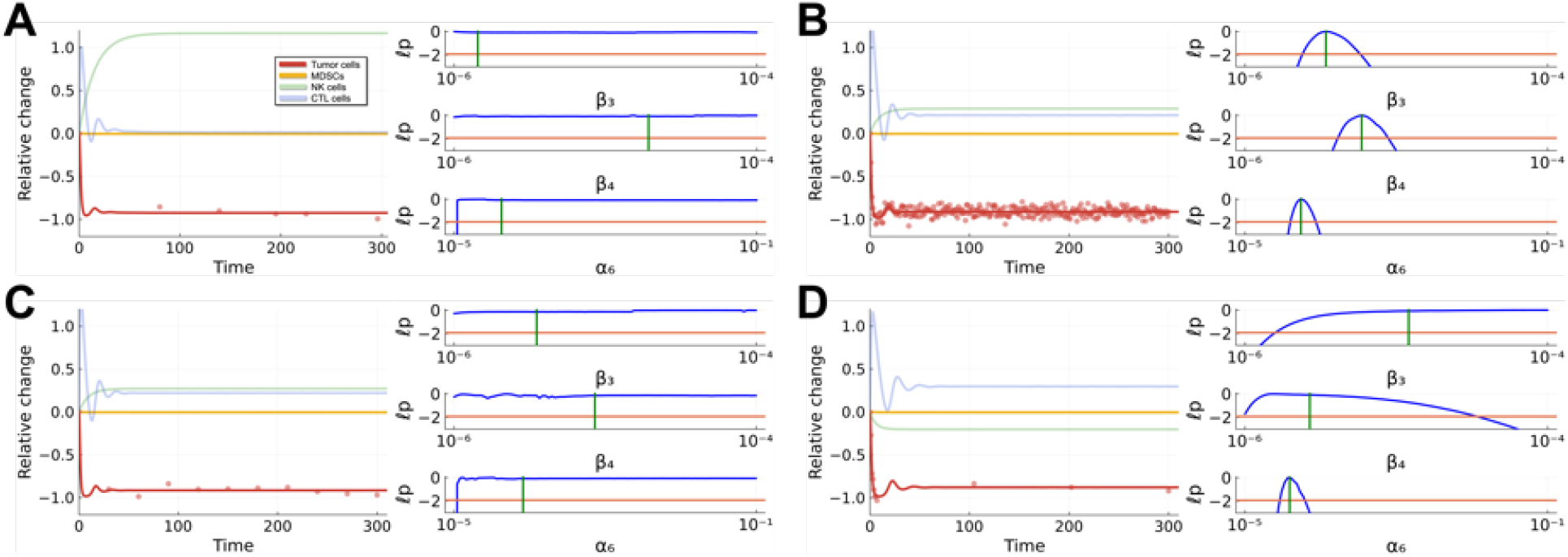
Parameter identifiability is constrained by temporal sampling. Left panels show tumor data (red dots) along with the mathematical model simulation with *β*_3_, *β*_4_, *α*_6_ at MLE values. Right panels show univariate profile likelihood functions for each of three parameters. The likelihood function (blue line) is superimposed with a vertical green line at the MLE and a horizontal red line at the asymptotic 95% threshold ^40^. **A**. Fits to actual tumor data from clinical trial (tumor 1). **B**. Fits to data simulated at a resolution of daily scans. All three parameters are identifiable and well-constrained. **C**. Fits to data simulated at 11 equally spaced points (every 30 days) over the one-year timespan. **D**. Fits to data simulated for daily scans for the first week after treatment initiation and three equally spaced scans subsequently; the total number of data points is equal to panel C.

Finally, we performed profile-wise analysis on simulated data from all four cell populations, including the immune cell states: myeloid-derived suppressor cells, natural killer cells, and CD8+ cytotoxic T cells. Fig. S85 shows the univariate likelihood profiles for a decreasing tumor and Fig. S86 shows the same for an increasing tumor under the same sampling assumptions as above (Fig. 6). Here, with all four populations measured, we observe that with uneven measurement dispersal of 11 measurements over 300 days, all of the parameters become identifiable.

## Discussion

We have presented an integrated mathematical modeling and Bayesian inference approach to fit models to clinical data. We demonstrated the utility of modeling to predict overall treatment outcomes and in doing so to measure how site of disease, individual patient dynamics, and tissue-based immune cell proportions influence response. Our *in silico* simulation pipeline links theory to data to accelerate biological discovery in complex TMEs ^43,10,26,13^. In developing this pipeline we have enabled the analysis of large virtual tumor & patient cohorts, helping to overcome the limitations of small clinical trials, and demonstrating the potential to further develop these approaches for use in treatment stratification.

We fit tumor data (RECIST) from individual patient lesions to a tumor-immune mathematical model to determine patient- and site-specific response dynamics. We find that TME interactions, rather than tumor inhibition or tumor cell-specific growth parameters, controlled tumor responses to combination therapy. Through posterior parameter sampling and simulation, we extrapolate beyond the data to predict response probability in virtual metastatic tumors at sites lacking data. This enables analysis of the responses of large virtual tumor cohorts to new treatments.

Via profile-wise identifiability approaches, we demonstrated that tumor data alone can identify important parameters representing interactions between TME populations. Parameter identifiability improves substantially with increased temporal resolution in tumor measurements immediately after treatment initiation, when TME dynamics change most rapidly. Our study therefore strongly suggests it would be beneficial to take more frequent measurements via patient scans in the weeks immediately following treatment, which can be offset by less frequent scans subsequently.

Overcoming patient- and organ-specific heterogeneity do remain sizeable challenges, exacerbated when by the small sample sizes that are inevitable in phase 1 clinical studies. Primarily for reasons of tractability and to facilitate inference, we do not currently consider spatial models, yet further analysis of the spatial proteomics data could inform spatial models of the TME dynamics during and after therapy. Recent studies incorporating spatial effects into models of tumor-immune dynamics have yielded new insights ^43,44,45^, though large-scale inference with such models remains a substantial hurdle.

The next steps for this model would be inclusion in a prospective study of patients with metastatic breast cancer who are receiving immunotherapy, specifically patients with triple negative breast cancer receiving pembrolizumab (anti-PD1) and chemotherapy which is currently standard of care for those whose tumors are expressing PD-L1. One use of the model prospectively will be to make predictions about which sites of disease are likely to progress vs respond and then follow patients over time to determine the accuracy of these predictions. Further validation will include serial biopsies of metastatic tumors at different sites of disease to determine if site-specific immune dynamics predictions can be validated. Subsequently, translational potential of this model could include informing the next line of therapy. For example, checkpoint inhibition is often not successful in treatment of liver metastasis thus, if a patient is predicted to progress within her liver metastases, checkpoint inhibition may not be the next best choice for therapy and a clinical trial could be considered for this patient. In comparison, if a patient is predicted to demonstrate progression in lymph nodes or lung metastases, immunotherapy may be the next best option for these patients.

Mathematical modeling has shown much promise for clinical translation in diagnostic fields like pathology and radiology but lacks wide-scale adoption in immune-oncology treatment decisions, in part understandably due to the complexity of these responses ^46,16,10^. We have shown how inference of patient-specific response dynamics and simulations *in silico* can boost the information we can gain from small trial cohorts, proving significantly more information than RECIST classifications. The approach has broad future utility to inform both clinical practice and drug development where trial response data are sparse.

## Methods

### Tumor data measured by RECIST

Tumor data is provided from the phase 1b cohort (NCI-9844: nivolumab + ipilimumab + entino-stat)^11^. There were 55 measurable target lesions (minimum two measurements) sampled from 20 patients in the trial, which are shown in Fig. 1A by response and Fig. 1B by site. Each tumor had a baseline assessment before the initiation of treatment in the clinical trial (for the purposes of fitting we set the time of the baseline assessment to be zero). Tumor size was then reassessed multiple times over the course of treatment. At each assessment, tumor size was measured in millimeters (mm) in one dimension (*x*), which we converted to a volume following the convention adopted by Laleh et al.^21,47^, i.e. estimating the volume in mm^3^ 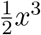. We estimated the absolute number of tumor cells from this volume ^48^ by multiplying by a factor of 10^7^. We considered the relative change in the tumor population, which was measured as the difference between the measurement and the baseline assessment, divided by the baseline assessment (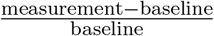, which gives a real number ∈ [−1, ∞)). As the relative change at the baseline assessment was always zero, we remove this data point for fitting for all tumors. Since only the tumor data is available, we fitted the log transformed data from this population (i.e. log(*x*_T_ + 1)). All of the data for each of the 55 tumors is available in a supplementary file (**data.xlsx**) available on GitHub. Further statistics are also available in the Supplementary Information.

### Spatial proteomics via imaging mass cytometry

Full details the imaging mass cytometry (IMC) data collection, processing, and analysis are described in Gonzalez et al. ^10^. Briefly, formalin-fixed, paraffin-embedded patient tumor biopsies from NCI-9844 were analyzed via IMC using a custom panel of 34 metal-isotope tagged antibodies. Raw IMC images were pre-processed using the IMmuneCite-IMClean pipeline in Python (version 3.12.10). Samples were processed through the Bodenmiller Steinbock pipeline and Mesmer (Deep-Cell) was used to perform cell segmentation. A table showing the proportion of all immune cell types in the IMC data is included in a supplementary file (**IMC proportions.xlsx**) available on GitHub.

Due to the combination of hormone positive and triple negative breast tumors included in the study and the tumor heterogeneity present between patients we were unable to identify a single marker exclusive to cancer cells in the IMC data. Instead, we identified epithelial cells using the joint expression of CK and E-cadherin. When these epithelial cells were within regions of interest (ROIs) that were identified by study pathologist Dr. Ashley Cimino-Mathews (Johns Hopkins University) as tumor, they were labeled as tumor epithelial cells. There are limitations to this assumption as it is possible that normal epithelial cells could be contained within a tumor bed, however the large majority of epithelial cells in these ROIs are malignant tumor cells and are thus all considered as such.

### Mathematical modeling of tumor-immune dynamics

We model the tumor-immune dynamics of four cell populations described by a system coupled nonlinear ordinary differential equations: the cancer cells, effector CD8 T cells, natural killer (NK) cells, and myeloid-derived suppressor cells (MDSCs). Where previously we considered a stochastic delay differential equation model of nascent tumor growth ^18^, here we model established tumors for which the delay in recruitment is no longer relevant and the role of noise is diminished (large initial tumor volumes).

In order to capture the dynamics of tumors undergoing treatment by entinostat and ICIs, we modified our previously published stochastic delay differential equation model according to the specificities of this microenvironment. We considered a deterministic differential equation model with the same four cell populations, with the following main changes: there is no MDSC recruitment delay to the tumor microenvironment (*τ* = 0), we initialized cell populations at tumor-steady state (Table 2), and we implemented a parameter regime change to match model population sizes to cellular proportions observed in the vehicle condition of the single-cell data in murine metastasis ^10^. We denoted *x*_T_, *x*_MDSC_, *x*_NK_, and *x*_CTL_ as the populations of tumor cells, MDSCs, NK cells, and CTL cells, respectively, at time *t*. We focused on the most important interactions between tumor, immune, and MDSC populations, which allowed us to gain insight into system dynamics and tumor growth. The model can be expressed conceptually (i.e. agnostic as yet to the form of the dynamics) in Eqns. (1a)-(1d), where *δx*_*i*_ denotes the rate of change of *x*_*i*_, *i* ∈ [T, MDSC, NK, CTL]. Based on these biological processes, we developed a model to characterize tumor-immune interactions that takes the form:

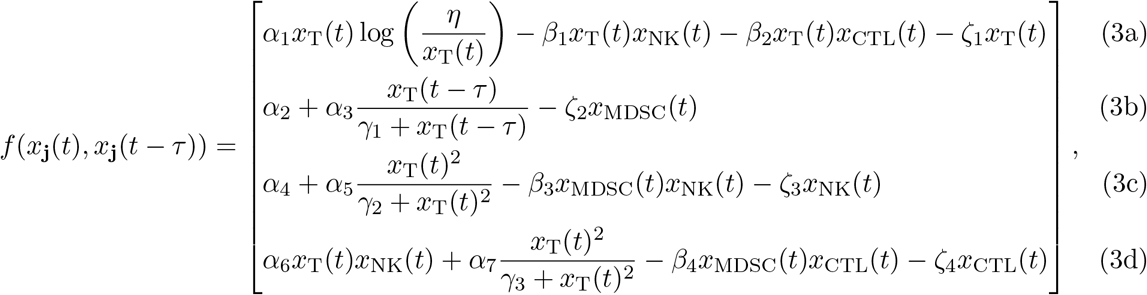

with description of the parameters is given in Table 2. A full derivation of each term in the model is given in Kreger et al. ^18^.

In simulations of established tumors undergoing treatment (Eqns. (3a)-(3d)), the initial conditions were set by the tumor steady state, which is numerically determined. Unless explicitly stated otherwise, all parameter values used for simulation are as defined in Table 2. All models were developed in the Julia programming language ^49^, using DifferentialEquations.jl^50^.

### Bayesian parameter inference

RECIST criteria have been developed for use in clinical trials to determine the change in tumor burden of selected target lesions to inform whether a patient is responding to a given therapy ^30,31^. We implemented Bayesian parameter inference to fit the model to tumor responses to classify tumor sizes and responses over time. We fitted differential equation-based models to data following a similar conceptual framework to ^18,21^. For this we employed Bayesian parameter inference, implemented in Turing.jl^51^.

For inference, a five-dimensional parameter space was considered, in which we fitted the following parameters (Fig. 3A): *β*_1_ (tumor cells inhibition rate by NK cells), *β*_2_ (tumor cells inhibition rate by CTL cells), *β*_3_ (NK cells inhibition rate by MDSCs), *β*_4_ (CTL cells inhibition rate by MDSCs), and *α*_6_ (CTL stimulation by tumor-NK cell interaction). We did not infer all parameters simultaneously due to the high dimensionality of the parameter space; we chose parameters to infer based on their importance as judged by prior knowledge and global sensitivity analysis ^18^. The parameters chosen affect all populations of the model.

A prior distribution over the five-dimensional parameter space was selected using weakly informative Normally distributed priors, with means set to the values given in Table 2, and relatively large variances:

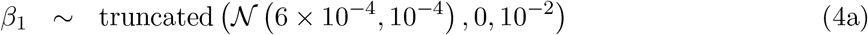

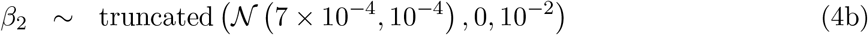

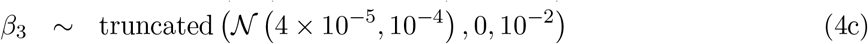

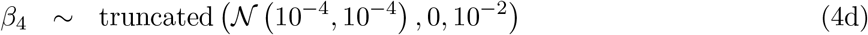

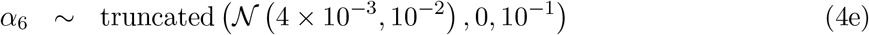

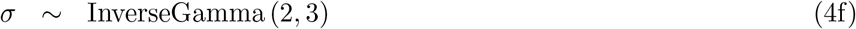

where *σ* is the observational noise. For each tumor we ran four independent Markov chain Monte Carlo (MCMC) simulations with 2 × 10^3^ iterations using the No U-Turn Sampler (NUTS) with a target acceptance ratio of 0.65^52^.

### Decision trees

We used a decision tree classifier to test whether different combinations of the marginal posterior parameters obtained from Bayesian parameter inference can classify tumor responses as 1) either decreasing or increasing over time or 2) based on site of tumor. Decision trees were built using DecisionTree.jl^53^ and cross validation was performed using scikit-learn ^54^.

### Virtual tumor simulation

To simulate virtual tumors based on site or patient, we use the posterior parameter values (e.g. liver tumors, patient R-18 tumors) to fit a Gamma posterior distribution for *β*_3_, *β*_4_, and *α*_6_ using the *fit()* function in the Distributions.jl package ^55^. Simulations of the mathematical model used values of *β*_3_, *β*_4_, and *α*_6_ randomly sampled from the relevant analytical posterior distribution. To simulate tumors based on a combination of site and patient (e.g. liver tumor in patient R-18) we sampled 10^6^ values of each parameter for the site and 10^6^ values of each parameter for the patient. We then combined these (for a total of 2 × 10^6^ samples) and fitted a Gamma distribution which represents the combined posterior distribution. We did not combine the site and patient posterior parameter values because the number of tumors varied; we assumed equal influence on the virtual tumor of the site and the patient in which it present.

For simulations with CTL stimulation modulation, we set the modified value of *α*_6_ to be 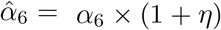, where *η* is the relative change in the combined proportion of effector CD8+ T cells, memory CD8+ T cells, and proliferating CD8+ T cells at week 8 compared to baseline.

### Profile-Wise Analysis

To quantify the identifiability of each fitted model parameter we implement a profile-wise parameter analysis approach ^40,42^. We adapt the publicly available workflow in Simpson and Maclaren ^40^, written in the Julia Programming language (https://github.com/ProfMJSimpson/NonidentifiableWorkflow), to our model and data. This results in two main modifications to their code, which are:

1. The models used in Simpson and Maclaren ^40^ (logistic growth model and Richards’ model) are analytically solvable single ODEs. The analytical solution of their model is an input to the loglikelihood function. Our model is not analytically solvable, and therefore we change the input to the loglikelihood function to the numerical solution of our model using the *Tsit5()* numerical ODE solution algorithm.
2. As the models used in Simpson and Maclaren ^40^ were single ODEs, the temporal data is one-dimensional as well. For our model, we consider scenarios in which we i) have only tumor population data, and ii) have four-dimensional data from all of the four model populations.

In order to generate simulated relative change data (either for one model population or for all model populations) we pick a representative decreasing tumor (tumor 1) and increasing tumor (10). We set *β*_3_, *β*_4_, *α*_6_ to be the median values from the Bayesian fitting of those respective tumors. We then add noise at every measured point taken from a normal distribution centered around 0 with a standard deviation of 0.05. The simulated data is either measured at constant intervals throughout the timespan (Fig. 6B,C) or more densely after the initiation of treatment and then at regularly spaced intervals after that (Fig. 6D).

## Author statements

### Software and Data Availability

All code and associated datasets generated in this study are available on GitHub at: https://github.com/maclean-lab/beyond_RECIST.

## Acknowledgements

We would like to thank Alexander Browning and members of the MacLean and Roussos Torres labs for helpful discussions. We also thank all of the patients (and their families) who participated in NCI-9844. We would like to acknowledge the Site-PIs from NCI-9844 Adam Brufsky, Patricia LoRusso, Vincent Chung, Yuan Yuan. We would also like to acknowledge others who contributed to ensuring completion of the trial and provided advice including Vered Sterns, Roisin Connolly, Richard Piekarz, Howard Streicher, and Elizabeth M. Jaffee, as well as Ashley O’Connor, who contributed to data management.

## Author Contributions

**Conceptualization**: JK, EG, ETRT, ALM. **Methodology**: JK, ALM. **Software**: JK, XW, JAM, ALM. **Investigation**: JK, EG, XW, JAM, ETRT, ALM. **Formal analysis**: JK, ALM. **Writing - original draft**: JK, ETRT, ALM. **Writing - reviewing and editing**: JK, EG, XW, JAM, ETRT, ALM. **Funding acquisition**: ETRT, ALM. **Supervision**: ETRT, ALM.

