## Supplementary Tables and Figures for "Beyond RECIST: mathematical modeling and Bayesian inference reveal the importance of immune suppressive parameters in metastatic breast cancer"

---

#### Supplementary Tables

S1 **Classification of tumor growth responses using posterior parameter values for increasing vs decreasing tumors** . . . . 4
S2 **Classification of tumor growth responses using posterior parameter values for liver vs lung tumors** . . . . 4
S3 **Classification of tumor growth responses using posterior parameter values for liver vs lymph node tumors** . . . . 4

#### Supplementary Figures

S1 **Number of tumors per patient** . . . . 5
S2 **Data points per tumor** . . . . 6
S3 **Tumor locations** . . . . 6
S4 **All data** . . . . 6
S5 **Other sites** . . . . 7
S6 **Relative change comparison** . . . . 7
S7 **Patient R-18** . . . . 8
S8 **Patient R-13** . . . . 8
S9 **Patient R-10** . . . . 8
S10 **Patient R-12** . . . . 9
S11 **Patient R-19** . . . . 9
S12 **Patient R-20** . . . . 9
S13 **Patient R-23** . . . . 10
S14 **Patient R-4** . . . . 10
S15 **Patient R-24** . . . . 10
S16 **Patient R-9** . . . . 11
S17 **Patient R-5** . . . . 11
S18 **Patient R-1** . . . . 11
S19 **Patient R-2** . . . . 12
S20 **Patient R-14** . . . . 12
S21 **Patient R-6** . . . . 12
S22 **Patient R-8** . . . . 13
S23 **Patient R-17** . . . . 13
S24 **Patient R-16** . . . . 13
S25 **Patient R-7** . . . . 14

S26 **Patient R-15** . . . . . 14
S27 **IMC subclusters** . . . . . 14
S28 **Tumor 1** . . . . . 15
S29 **Tumor 2** . . . . . 15
S30 **Tumor 3** . . . . . 15
S31 **Tumor 4** . . . . . 16
S32 **Tumor 5** . . . . . 16
S33 **Tumor 6** . . . . . 16
S34 **Tumor 7** . . . . . 17
S35 **Tumor 8** . . . . . 17
S36 **Tumor 9** . . . . . 17
S37 **Tumor 10** . . . . . 18
S38 **Tumor 11** . . . . . 18
S39 **Tumor 12** . . . . . 18
S40 **Tumor 13** . . . . . 19
S41 **Tumor 14** . . . . . 19
S42 **Tumor 15** . . . . . 19
S43 **Tumor 16** . . . . . 20
S44 **Tumor 17** . . . . . 20
S45 **Tumor 18** . . . . . 20
S46 **Tumor 19** . . . . . 21
S47 **Tumor 20** . . . . . 21
S48 **Tumor 21** . . . . . 21
S49 **Tumor 22** . . . . . 22
S50 **Tumor 23** . . . . . 22
S51 **Tumor 24** . . . . . 22
S52 **Tumor 25** . . . . . 23
S53 **Tumor 26** . . . . . 23
S54 **Tumor 27** . . . . . 23
S55 **Tumor 28** . . . . . 24
S56 **Tumor 29** . . . . . 24
S57 **Tumor 30** . . . . . 24
S58 **Tumor 31** . . . . . 25
S59 **Tumor 32** . . . . . 25
S60 **Tumor 33** . . . . . 25
S61 **Tumor 34** . . . . . 26
S62 **Tumor 35** . . . . . 26
S63 **Tumor 36** . . . . . 26
S64 **Tumor 37** . . . . . 27
S65 **Tumor 38** . . . . . 27
S66 **Tumor 39** . . . . . 27
S67 **Tumor 40** . . . . . 28
S68 **Tumor 41** . . . . . 28
S69 **Tumor 42** . . . . . 28
S70 **Tumor 43** . . . . . 29
S71 **Tumor 44** . . . . . 29
S72 **Tumor 45** . . . . . 29
S73 **Tumor 46** . . . . . 30
S74 **Tumor 47** . . . . . 30
S75 **Tumor 48** . . . . . 30
S76 **Tumor 49** . . . . . 31

S77 **Tumor 50** . . . . . . . . . . 31
S78 **Tumor 51** . . . . . . . . . . 31
S79 **Tumor 52** . . . . . . . . . . 32
S80 **Tumor 53** . . . . . . . . . . 32
S81 **Tumor 54** . . . . . . . . . . 32
S82 **Tumor 55** . . . . . . . . . . 33
S83 **Posterior distributions for simulations** . . . . . . . . . . 34
S84 **Parameter identifiability is constrained by the temporal density of tumor data for increasing tumors** . . . . . . . . . . 35
S85 **Parameter identifiability for decreasing tumors improves with data from all four model populations** . . . . . . . . . . 36
S86 **Parameter identifiability for increasing tumors improves with data from all four model populations** . . . . . . . . . . 37

### Supplementary Tables

| Features for prediction | Three-fold cross validation scores | Accuracy range (min, max) |
|---|---|---|
| $\beta_3$ | $65.7 \pm 8.68$ | (55.8, 72.1) |
| $\beta_4$ | $65.0 \pm 7.46$ | (56.5, 70.5) |
| $\alpha_6$ | $64.7 \pm 3.71$ | (60.6, 67.7) |
| $\beta_3, \beta_4$ | $69.0 \pm 11.01$ | (56.2, 75.7) |
| $\beta_3, \alpha_6$ | $66.9 \pm 9.95$ | (55.6, 74.1) |
| $\beta_4, \alpha_6$ | $66.3 \pm 10.07$ | (54.8, 73.4) |
| $\beta_3, \beta_4, \alpha_6$ | $68.8 \pm 13.72$ | (53.1, 78.4) |

Table S1: **Classification of tumor growth responses using posterior parameter values for increasing vs decreasing tumors.** Decision trees were used to classify tumor responses as either decreasing or increasing based on sets of posterior parameters as features. Three-fold cross validation scores are given as mean ± standard deviation. There are 23 increasing tumors and 32 decreasing tumors.

| Features for prediction | Three-fold cross validation scores | Accuracy range (min, max) |
|---|---|---|
| $\beta_3$ | $66.9 \pm 3.19$ | (63.4, 69.6) |
| $\beta_4$ | $65.4 \pm 3.47$ | (61.6, 68.5) |
| $\alpha_6$ | $62.7 \pm 2.75$ | (59.9, 6.4) |
| $\beta_3, \beta_4$ | $71.8 \pm 3.05$ | (68.6, 74.7) |
| $\beta_3, \alpha_6$ | $70.6 \pm 3.82$ | (66.4, 73.9) |
| $\beta_4, \alpha_6$ | $69.1 \pm 3.87$ | (65.0, 72.7) |
| $\beta_3, \beta_4, \alpha_6$ | $74.3 \pm 3.25$ | (70.9, 77.4) |

Table S2: **Classification of tumor growth responses using posterior parameter values for liver vs lung tumors.** Decision trees were used to classify tumor responses as either liver vs lung tumors based on sets of posterior parameters as features. Three-fold cross validation scores are given as mean ± standard deviation. There are 13 liver tumors and 12 lung tumors.

| Features for prediction | Three-fold cross validation scores | Accuracy range (min, max) |
|---|---|---|
| $\beta_3$ | $67.7 \pm 2.83$ | (65.1, 70.7) |
| $\beta_4$ | $66.8 \pm 3.57$ | (63.2, 70.3) |
| $\alpha_6$ | $62.3 \pm 1.93$ | (60.4, 64.3) |
| $\beta_3, \beta_4$ | $72.6 \pm 2.69$ | (69.7, 75.0) |
| $\beta_3, \alpha_6$ | $71.5 \pm 3.02$ | (68.6, 74.6) |
| $\beta_4, \alpha_6$ | $70.7 \pm 3.86$ | (66.8, 74.5) |
| $\beta_3, \beta_4, \alpha_6$ | $74.3 \pm 3.08$ | (70.7, 76.2) |

Table S3: **Classification of tumor growth responses using posterior parameter values for liver vs lymph node tumors.** Decision trees were used to classify tumor responses as either liver vs lymph node tumors based on sets of posterior parameters as features. Three-fold cross validation scores are given as mean ± standard deviation. There are 13 liver tumors and 12 lymph node tumors.

### Supplementary Figures

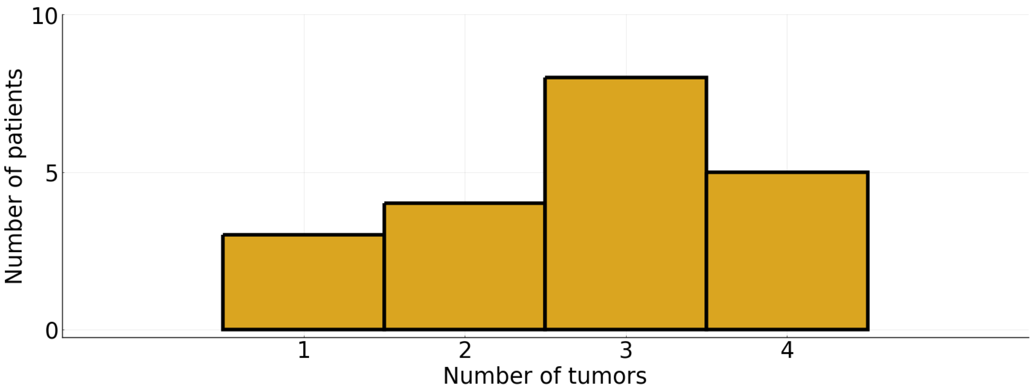


Figure S1: **Number of tumors per patient.** Histogram of the number of tumors per patient.

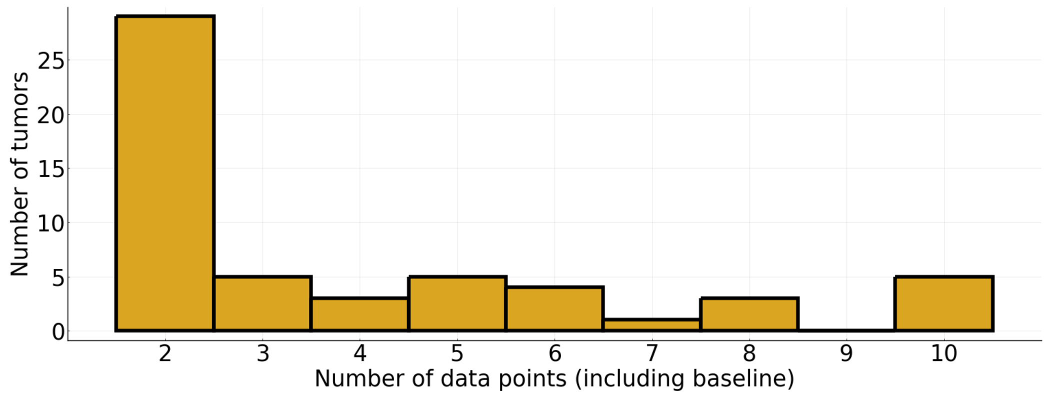


Figure S2: **Data points per tumor.** Histogram of the number of data points per tumor.

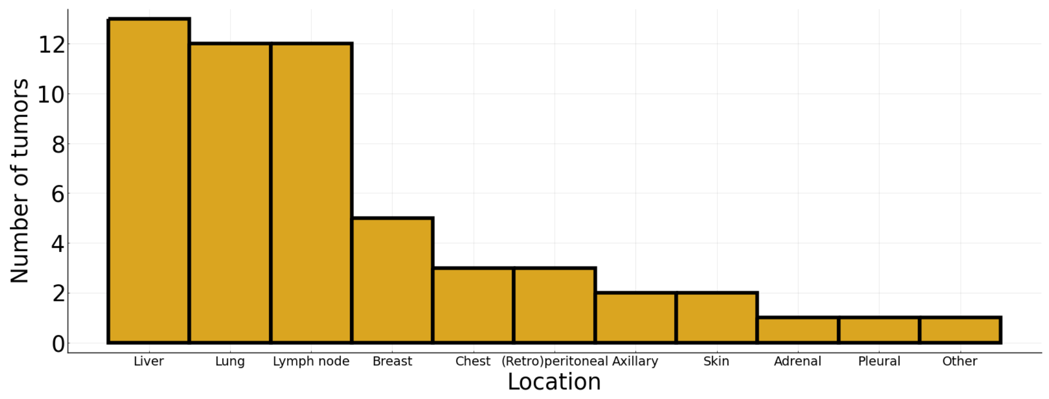


Figure S3: **Tumor locations.** Histogram of tumor locations.

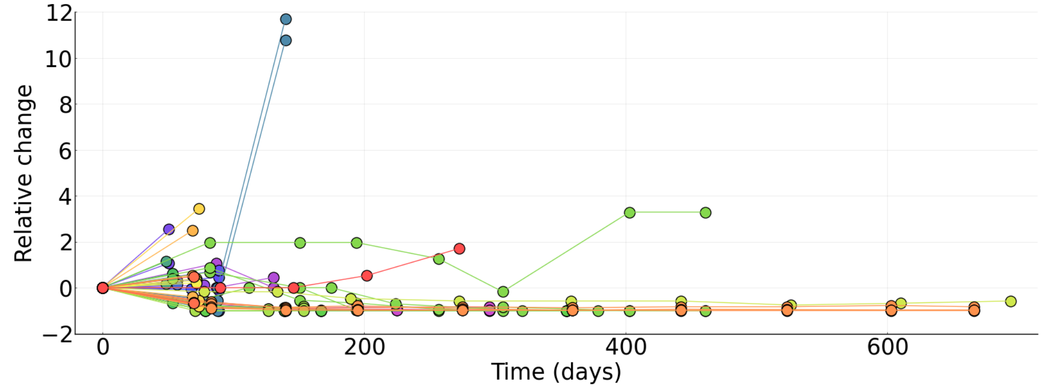


Figure S4: **All data.** Different colors represent different patients.

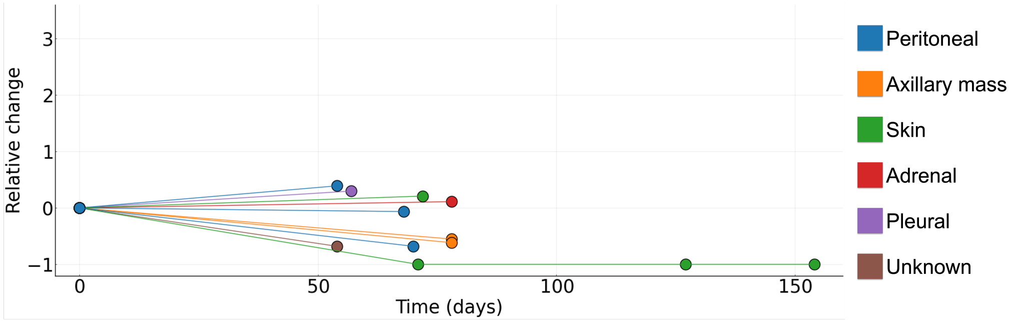


Figure S5: **Other sites.** All tumors not in liver, lung, lymph node, breast, or chest (see main text Fig. 1).

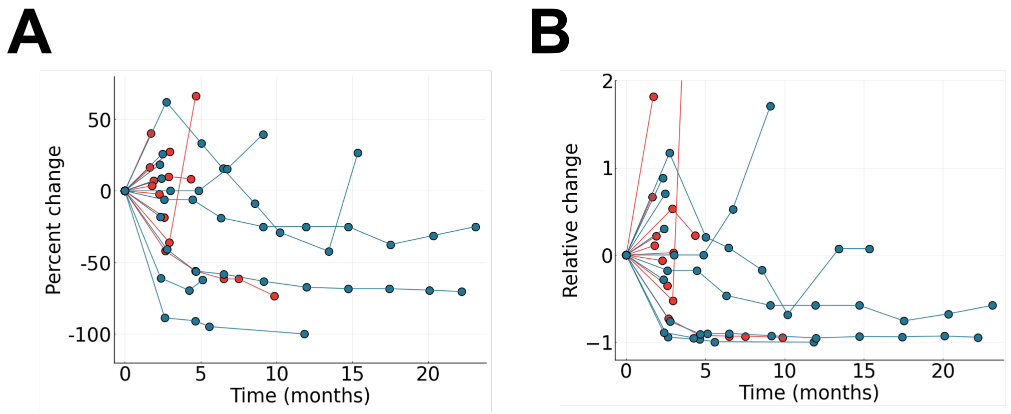


Figure S6: **Relative change comparison.** Comparing the relative change in sum of tumor diameters per patient vs change in tumor volume per patient.

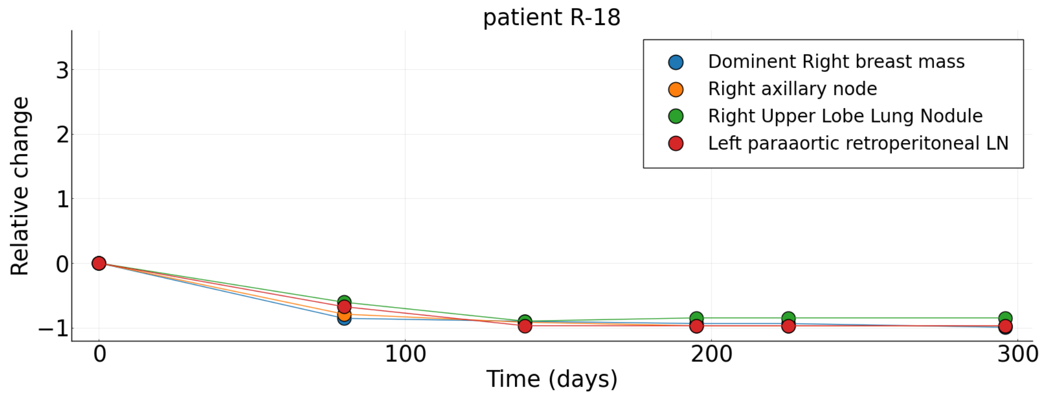


Figure S7: **Patient R-18. See main text Table 1 for details.**

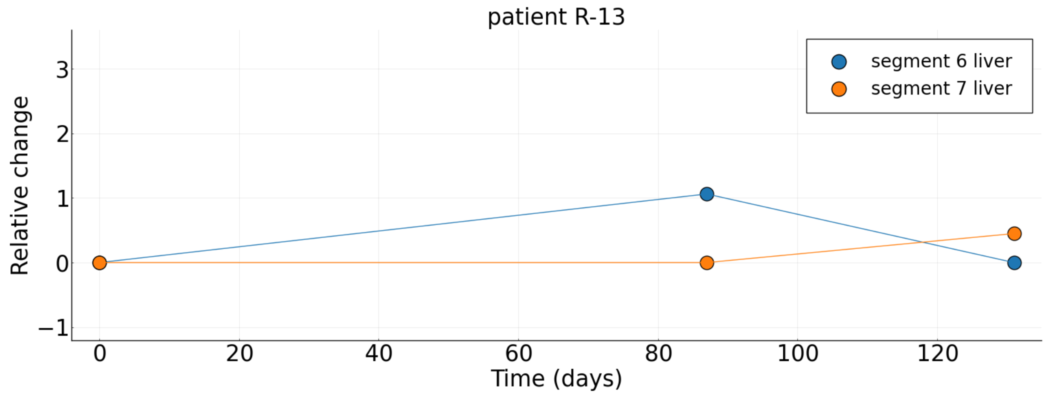


Figure S8: **Patient R-13. See main text Table 1 for details.**

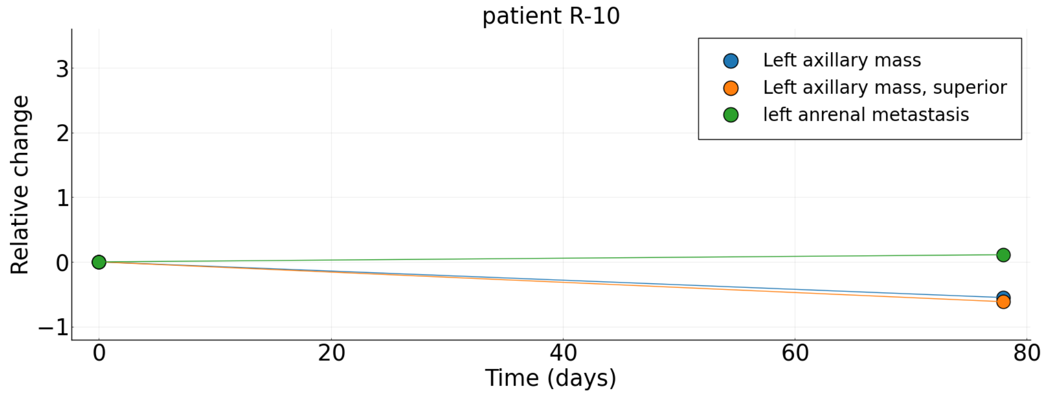


Figure S9: **Patient R-10. See main text Table 1 for details.**

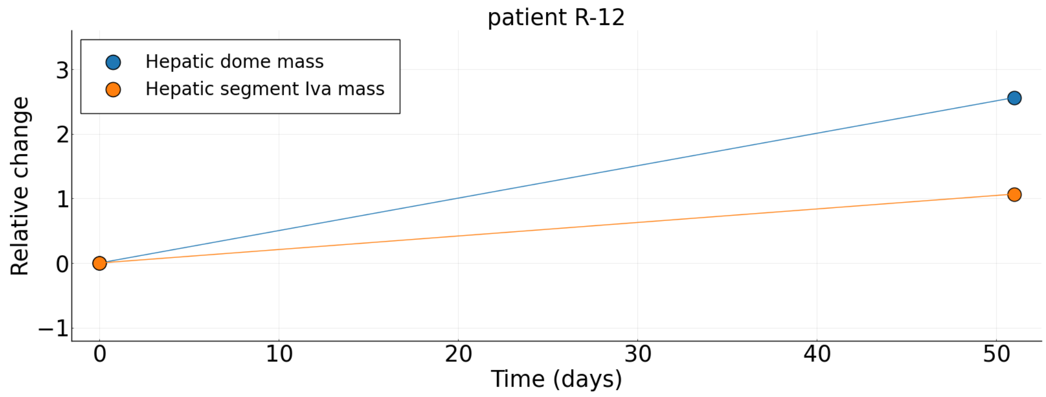


Figure S10: **Patient R-12. See main text Table 1 for details.**

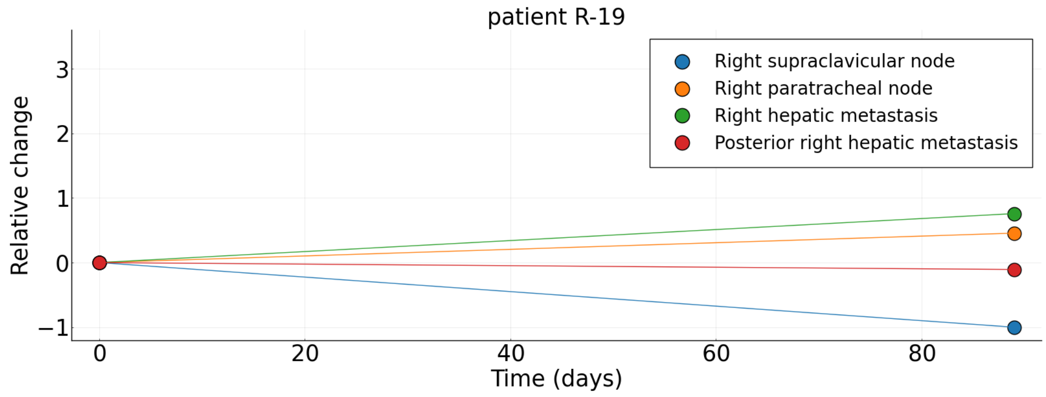


Figure S11: **Patient R-19. See main text Table 1 for details.**

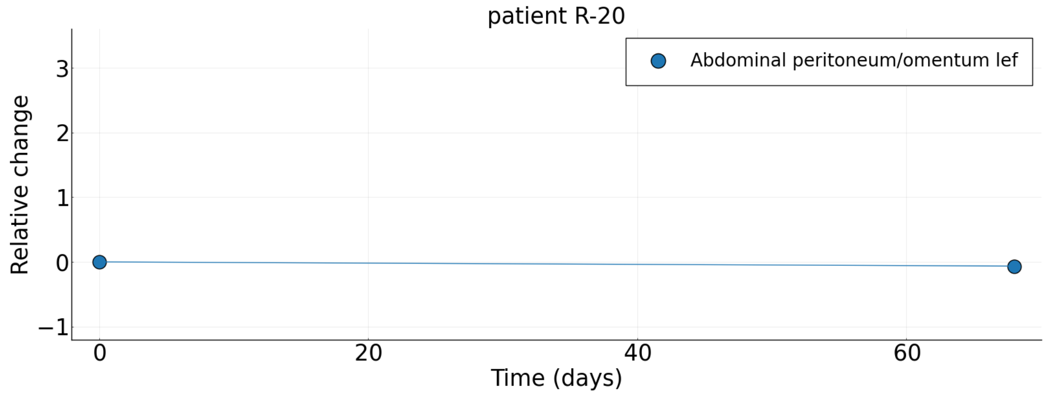


Figure S12: **Patient R-20. See main text Table 1 for details.**

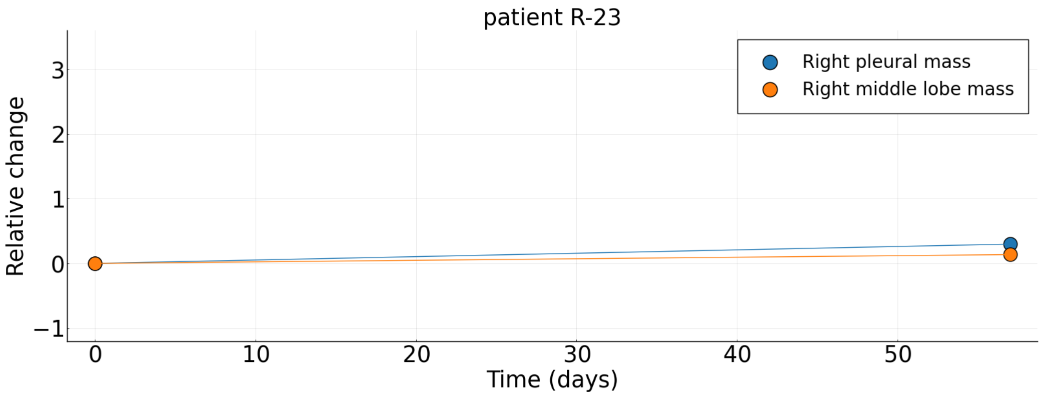


Figure S13: **Patient R-23. See main text Table 1 for details.**

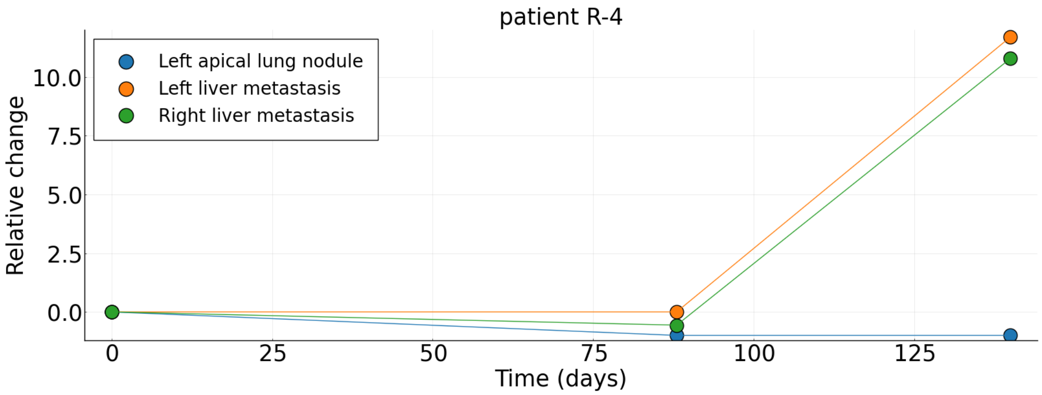


Figure S14: **Patient R-4. See main text Table 1 for details.**

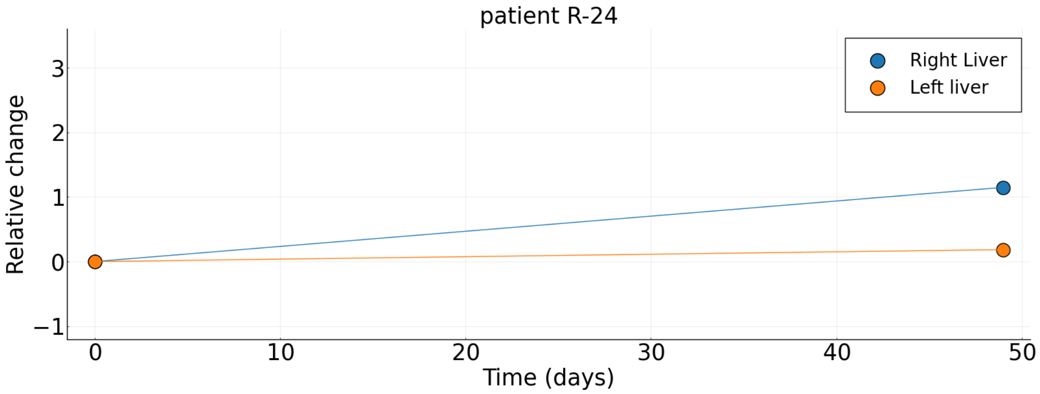


Figure S15: **Patient R-24. See main text Table 1 for details.**

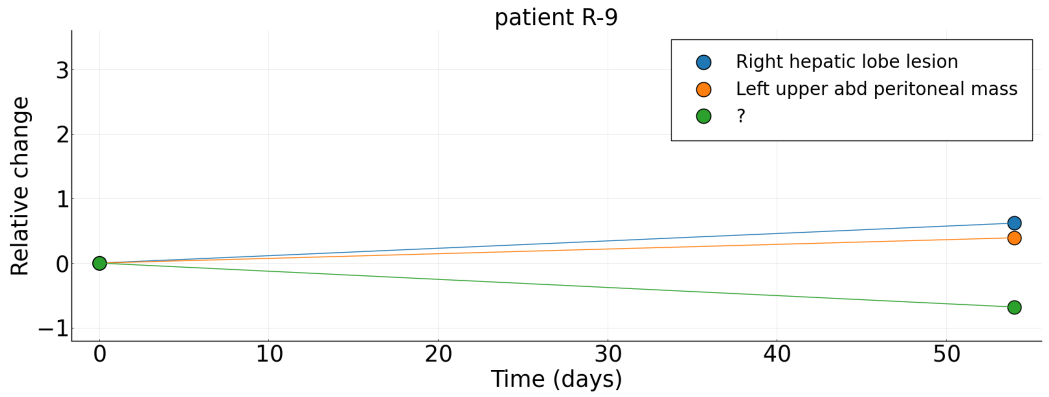


Figure S16: **Patient R-9. See main text Table 1 for details.**

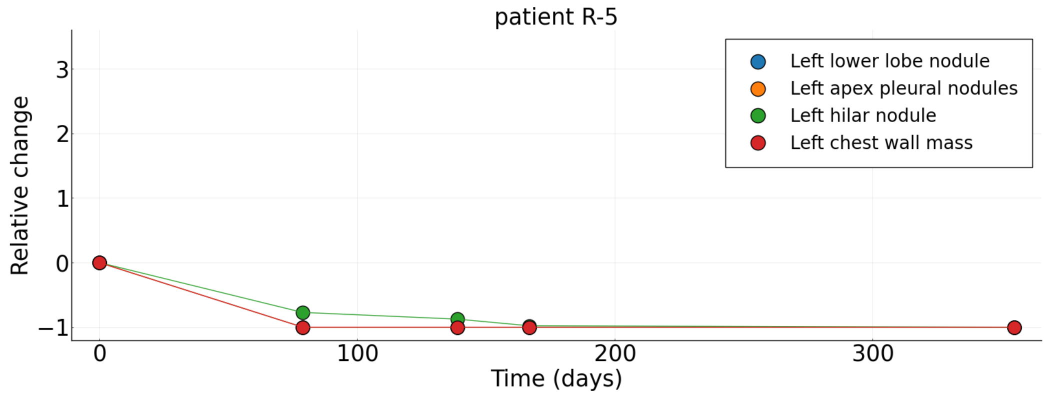


Figure S17: **Patient R-5. See main text Table 1 for details.**

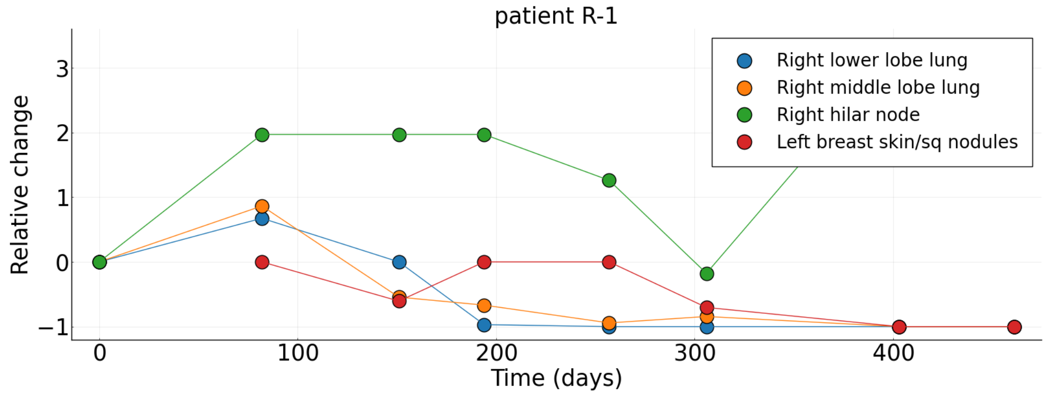


Figure S18: **Patient R-1. See main text Table 1 for details.**

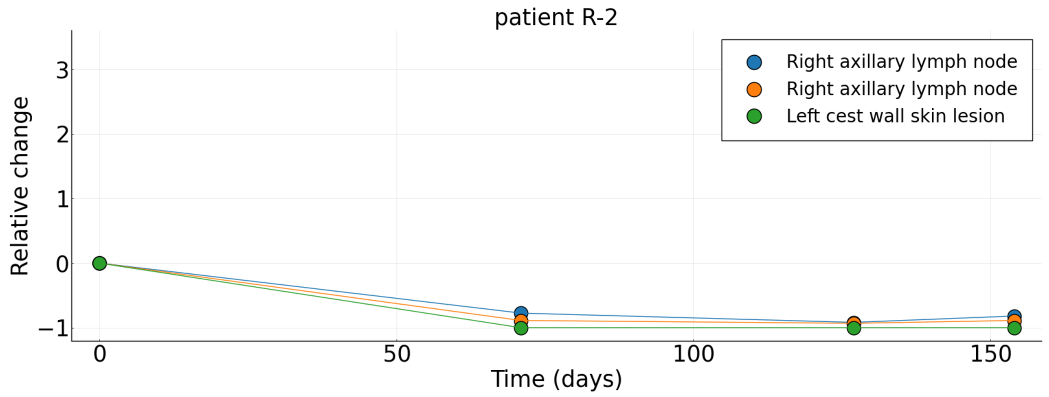


Figure S19: **Patient R-2. See main text Table 1 for details.**

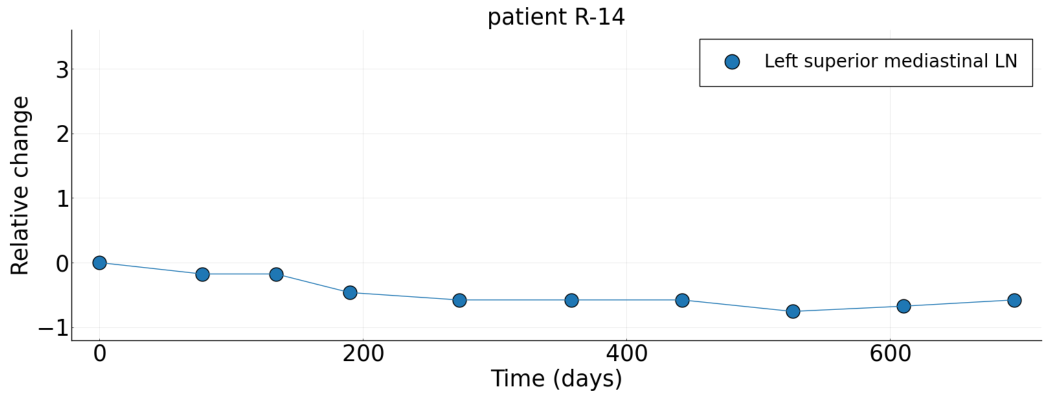


Figure S20: **Patient R-14. See main text Table 1 for details.**

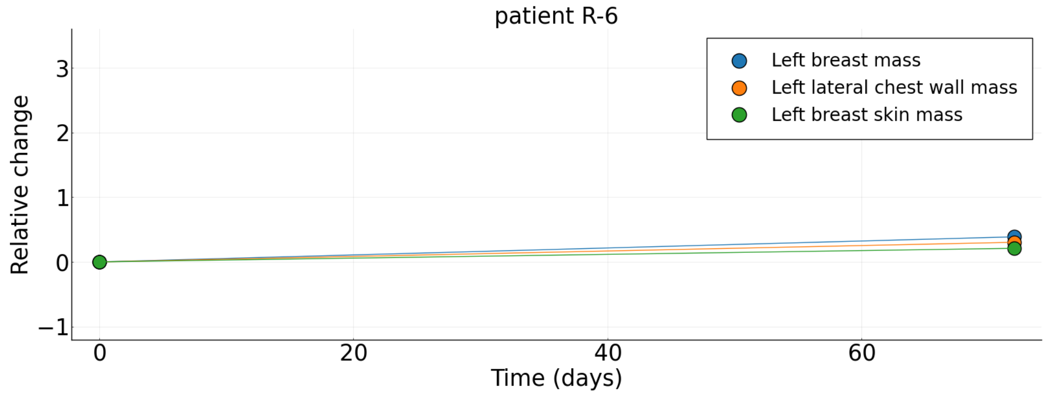


Figure S21: **Patient R-6. See main text Table 1 for details.**

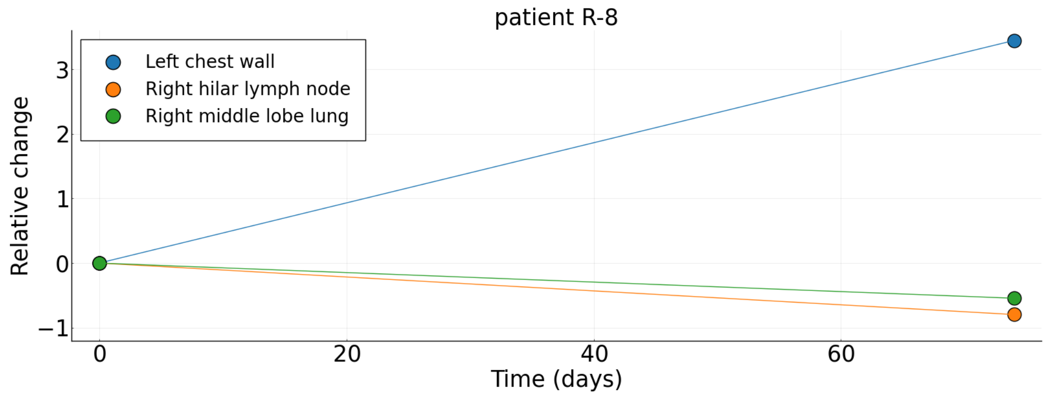


Figure S22: **Patient R-8. See main text Table 1 for details.**

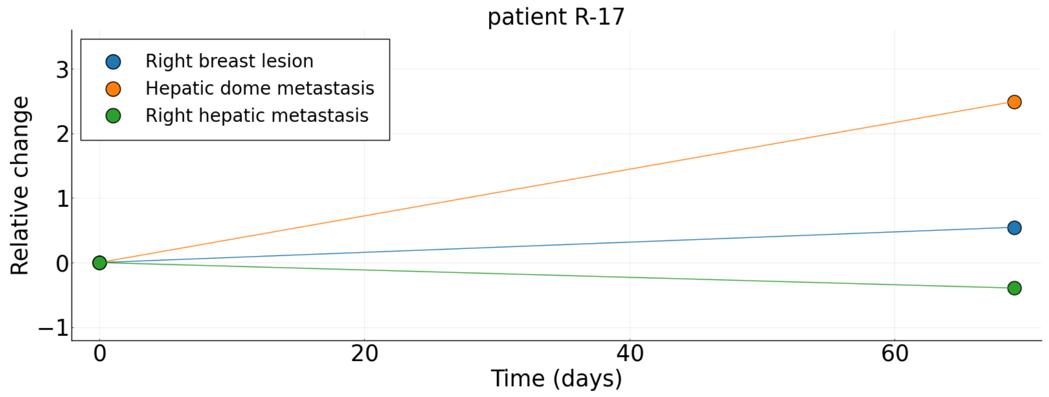


Figure S23: **Patient R-17. See main text Table 1 for details.**

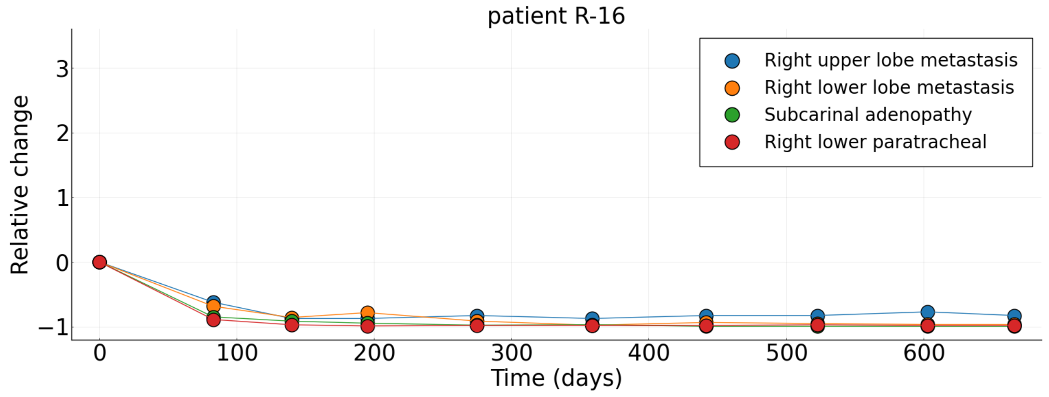


Figure S24: **Patient R-16. See main text Table 1 for details.**

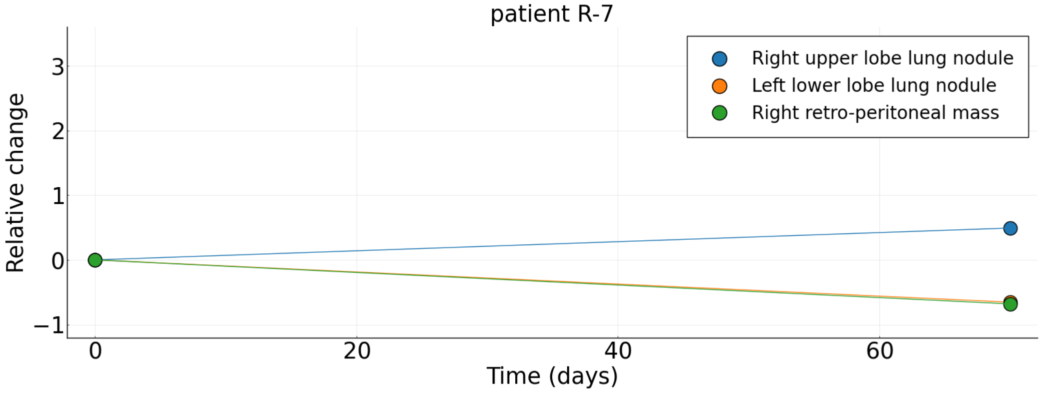


Figure S25: **Patient R-7. See main text Table 1 for details.**

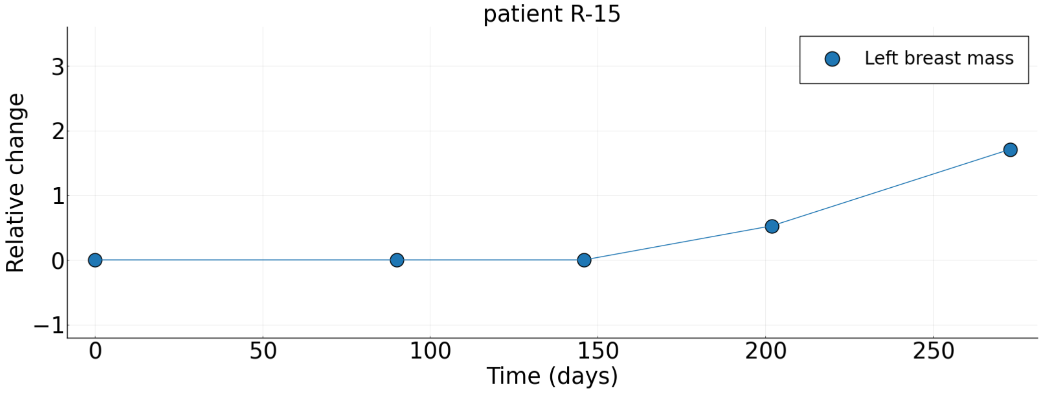


Figure S26: **Patient R-15. See main text Table 1 for details.**

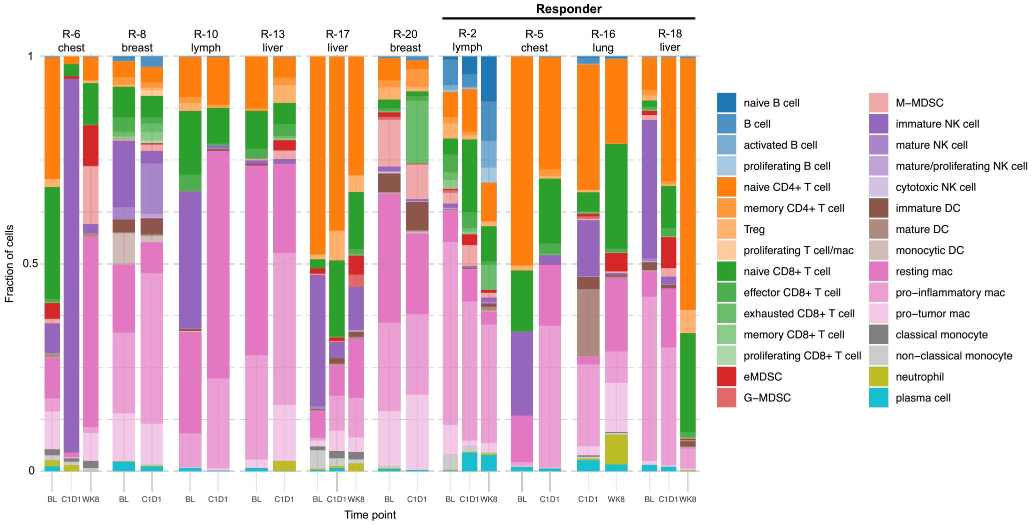


Figure S27: **IMC subclusters.** IMC subclusters for all immune cell subpopulations.

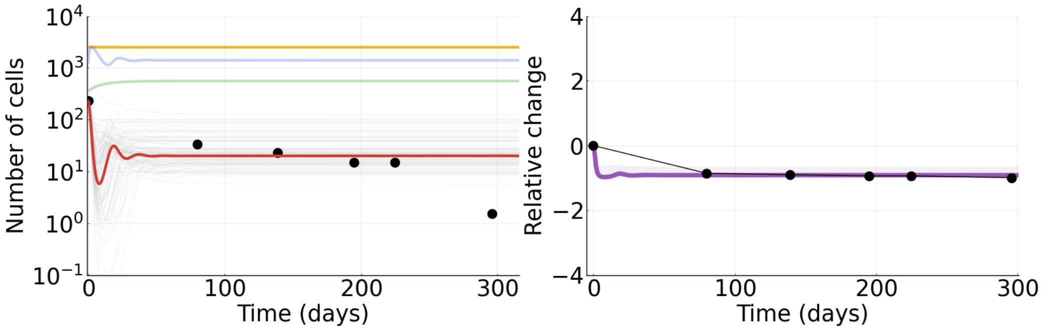


Figure S28: **Tumor 1.** Tumor data fit to mathematical model (model parameters set to be the median of their respective posterior distribution). Left panel: black dots represent the data, the red line represents tumor cells, the yellow line represents MDSCs, the green line represents NK cells, and the blue line represents CTL cells. Gray lines represent individual tumor cell population trajectories from the MCMC. Right panel: black dots represent the data, the purple line represents the tumor cell population, and the shaded area denotes the 90% credible interval where 90% of the posterior trajectories lie.

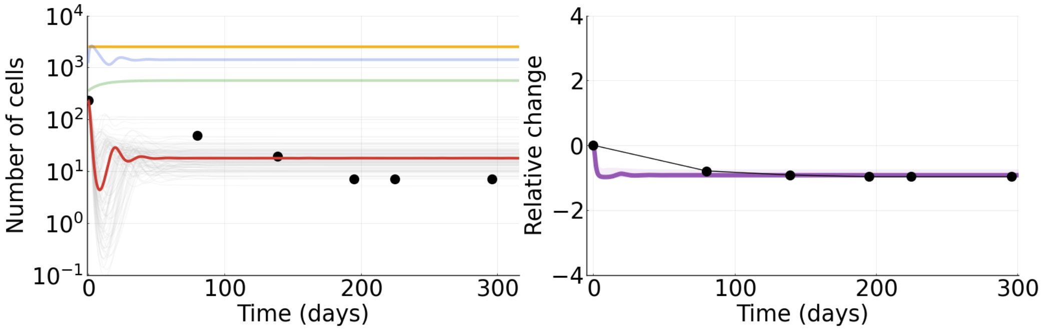


Figure S29: **Tumor 2.** Tumor data fit to mathematical model (model parameters set to be the median of their respective posterior distribution). Left panel: black dots represent the data, the red line represents tumor cells, the yellow line represents MDSCs, the green line represents NK cells, and the blue line represents CTL cells. Gray lines represent individual tumor cell population trajectories from the MCMC. Right panel: black dots represent the data, the purple line represents the tumor cell population, and the shaded area denotes the 90% credible interval where 90% of the posterior trajectories lie.

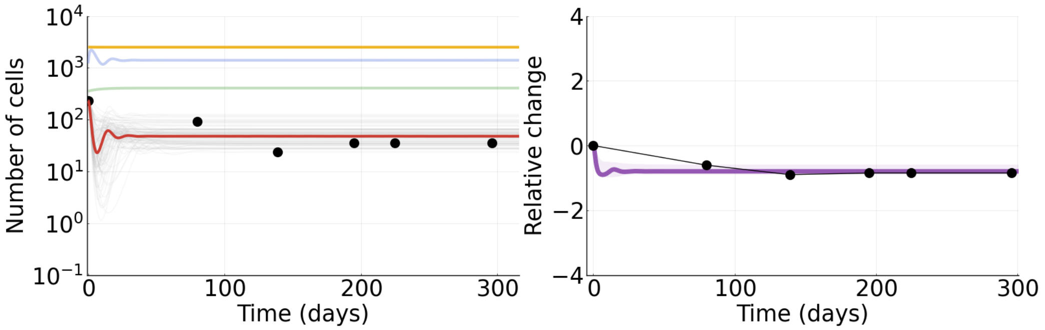


Figure S30: **Tumor 3.** Tumor data fit to mathematical model (model parameters set to be the median of their respective posterior distribution). Left panel: black dots represent the data, the red line represents tumor cells, the yellow line represents MDSCs, the green line represents NK cells, and the blue line represents CTL cells. Gray lines represent individual tumor cell population trajectories from the MCMC. Right panel: black dots represent the data, the purple line represents the tumor cell population, and the shaded area denotes the 90% credible interval where 90% of the posterior trajectories lie.

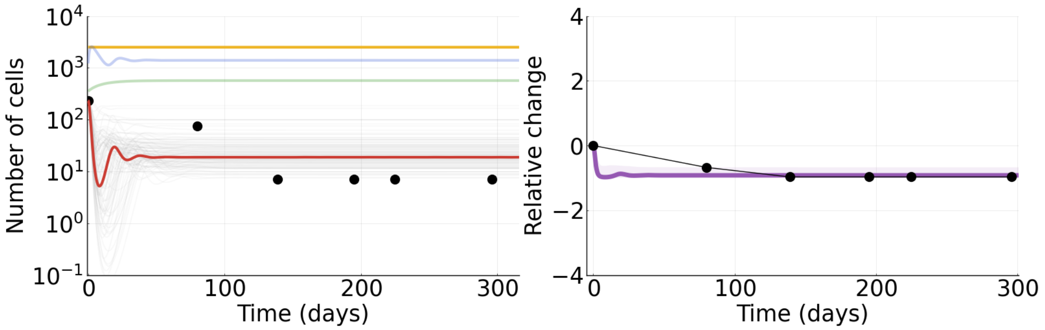


Figure S31: **Tumor 4.** Tumor data fit to mathematical model (model parameters set to be the median of their respective posterior distribution). Left panel: black dots represent the data, the red line represents tumor cells, the yellow line represents MDSCs, the green line represents NK cells, and the blue line represents CTL cells. Gray lines represent individual tumor cell population trajectories from the MCMC. Right panel: black dots represent the data, the purple line represents the tumor cell population, and the shaded area denotes the 90% credible interval where 90% of the posterior trajectories lie.

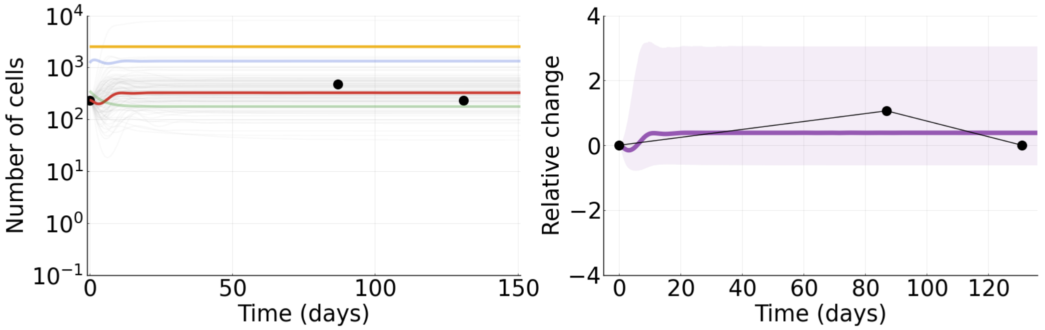


Figure S32: **Tumor 5.** Tumor data fit to mathematical model (model parameters set to be the median of their respective posterior distribution). Left panel: black dots represent the data, the red line represents tumor cells, the yellow line represents MDSCs, the green line represents NK cells, and the blue line represents CTL cells. Gray lines represent individual tumor cell population trajectories from the MCMC. Right panel: black dots represent the data, the purple line represents the tumor cell population, and the shaded area denotes the 90% credible interval where 90% of the posterior trajectories lie.

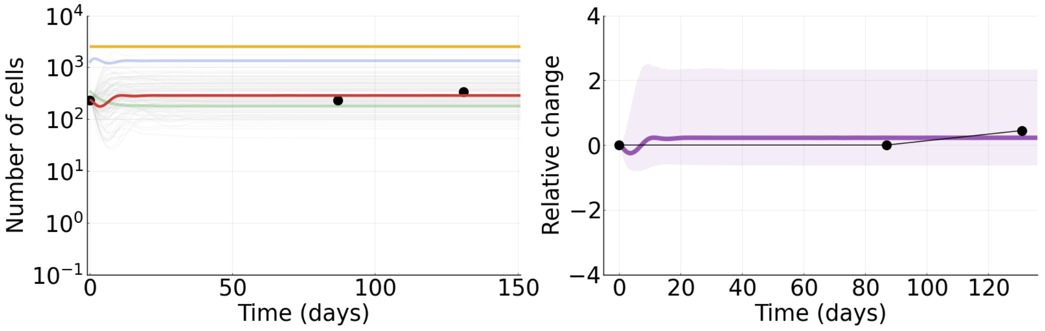


Figure S33: **Tumor 6.** Tumor data fit to mathematical model (model parameters set to be the median of their respective posterior distribution). Left panel: black dots represent the data, the red line represents tumor cells, the yellow line represents MDSCs, the green line represents NK cells, and the blue line represents CTL cells. Gray lines represent individual tumor cell population trajectories from the MCMC. Right panel: black dots represent the data, the purple line represents the tumor cell population, and the shaded area denotes the 90% credible interval where 90% of the posterior trajectories lie.

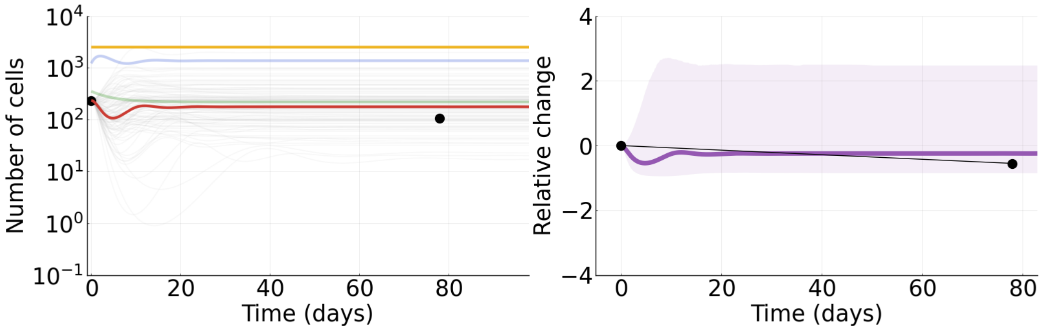


Figure S34: **Tumor 7.** Tumor data fit to mathematical model (model parameters set to be the median of their respective posterior distribution). Left panel: black dots represent the data, the red line represents tumor cells, the yellow line represents MDSCs, the green line represents NK cells, and the blue line represents CTL cells. Gray lines represent individual tumor cell population trajectories from the MCMC. Right panel: black dots represent the data, the purple line represents the tumor cell population, and the shaded area denotes the 90% credible interval where 90% of the posterior trajectories lie.

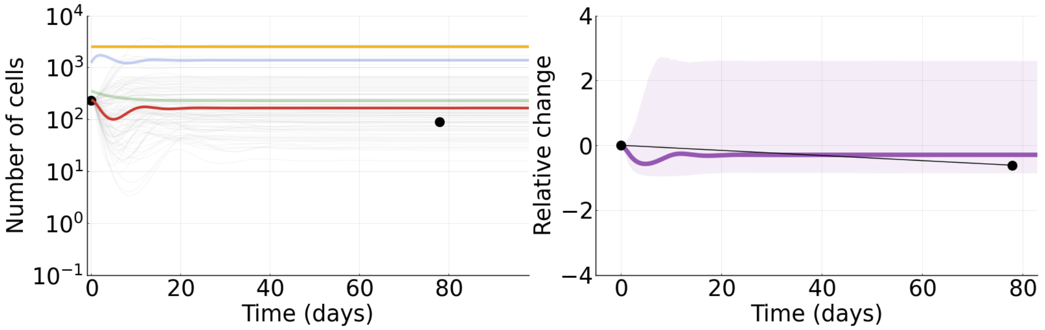


Figure S35: **Tumor 8.** Tumor data fit to mathematical model (model parameters set to be the median of their respective posterior distribution). Left panel: black dots represent the data, the red line represents tumor cells, the yellow line represents MDSCs, the green line represents NK cells, and the blue line represents CTL cells. Gray lines represent individual tumor cell population trajectories from the MCMC. Right panel: black dots represent the data, the purple line represents the tumor cell population, and the shaded area denotes the 90% credible interval where 90% of the posterior trajectories lie.

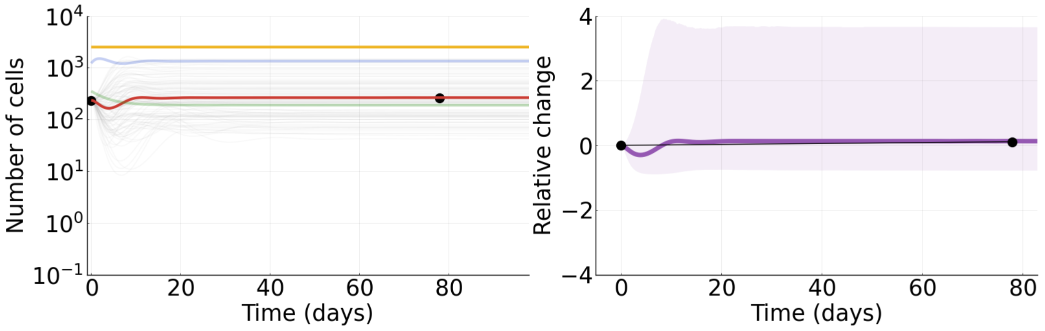


Figure S36: **Tumor 9.** Tumor data fit to mathematical model (model parameters set to be the median of their respective posterior distribution). Left panel: black dots represent the data, the red line represents tumor cells, the yellow line represents MDSCs, the green line represents NK cells, and the blue line represents CTL cells. Gray lines represent individual tumor cell population trajectories from the MCMC. Right panel: black dots represent the data, the purple line represents the tumor cell population, and the shaded area denotes the 90% credible interval where 90% of the posterior trajectories lie.

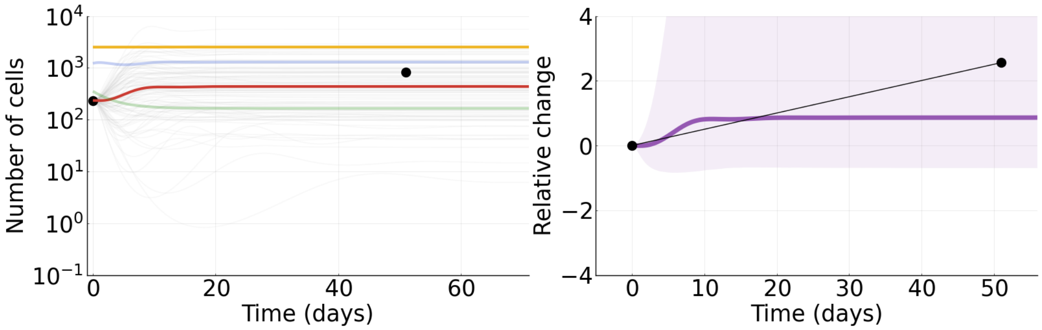


Figure S37: **Tumor 10.** Tumor data fit to mathematical model (model parameters set to be the median of their respective posterior distribution). Left panel: black dots represent the data, the red line represents tumor cells, the yellow line represents MDSCs, the green line represents NK cells, and the blue line represents CTL cells. Gray lines represent individual tumor cell population trajectories from the MCMC. Right panel: black dots represent the data, the purple line represents the tumor cell population, and the shaded area denotes the 90% credible interval where 90% of the posterior trajectories lie.

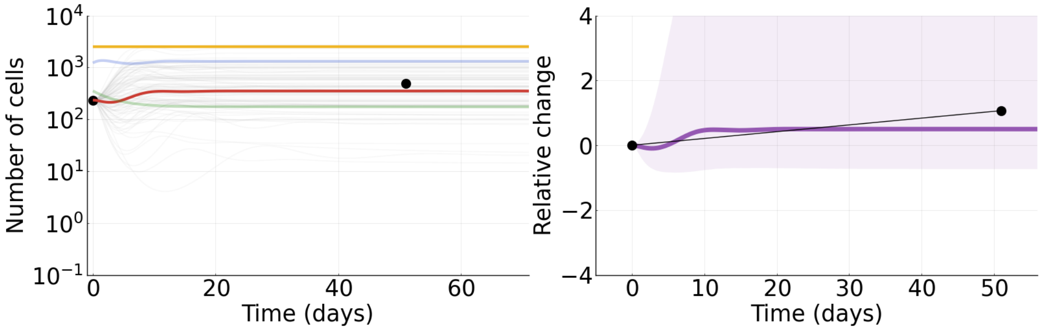


Figure S38: **Tumor 11.** Tumor data fit to mathematical model (model parameters set to be the median of their respective posterior distribution). Left panel: black dots represent the data, the red line represents tumor cells, the yellow line represents MDSCs, the green line represents NK cells, and the blue line represents CTL cells. Gray lines represent individual tumor cell population trajectories from the MCMC. Right panel: black dots represent the data, the purple line represents the tumor cell population, and the shaded area denotes the 90% credible interval where 90% of the posterior trajectories lie.

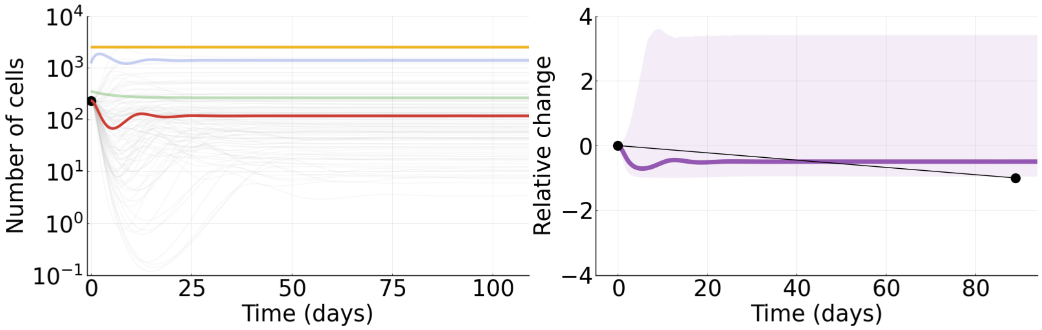


Figure S39: **Tumor 12.** Tumor data fit to mathematical model (model parameters set to be the median of their respective posterior distribution). Left panel: black dots represent the data, the red line represents tumor cells, the yellow line represents MDSCs, the green line represents NK cells, and the blue line represents CTL cells. Gray lines represent individual tumor cell population trajectories from the MCMC. Right panel: black dots represent the data, the purple line represents the tumor cell population, and the shaded area denotes the 90% credible interval where 90% of the posterior trajectories lie.

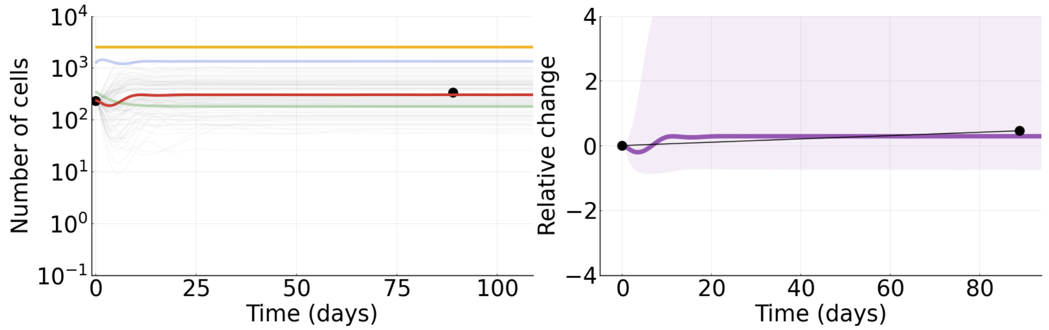


Figure S40: **Tumor 13.** Tumor data fit to mathematical model (model parameters set to be the median of their respective posterior distribution). Left panel: black dots represent the data, the red line represents tumor cells, the yellow line represents MDSCs, the green line represents NK cells, and the blue line represents CTL cells. Gray lines represent individual tumor cell population trajectories from the MCMC. Right panel: black dots represent the data, the purple line represents the tumor cell population, and the shaded area denotes the 90% credible interval where 90% of the posterior trajectories lie.

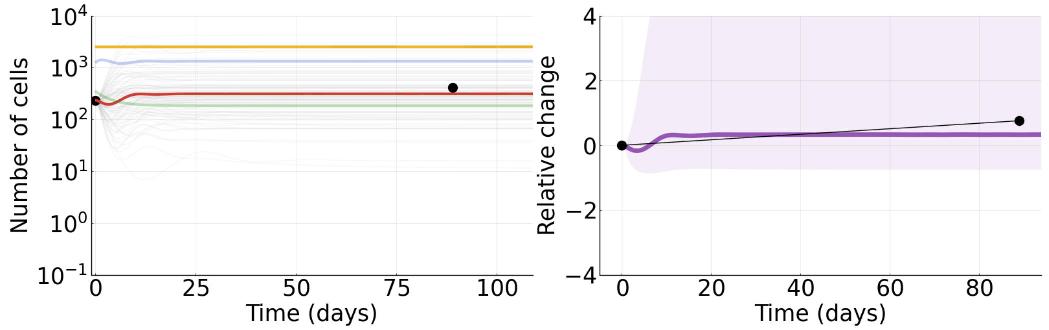


Figure S41: **Tumor 14.** Tumor data fit to mathematical model (model parameters set to be the median of their respective posterior distribution). Left panel: black dots represent the data, the red line represents tumor cells, the yellow line represents MDSCs, the green line represents NK cells, and the blue line represents CTL cells. Gray lines represent individual tumor cell population trajectories from the MCMC. Right panel: black dots represent the data, the purple line represents the tumor cell population, and the shaded area denotes the 90% credible interval where 90% of the posterior trajectories lie.

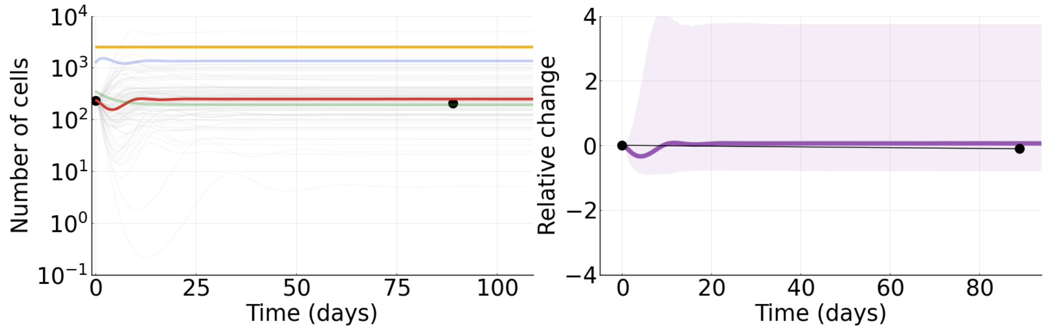


Figure S42: **Tumor 15.** Tumor data fit to mathematical model (model parameters set to be the median of their respective posterior distribution). Left panel: black dots represent the data, the red line represents tumor cells, the yellow line represents MDSCs, the green line represents NK cells, and the blue line represents CTL cells. Gray lines represent individual tumor cell population trajectories from the MCMC. Right panel: black dots represent the data, the purple line represents the tumor cell population, and the shaded area denotes the 90% credible interval where 90% of the posterior trajectories lie.

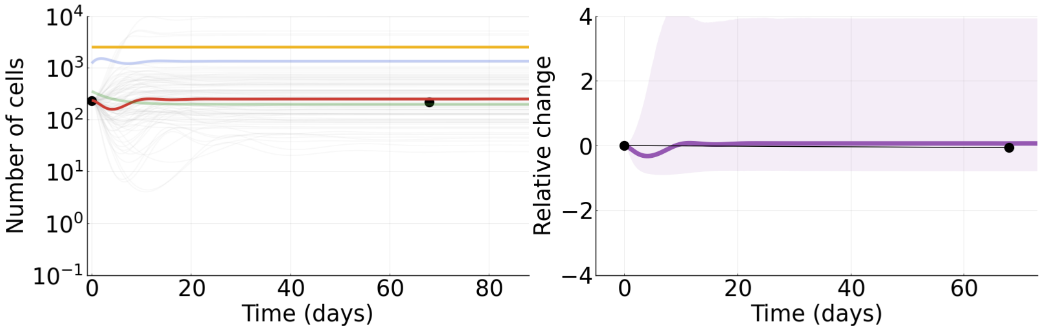


Figure S43: **Tumor 16.** Tumor data fit to mathematical model (model parameters set to be the median of their respective posterior distribution). Left panel: black dots represent the data, the red line represents tumor cells, the yellow line represents MDSCs, the green line represents NK cells, and the blue line represents CTL cells. Gray lines represent individual tumor cell population trajectories from the MCMC. Right panel: black dots represent the data, the purple line represents the tumor cell population, and the shaded area denotes the 90% credible interval where 90% of the posterior trajectories lie.

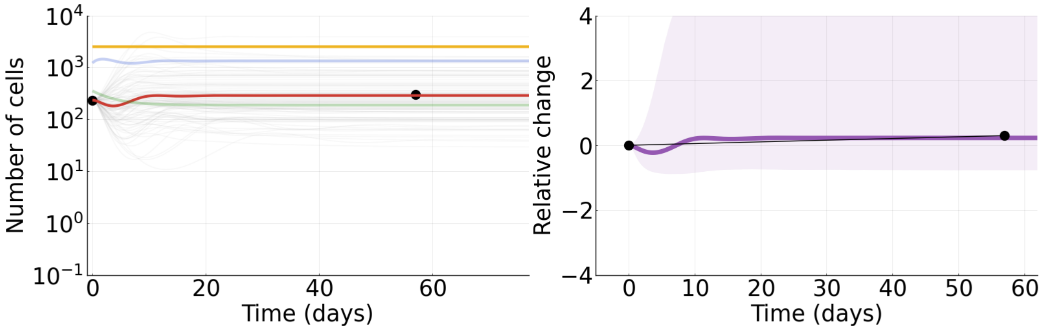


Figure S44: **Tumor 17.** Tumor data fit to mathematical model (model parameters set to be the median of their respective posterior distribution). Left panel: black dots represent the data, the red line represents tumor cells, the yellow line represents MDSCs, the green line represents NK cells, and the blue line represents CTL cells. Gray lines represent individual tumor cell population trajectories from the MCMC. Right panel: black dots represent the data, the purple line represents the tumor cell population, and the shaded area denotes the 90% credible interval where 90% of the posterior trajectories lie.

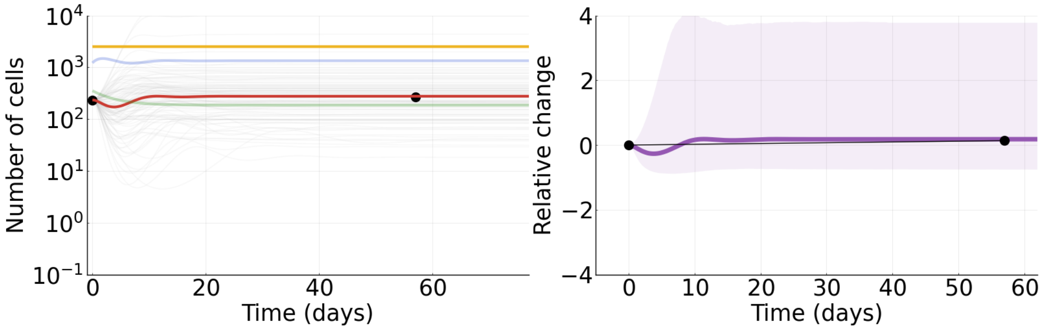


Figure S45: **Tumor 18.** Tumor data fit to mathematical model (model parameters set to be the median of their respective posterior distribution). Left panel: black dots represent the data, the red line represents tumor cells, the yellow line represents MDSCs, the green line represents NK cells, and the blue line represents CTL cells. Gray lines represent individual tumor cell population trajectories from the MCMC. Right panel: black dots represent the data, the purple line represents the tumor cell population, and the shaded area denotes the 90% credible interval where 90% of the posterior trajectories lie.

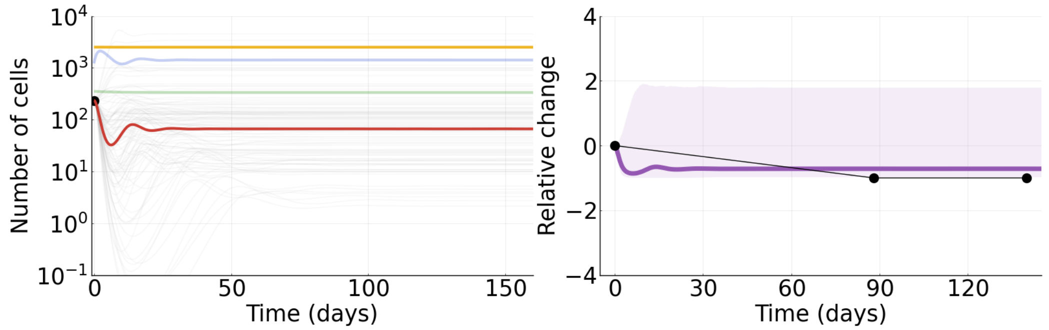


Figure S46: **Tumor 19.** Tumor data fit to mathematical model (model parameters set to be the median of their respective posterior distribution). Left panel: black dots represent the data, the red line represents tumor cells, the yellow line represents MDSCs, the green line represents NK cells, and the blue line represents CTL cells. Gray lines represent individual tumor cell population trajectories from the MCMC. Right panel: black dots represent the data, the purple line represents the tumor cell population, and the shaded area denotes the 90% credible interval where 90% of the posterior trajectories lie.

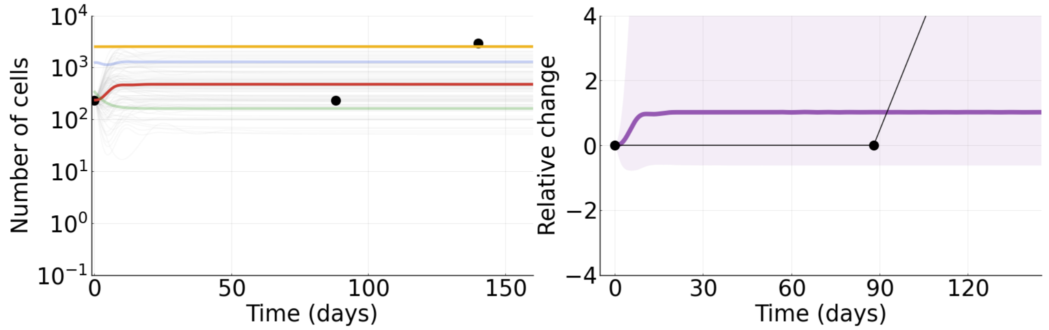


Figure S47: **Tumor 20.** Tumor data fit to mathematical model (model parameters set to be the median of their respective posterior distribution). Left panel: black dots represent the data, the red line represents tumor cells, the yellow line represents MDSCs, the green line represents NK cells, and the blue line represents CTL cells. Gray lines represent individual tumor cell population trajectories from the MCMC. Right panel: black dots represent the data, the purple line represents the tumor cell population, and the shaded area denotes the 90% credible interval where 90% of the posterior trajectories lie.

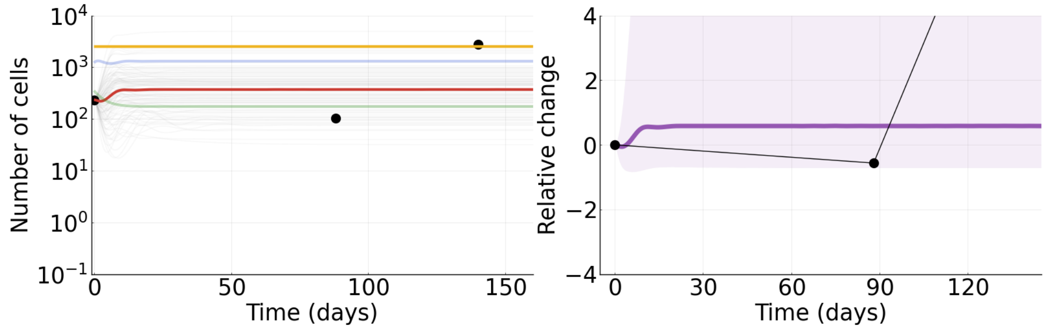


Figure S48: **Tumor 21.** Tumor data fit to mathematical model (model parameters set to be the median of their respective posterior distribution). Left panel: black dots represent the data, the red line represents tumor cells, the yellow line represents MDSCs, the green line represents NK cells, and the blue line represents CTL cells. Gray lines represent individual tumor cell population trajectories from the MCMC. Right panel: black dots represent the data, the purple line represents the tumor cell population, and the shaded area denotes the 90% credible interval where 90% of the posterior trajectories lie.

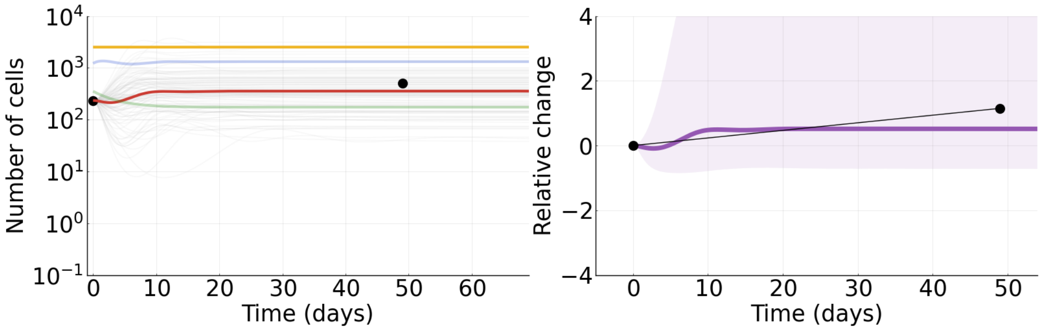


Figure S49: **Tumor 22.** Tumor data fit to mathematical model (model parameters set to be the median of their respective posterior distribution). Left panel: black dots represent the data, the red line represents tumor cells, the yellow line represents MDSCs, the green line represents NK cells, and the blue line represents CTL cells. Gray lines represent individual tumor cell population trajectories from the MCMC. Right panel: black dots represent the data, the purple line represents the tumor cell population, and the shaded area denotes the 90% credible interval where 90% of the posterior trajectories lie.

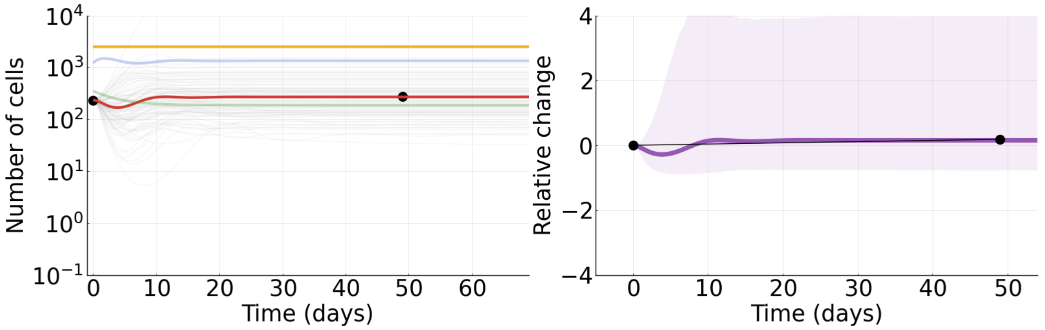


Figure S50: **Tumor 23.** Tumor data fit to mathematical model (model parameters set to be the median of their respective posterior distribution). Left panel: black dots represent the data, the red line represents tumor cells, the yellow line represents MDSCs, the green line represents NK cells, and the blue line represents CTL cells. Gray lines represent individual tumor cell population trajectories from the MCMC. Right panel: black dots represent the data, the purple line represents the tumor cell population, and the shaded area denotes the 90% credible interval where 90% of the posterior trajectories lie.

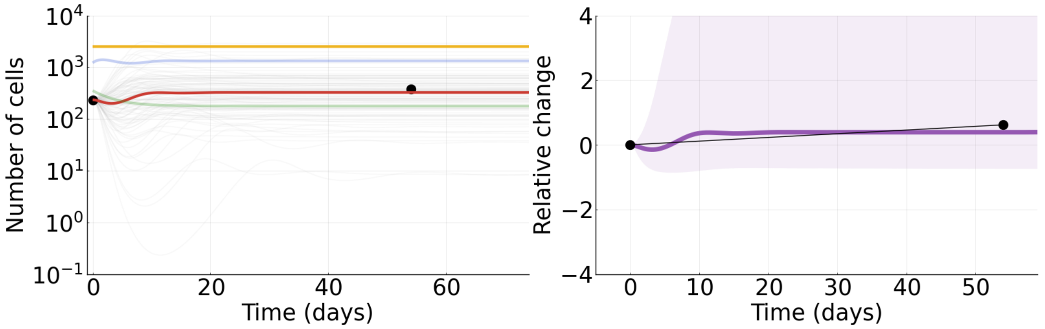


Figure S51: **Tumor 24.** Tumor data fit to mathematical model (model parameters set to be the median of their respective posterior distribution). Left panel: black dots represent the data, the red line represents tumor cells, the yellow line represents MDSCs, the green line represents NK cells, and the blue line represents CTL cells. Gray lines represent individual tumor cell population trajectories from the MCMC. Right panel: black dots represent the data, the purple line represents the tumor cell population, and the shaded area denotes the 90% credible interval where 90% of the posterior trajectories lie.

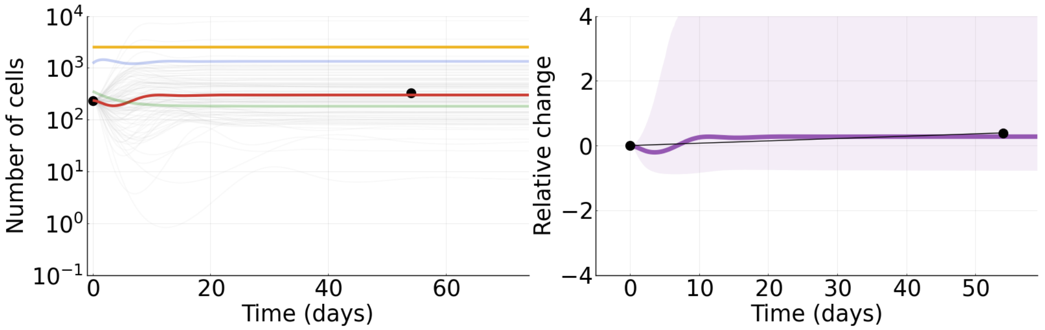


Figure S52: **Tumor 25.** Tumor data fit to mathematical model (model parameters set to be the median of their respective posterior distribution). Left panel: black dots represent the data, the red line represents tumor cells, the yellow line represents MDSCs, the green line represents NK cells, and the blue line represents CTL cells. Gray lines represent individual tumor cell population trajectories from the MCMC. Right panel: black dots represent the data, the purple line represents the tumor cell population, and the shaded area denotes the 90% credible interval where 90% of the posterior trajectories lie.

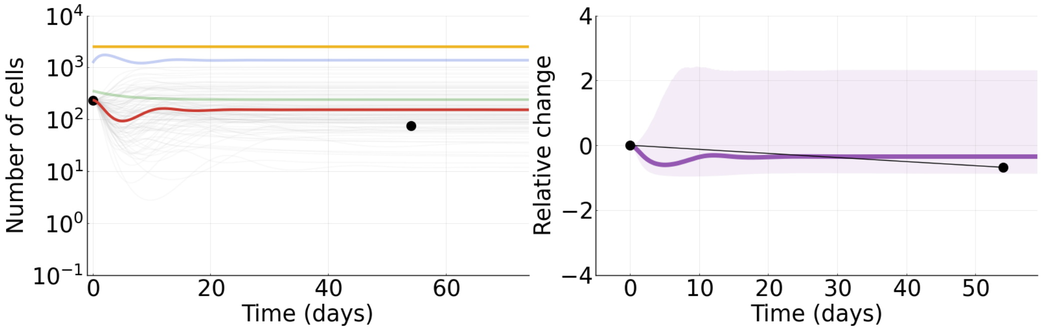


Figure S53: **Tumor 26.** Tumor data fit to mathematical model (model parameters set to be the median of their respective posterior distribution). Left panel: black dots represent the data, the red line represents tumor cells, the yellow line represents MDSCs, the green line represents NK cells, and the blue line represents CTL cells. Gray lines represent individual tumor cell population trajectories from the MCMC. Right panel: black dots represent the data, the purple line represents the tumor cell population, and the shaded area denotes the 90% credible interval where 90% of the posterior trajectories lie.

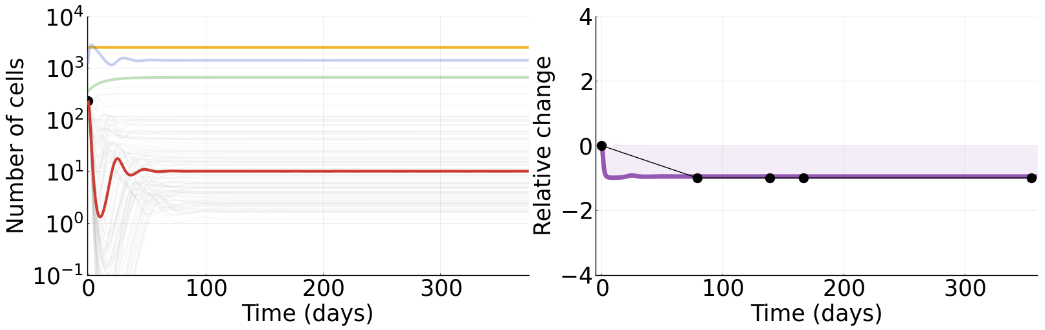


Figure S54: **Tumor 27.** Tumor data fit to mathematical model (model parameters set to be the median of their respective posterior distribution). Left panel: black dots represent the data, the red line represents tumor cells, the yellow line represents MDSCs, the green line represents NK cells, and the blue line represents CTL cells. Gray lines represent individual tumor cell population trajectories from the MCMC. Right panel: black dots represent the data, the purple line represents the tumor cell population, and the shaded area denotes the 90% credible interval where 90% of the posterior trajectories lie.

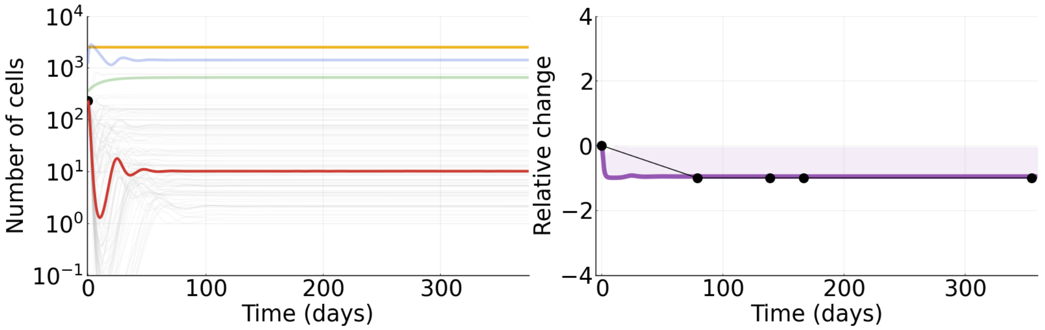


Figure S55: **Tumor 28.** Tumor data fit to mathematical model (model parameters set to be the median of their respective posterior distribution). Left panel: black dots represent the data, the red line represents tumor cells, the yellow line represents MDSCs, the green line represents NK cells, and the blue line represents CTL cells. Gray lines represent individual tumor cell population trajectories from the MCMC. Right panel: black dots represent the data, the purple line represents the tumor cell population, and the shaded area denotes the 90% credible interval where 90% of the posterior trajectories lie.

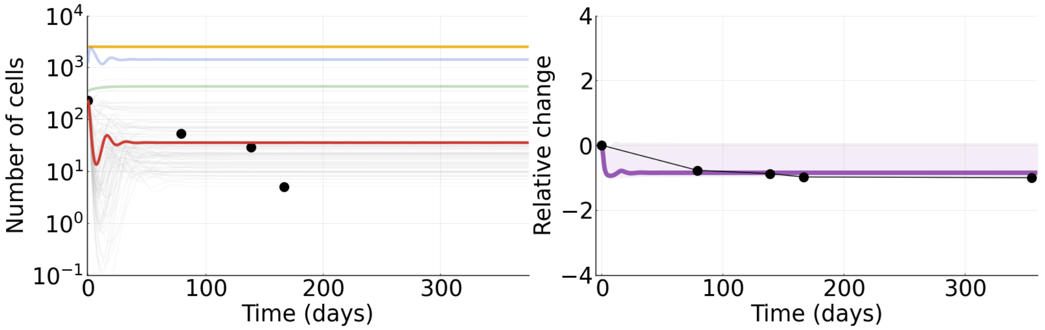


Figure S56: **Tumor 29.** Tumor data fit to mathematical model (model parameters set to be the median of their respective posterior distribution). Left panel: black dots represent the data, the red line represents tumor cells, the yellow line represents MDSCs, the green line represents NK cells, and the blue line represents CTL cells. Gray lines represent individual tumor cell population trajectories from the MCMC. Right panel: black dots represent the data, the purple line represents the tumor cell population, and the shaded area denotes the 90% credible interval where 90% of the posterior trajectories lie.

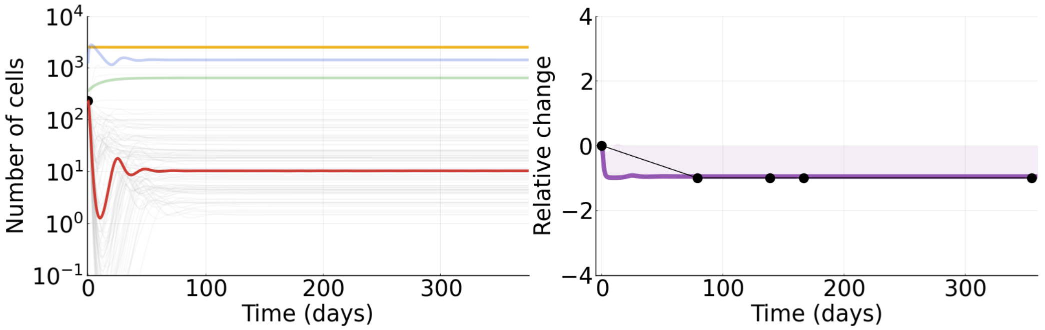


Figure S57: **Tumor 30.** Tumor data fit to mathematical model (model parameters set to be the median of their respective posterior distribution). Left panel: black dots represent the data, the red line represents tumor cells, the yellow line represents MDSCs, the green line represents NK cells, and the blue line represents CTL cells. Gray lines represent individual tumor cell population trajectories from the MCMC. Right panel: black dots represent the data, the purple line represents the tumor cell population, and the shaded area denotes the 90% credible interval where 90% of the posterior trajectories lie.

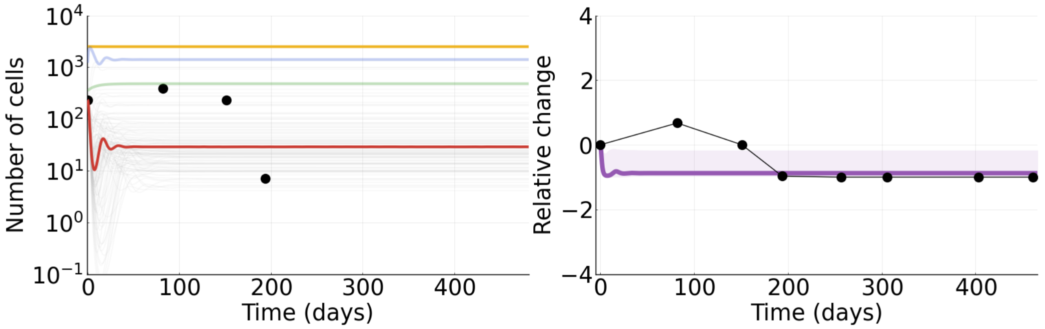


Figure S58: **Tumor 31.** Tumor data fit to mathematical model (model parameters set to be the median of their respective posterior distribution). Left panel: black dots represent the data, the red line represents tumor cells, the yellow line represents MDSCs, the green line represents NK cells, and the blue line represents CTL cells. Gray lines represent individual tumor cell population trajectories from the MCMC. Right panel: black dots represent the data, the purple line represents the tumor cell population, and the shaded area denotes the 90% credible interval where 90% of the posterior trajectories lie.

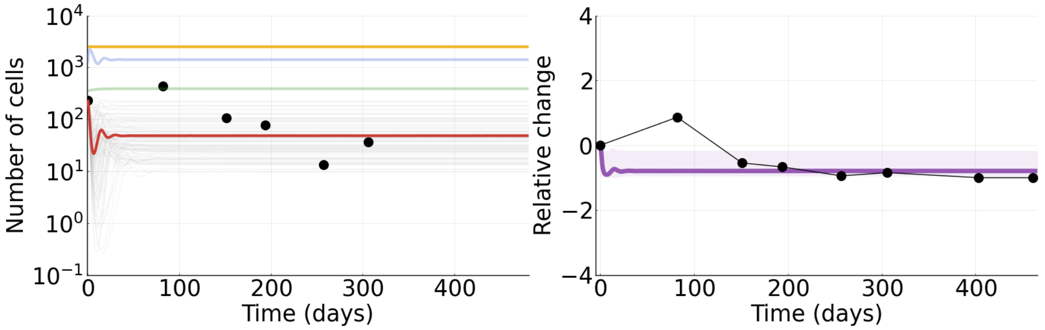


Figure S59: **Tumor 32.** Tumor data fit to mathematical model (model parameters set to be the median of their respective posterior distribution). Left panel: black dots represent the data, the red line represents tumor cells, the yellow line represents MDSCs, the green line represents NK cells, and the blue line represents CTL cells. Gray lines represent individual tumor cell population trajectories from the MCMC. Right panel: black dots represent the data, the purple line represents the tumor cell population, and the shaded area denotes the 90% credible interval where 90% of the posterior trajectories lie.

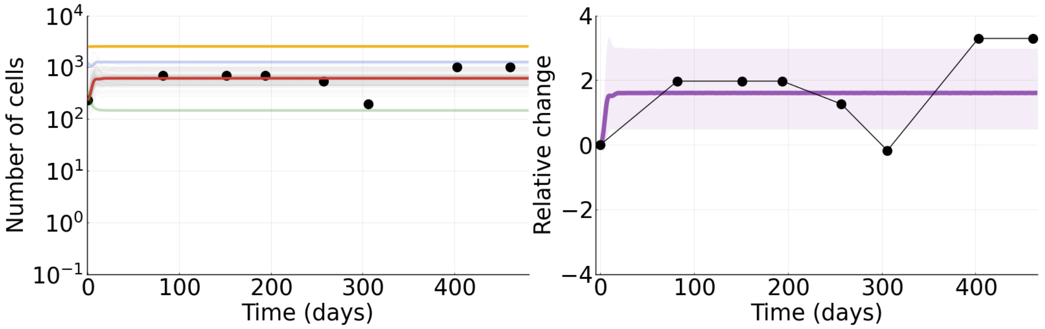


Figure S60: **Tumor 33.** Tumor data fit to mathematical model (model parameters set to be the median of their respective posterior distribution). Left panel: black dots represent the data, the red line represents tumor cells, the yellow line represents MDSCs, the green line represents NK cells, and the blue line represents CTL cells. Gray lines represent individual tumor cell population trajectories from the MCMC. Right panel: black dots represent the data, the purple line represents the tumor cell population, and the shaded area denotes the 90% credible interval where 90% of the posterior trajectories lie.

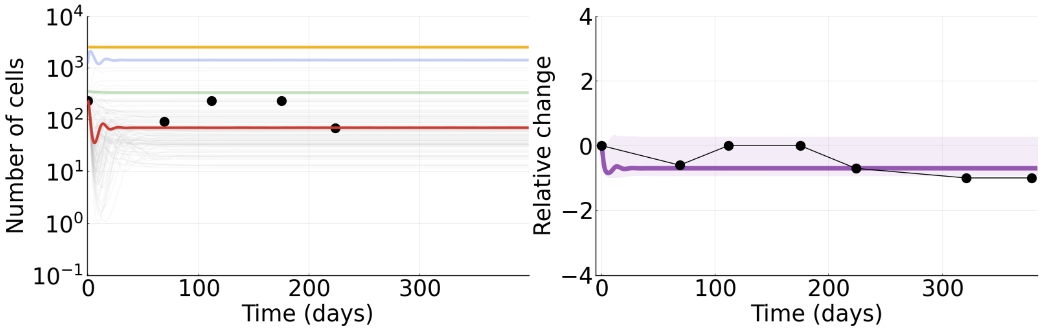


Figure S61: **Tumor 34.** Tumor data fit to mathematical model (model parameters set to be the median of their respective posterior distribution). Left panel: black dots represent the data, the red line represents tumor cells, the yellow line represents MDSCs, the green line represents NK cells, and the blue line represents CTL cells. Gray lines represent individual tumor cell population trajectories from the MCMC. Right panel: black dots represent the data, the purple line represents the tumor cell population, and the shaded area denotes the 90% credible interval where 90% of the posterior trajectories lie.

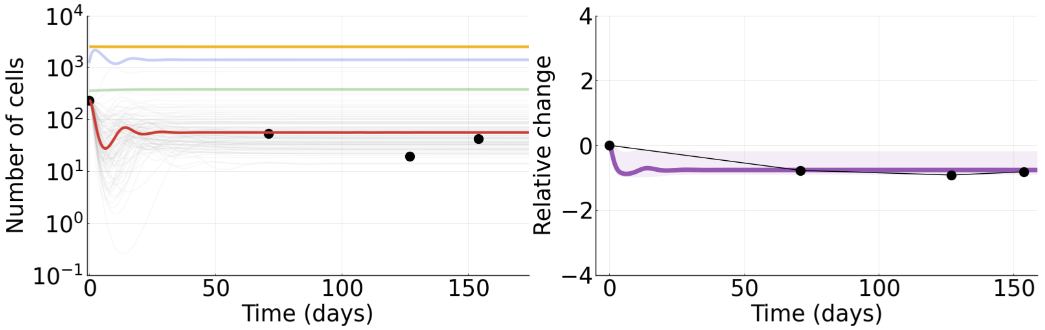


Figure S62: **Tumor 35.** Tumor data fit to mathematical model (model parameters set to be the median of their respective posterior distribution). Left panel: black dots represent the data, the red line represents tumor cells, the yellow line represents MDSCs, the green line represents NK cells, and the blue line represents CTL cells. Gray lines represent individual tumor cell population trajectories from the MCMC. Right panel: black dots represent the data, the purple line represents the tumor cell population, and the shaded area denotes the 90% credible interval where 90% of the posterior trajectories lie.

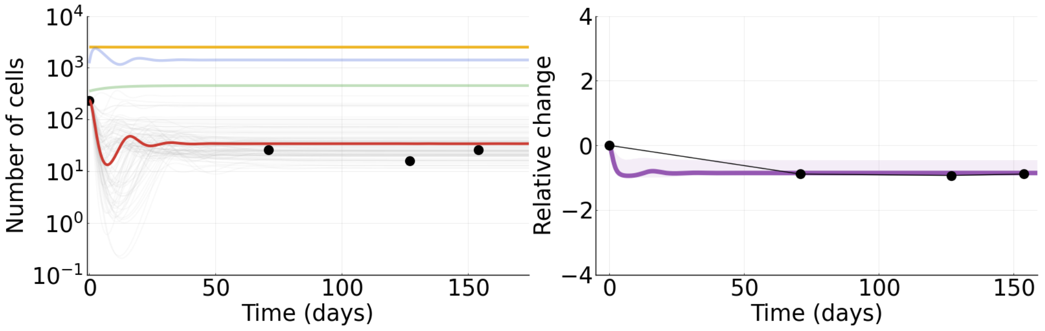


Figure S63: **Tumor 36.** Tumor data fit to mathematical model (model parameters set to be the median of their respective posterior distribution). Left panel: black dots represent the data, the red line represents tumor cells, the yellow line represents MDSCs, the green line represents NK cells, and the blue line represents CTL cells. Gray lines represent individual tumor cell population trajectories from the MCMC. Right panel: black dots represent the data, the purple line represents the tumor cell population, and the shaded area denotes the 90% credible interval where 90% of the posterior trajectories lie.

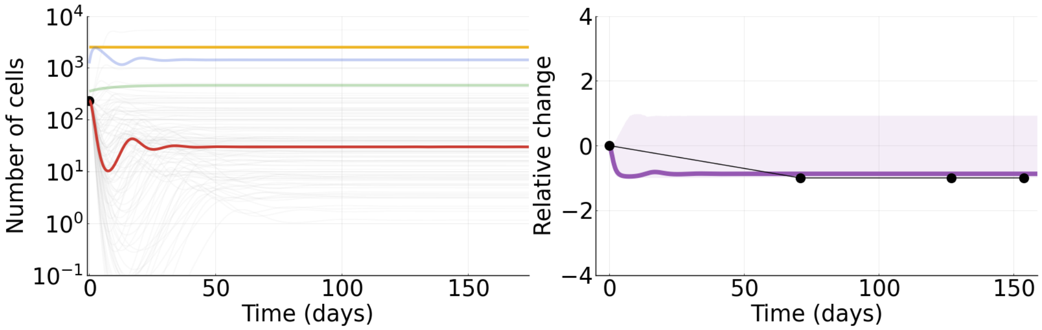


Figure S64: **Tumor 37.** Tumor data fit to mathematical model (model parameters set to be the median of their respective posterior distribution). Left panel: black dots represent the data, the red line represents tumor cells, the yellow line represents MDSCs, the green line represents NK cells, and the blue line represents CTL cells. Gray lines represent individual tumor cell population trajectories from the MCMC. Right panel: black dots represent the data, the purple line represents the tumor cell population, and the shaded area denotes the 90% credible interval where 90% of the posterior trajectories lie.

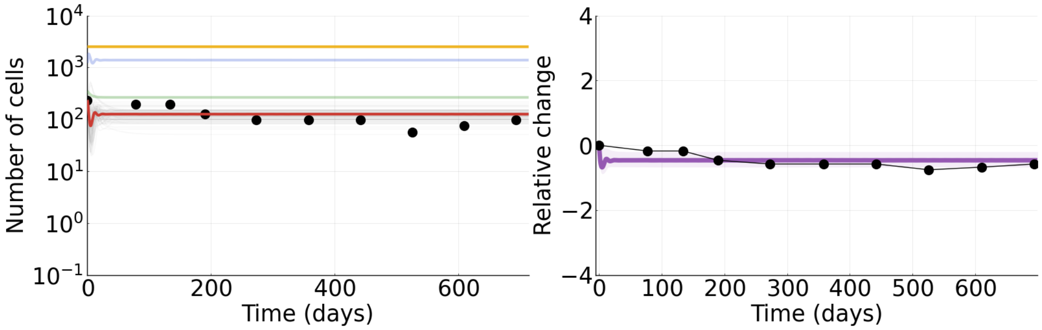


Figure S65: **Tumor 38.** Tumor data fit to mathematical model (model parameters set to be the median of their respective posterior distribution). Left panel: black dots represent the data, the red line represents tumor cells, the yellow line represents MDSCs, the green line represents NK cells, and the blue line represents CTL cells. Gray lines represent individual tumor cell population trajectories from the MCMC. Right panel: black dots represent the data, the purple line represents the tumor cell population, and the shaded area denotes the 90% credible interval where 90% of the posterior trajectories lie.

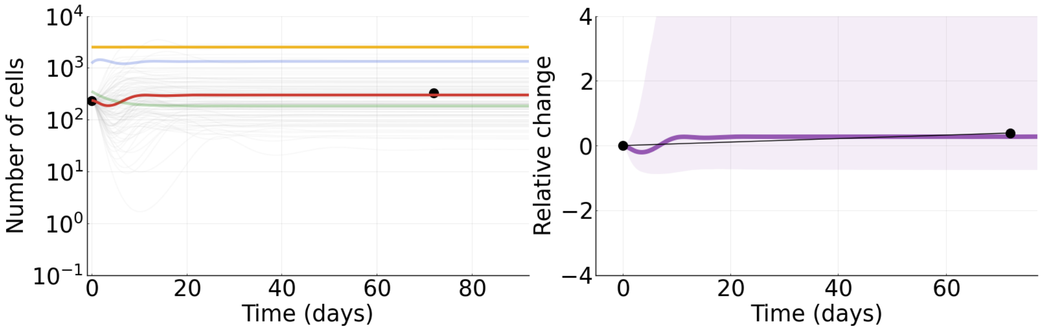


Figure S66: **Tumor 39.** Tumor data fit to mathematical model (model parameters set to be the median of their respective posterior distribution). Left panel: black dots represent the data, the red line represents tumor cells, the yellow line represents MDSCs, the green line represents NK cells, and the blue line represents CTL cells. Gray lines represent individual tumor cell population trajectories from the MCMC. Right panel: black dots represent the data, the purple line represents the tumor cell population, and the shaded area denotes the 90% credible interval where 90% of the posterior trajectories lie.

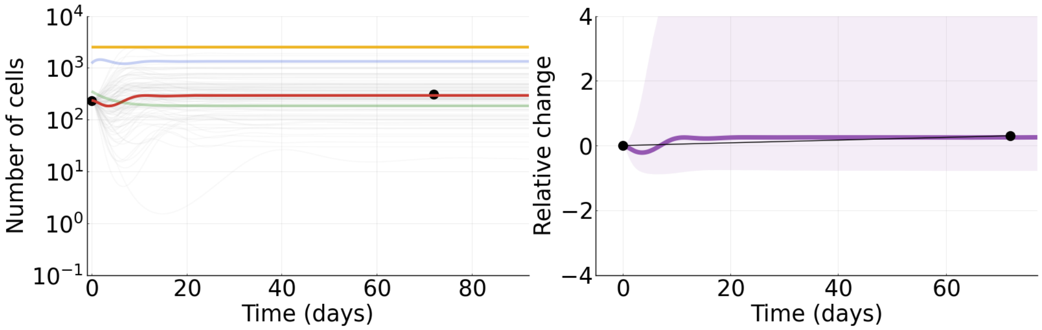


Figure S67: **Tumor 40.** Tumor data fit to mathematical model (model parameters set to be the median of their respective posterior distribution). Left panel: black dots represent the data, the red line represents tumor cells, the yellow line represents MDSCs, the green line represents NK cells, and the blue line represents CTL cells. Gray lines represent individual tumor cell population trajectories from the MCMC. Right panel: black dots represent the data, the purple line represents the tumor cell population, and the shaded area denotes the 90% credible interval where 90% of the posterior trajectories lie.

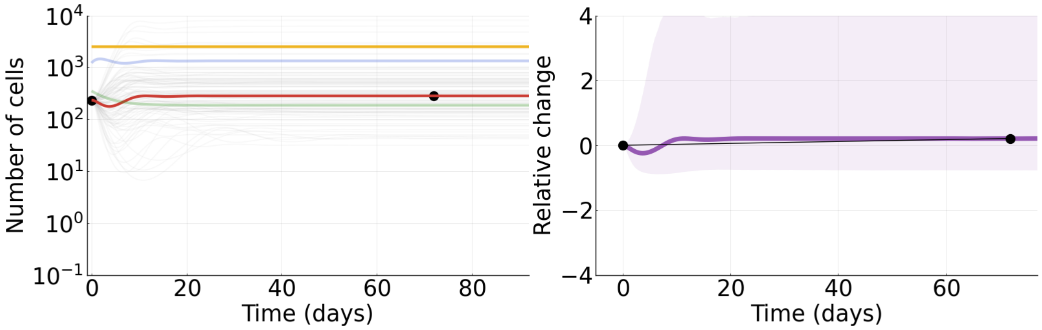


Figure S68: **Tumor 41.** Tumor data fit to mathematical model (model parameters set to be the median of their respective posterior distribution). Left panel: black dots represent the data, the red line represents tumor cells, the yellow line represents MDSCs, the green line represents NK cells, and the blue line represents CTL cells. Gray lines represent individual tumor cell population trajectories from the MCMC. Right panel: black dots represent the data, the purple line represents the tumor cell population, and the shaded area denotes the 90% credible interval where 90% of the posterior trajectories lie.

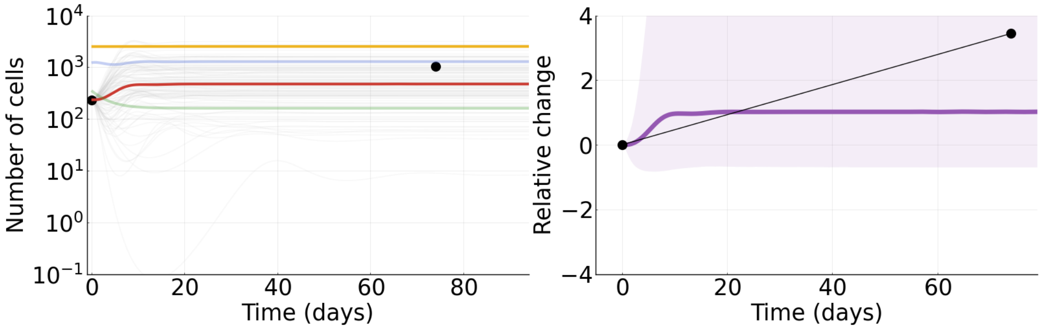


Figure S69: **Tumor 42.** Tumor data fit to mathematical model (model parameters set to be the median of their respective posterior distribution). Left panel: black dots represent the data, the red line represents tumor cells, the yellow line represents MDSCs, the green line represents NK cells, and the blue line represents CTL cells. Gray lines represent individual tumor cell population trajectories from the MCMC. Right panel: black dots represent the data, the purple line represents the tumor cell population, and the shaded area denotes the 90% credible interval where 90% of the posterior trajectories lie.

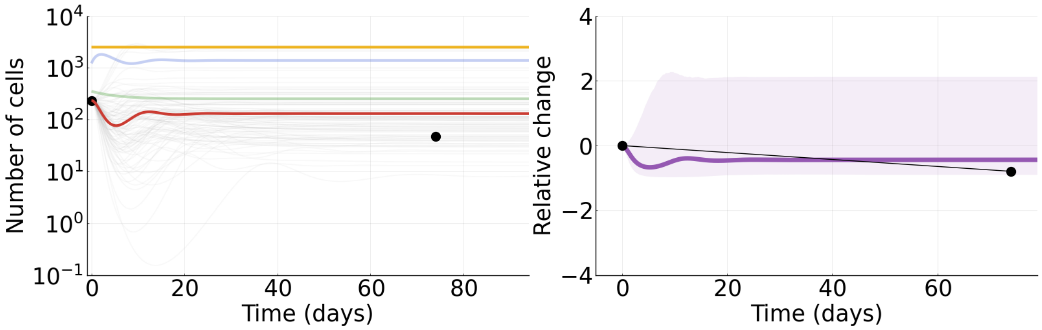


Figure S70: **Tumor 43.** Tumor data fit to mathematical model (model parameters set to be the median of their respective posterior distribution). Left panel: black dots represent the data, the red line represents tumor cells, the yellow line represents MDSCs, the green line represents NK cells, and the blue line represents CTL cells. Gray lines represent individual tumor cell population trajectories from the MCMC. Right panel: black dots represent the data, the purple line represents the tumor cell population, and the shaded area denotes the 90% credible interval where 90% of the posterior trajectories lie.

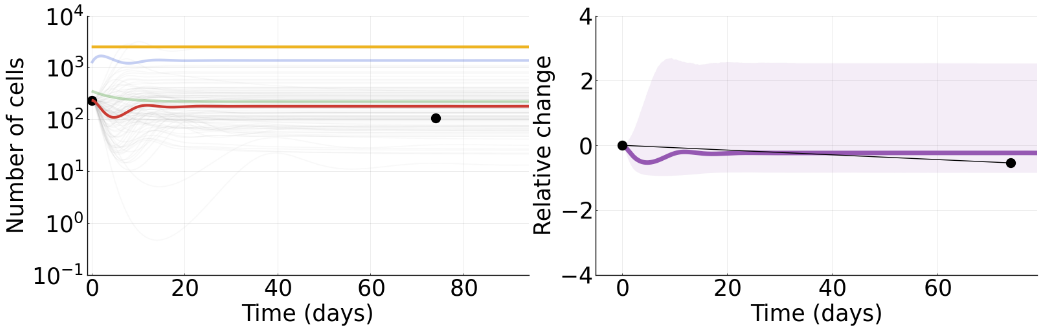


Figure S71: **Tumor 44.** Tumor data fit to mathematical model (model parameters set to be the median of their respective posterior distribution). Left panel: black dots represent the data, the red line represents tumor cells, the yellow line represents MDSCs, the green line represents NK cells, and the blue line represents CTL cells. Gray lines represent individual tumor cell population trajectories from the MCMC. Right panel: black dots represent the data, the purple line represents the tumor cell population, and the shaded area denotes the 90% credible interval where 90% of the posterior trajectories lie.

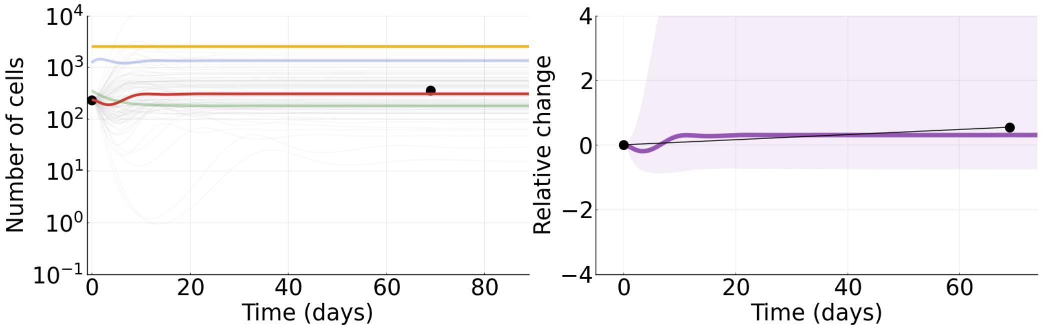


Figure S72: **Tumor 45.** Tumor data fit to mathematical model (model parameters set to be the median of their respective posterior distribution). Left panel: black dots represent the data, the red line represents tumor cells, the yellow line represents MDSCs, the green line represents NK cells, and the blue line represents CTL cells. Gray lines represent individual tumor cell population trajectories from the MCMC. Right panel: black dots represent the data, the purple line represents the tumor cell population, and the shaded area denotes the 90% credible interval where 90% of the posterior trajectories lie.

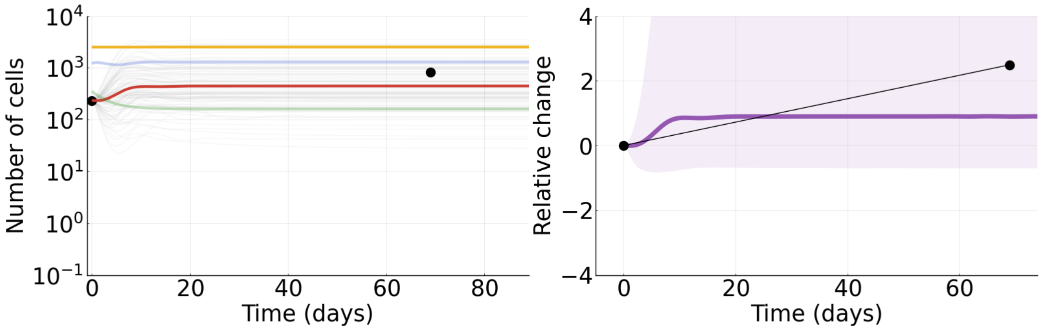


Figure S73: **Tumor 46.** Tumor data fit to mathematical model (model parameters set to be the median of their respective posterior distribution). Left panel: black dots represent the data, the red line represents tumor cells, the yellow line represents MDSCs, the green line represents NK cells, and the blue line represents CTL cells. Gray lines represent individual tumor cell population trajectories from the MCMC. Right panel: black dots represent the data, the purple line represents the tumor cell population, and the shaded area denotes the 90% credible interval where 90% of the posterior trajectories lie.

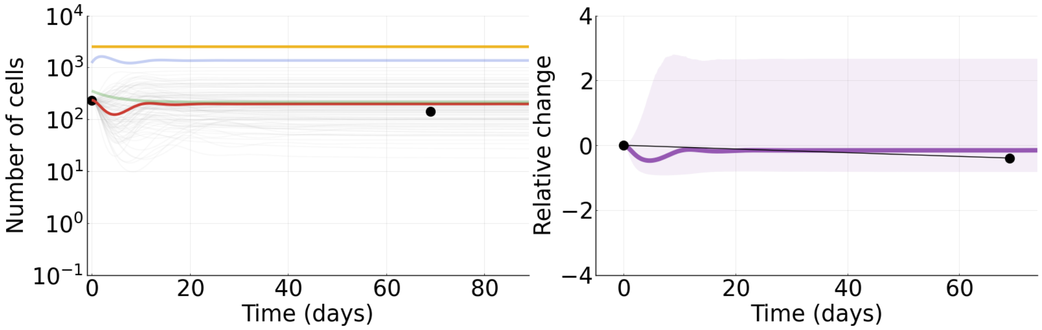


Figure S74: **Tumor 47.** Tumor data fit to mathematical model (model parameters set to be the median of their respective posterior distribution). Left panel: black dots represent the data, the red line represents tumor cells, the yellow line represents MDSCs, the green line represents NK cells, and the blue line represents CTL cells. Gray lines represent individual tumor cell population trajectories from the MCMC. Right panel: black dots represent the data, the purple line represents the tumor cell population, and the shaded area denotes the 90% credible interval where 90% of the posterior trajectories lie.

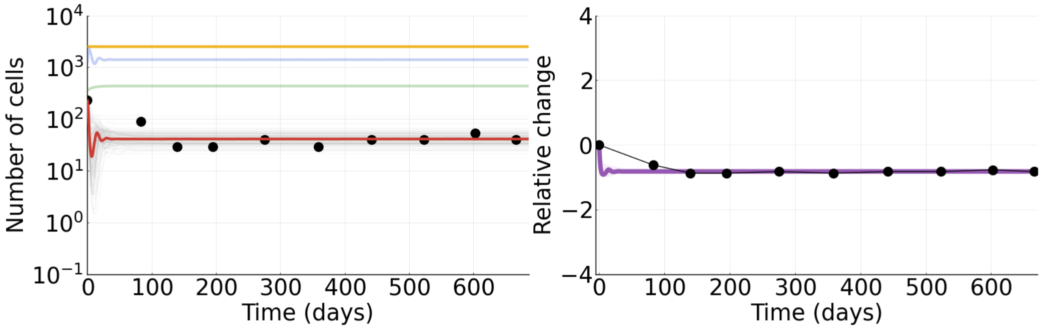


Figure S75: **Tumor 48.** Tumor data fit to mathematical model (model parameters set to be the median of their respective posterior distribution). Left panel: black dots represent the data, the red line represents tumor cells, the yellow line represents MDSCs, the green line represents NK cells, and the blue line represents CTL cells. Gray lines represent individual tumor cell population trajectories from the MCMC. Right panel: black dots represent the data, the purple line represents the tumor cell population, and the shaded area denotes the 90% credible interval where 90% of the posterior trajectories lie.

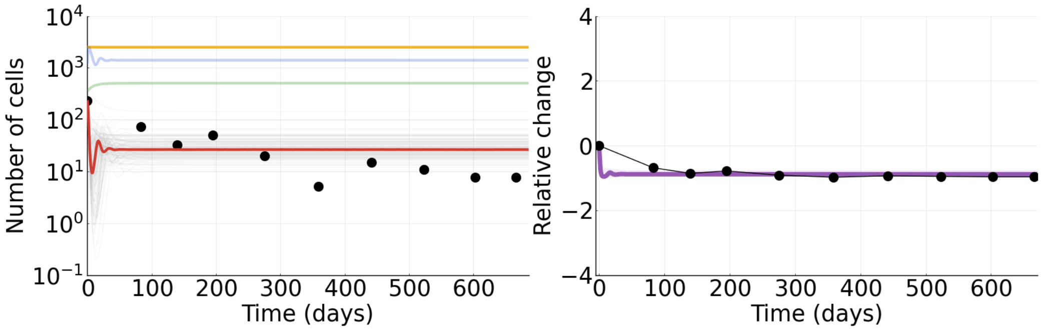


Figure S76: **Tumor 49.** Tumor data fit to mathematical model (model parameters set to be the median of their respective posterior distribution). Left panel: black dots represent the data, the red line represents tumor cells, the yellow line represents MDSCs, the green line represents NK cells, and the blue line represents CTL cells. Gray lines represent individual tumor cell population trajectories from the MCMC. Right panel: black dots represent the data, the purple line represents the tumor cell population, and the shaded area denotes the 90% credible interval where 90% of the posterior trajectories lie.

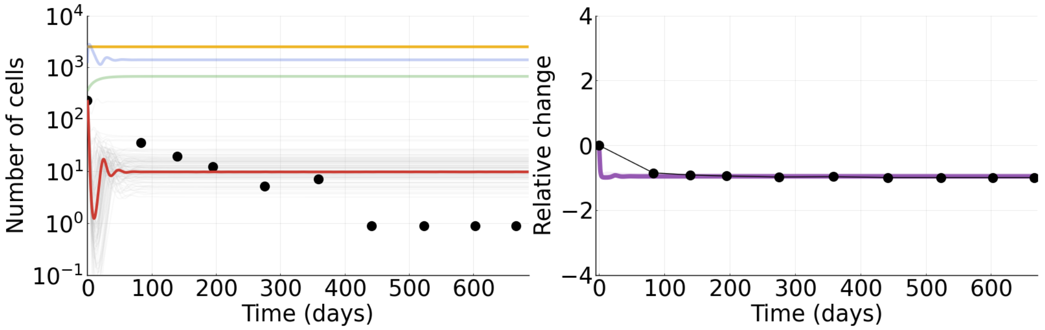


Figure S77: **Tumor 50.** Tumor data fit to mathematical model (model parameters set to be the median of their respective posterior distribution). Left panel: black dots represent the data, the red line represents tumor cells, the yellow line represents MDSCs, the green line represents NK cells, and the blue line represents CTL cells. Gray lines represent individual tumor cell population trajectories from the MCMC. Right panel: black dots represent the data, the purple line represents the tumor cell population, and the shaded area denotes the 90% credible interval where 90% of the posterior trajectories lie.

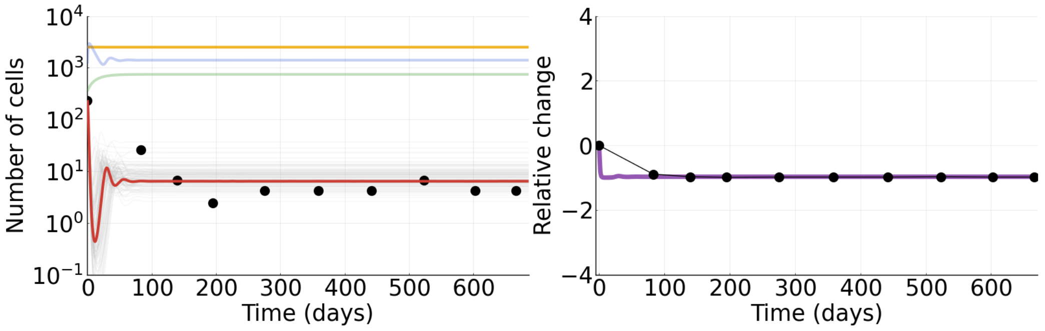


Figure S78: **Tumor 51.** Tumor data fit to mathematical model (model parameters set to be the median of their respective posterior distribution). Left panel: black dots represent the data, the red line represents tumor cells, the yellow line represents MDSCs, the green line represents NK cells, and the blue line represents CTL cells. Gray lines represent individual tumor cell population trajectories from the MCMC. Right panel: black dots represent the data, the purple line represents the tumor cell population, and the shaded area denotes the 90% credible interval where 90% of the posterior trajectories lie.

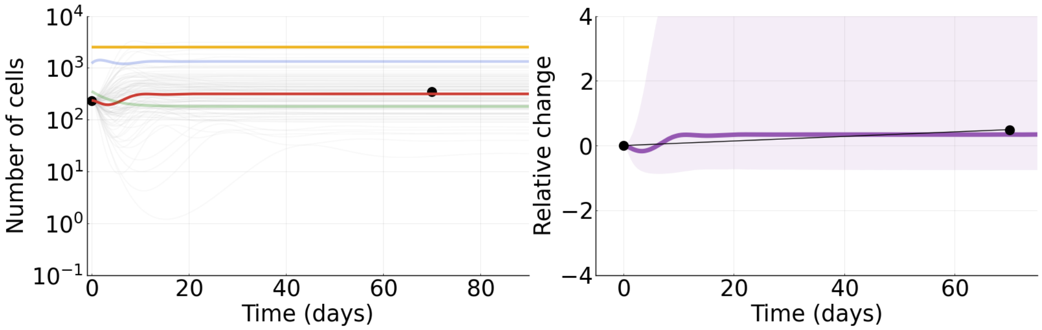


Figure S79: **Tumor 52.** Tumor data fit to mathematical model (model parameters set to be the median of their respective posterior distribution). Left panel: black dots represent the data, the red line represents tumor cells, the yellow line represents MDSCs, the green line represents NK cells, and the blue line represents CTL cells. Gray lines represent individual tumor cell population trajectories from the MCMC. Right panel: black dots represent the data, the purple line represents the tumor cell population, and the shaded area denotes the 90% credible interval where 90% of the posterior trajectories lie.

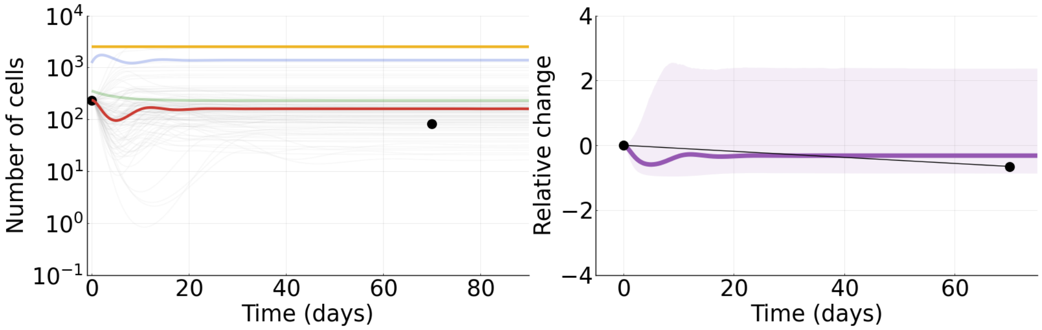


Figure S80: **Tumor 53.** Tumor data fit to mathematical model (model parameters set to be the median of their respective posterior distribution). Left panel: black dots represent the data, the red line represents tumor cells, the yellow line represents MDSCs, the green line represents NK cells, and the blue line represents CTL cells. Gray lines represent individual tumor cell population trajectories from the MCMC. Right panel: black dots represent the data, the purple line represents the tumor cell population, and the shaded area denotes the 90% credible interval where 90% of the posterior trajectories lie.

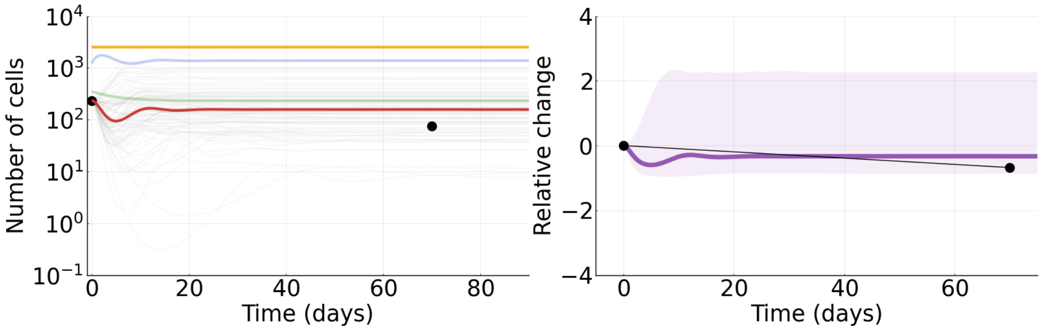


Figure S81: **Tumor 54.** Tumor data fit to mathematical model (model parameters set to be the median of their respective posterior distribution). Left panel: black dots represent the data, the red line represents tumor cells, the yellow line represents MDSCs, the green line represents NK cells, and the blue line represents CTL cells. Gray lines represent individual tumor cell population trajectories from the MCMC. Right panel: black dots represent the data, the purple line represents the tumor cell population, and the shaded area denotes the 90% credible interval where 90% of the posterior trajectories lie.

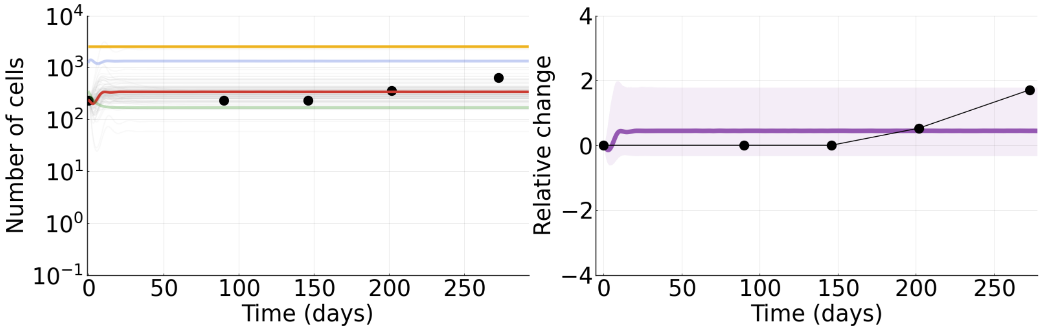


Figure S82: **Tumor 55.** Tumor data fit to mathematical model (model parameters set to be the median of their respective posterior distribution). Left panel: black dots represent the data, the red line represents tumor cells, the yellow line represents MDSCs, the green line represents NK cells, and the blue line represents CTL cells. Gray lines represent individual tumor cell population trajectories from the MCMC. Right panel: black dots represent the data, the purple line represents the tumor cell population, and the shaded area denotes the 90% credible interval where 90% of the posterior trajectories lie.

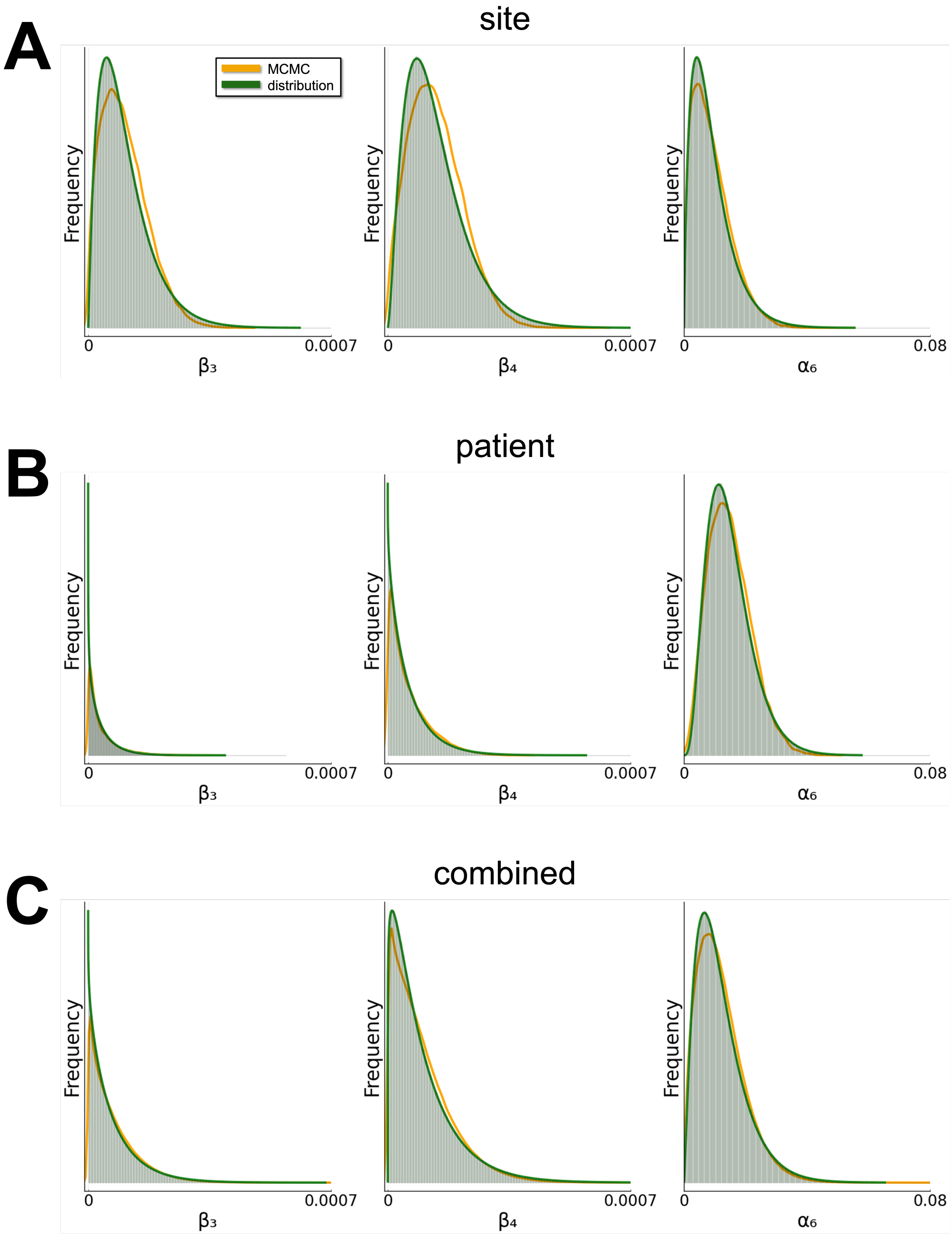


Figure S83: **Posterior distributions for simulations.** Example of posterior distributions for $\beta_3$, $\beta_4$, and $\alpha_6$ for simulations. Yellow lines represent the density of the posterior parameter values, green lines represent the fitted Gamma distributions, and the green shading represents the density of $10^6$ samples from the fitted Gamma distribution. **A**. Liver. **B**. Patient R-18. **C**. Combined. See Methods for details.

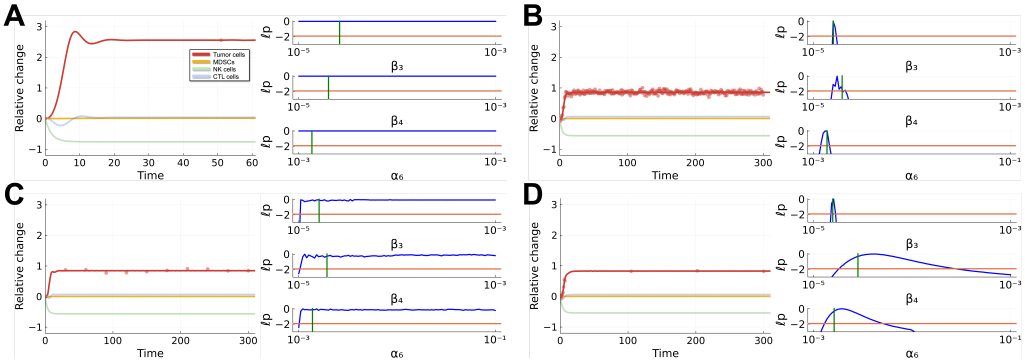


Figure S84: **Parameter identifiability is constrained by the temporal density of tumor data for increasing tumors.** Same as Fig. 6 in the main text but for an increasing tumor. **A**. Actual tumor data from clinical trial (tumor 10). **B**. Simulated tumor data every day. All three parameters are identifiable and well-constrained. **C**. Simulated tumor data at 11 equally spaced points (every 30 days) over the timespan. **D**. Simulated tumor data every day for the first week after treatment initiation and then three equally spaced points after that, same number of data points as panel C.

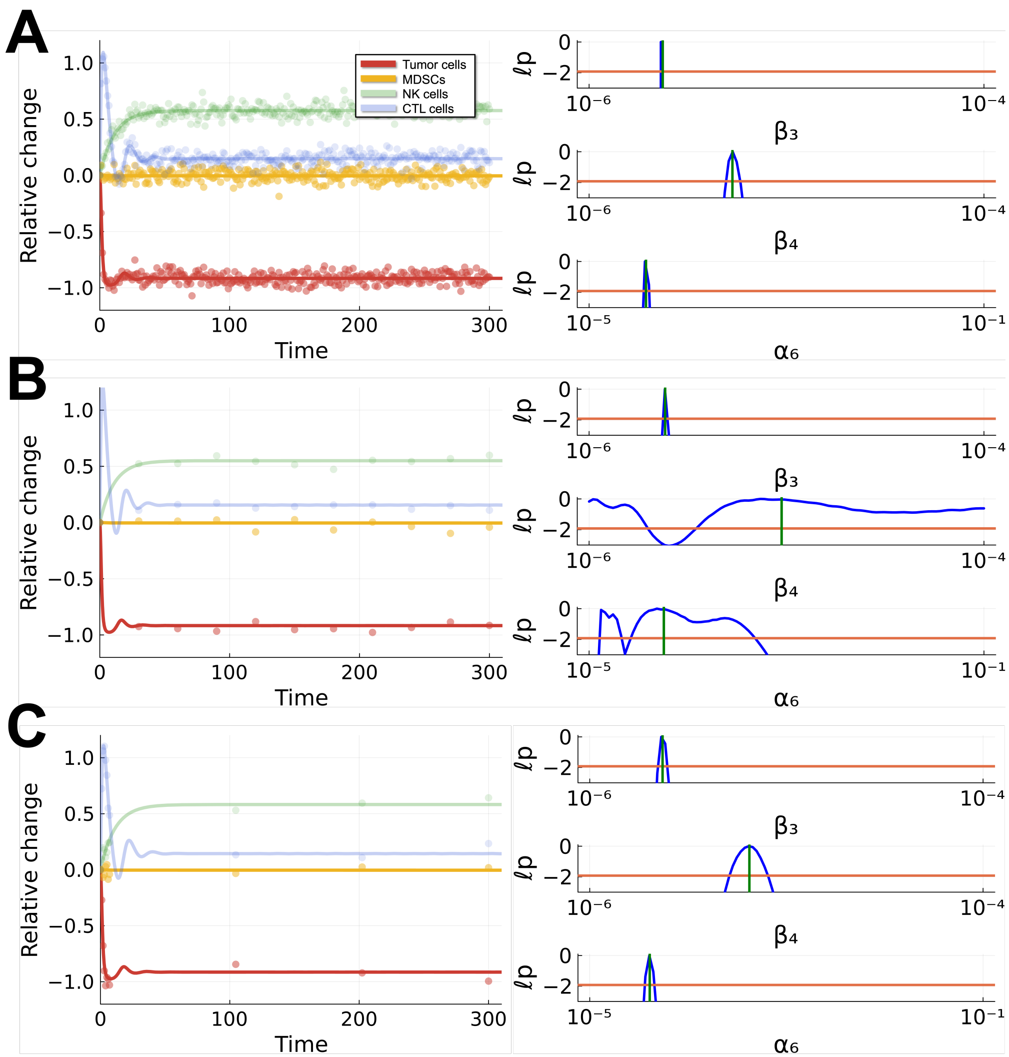


Figure S85: **Parameter identifiability for decreasing tumors improves with data from all four model populations.** Same as Fig. 6 in the main text but data simulated from all four model populations. **A**. Simulated data every day. All three parameters are identifiable and well-constrained. **B**. Simulated data at 11 equally spaced points over the timespan. **C**. Simulated data every day for the first week after treatment initiation and then three equally spaced points after that, same number of data points as panel B.

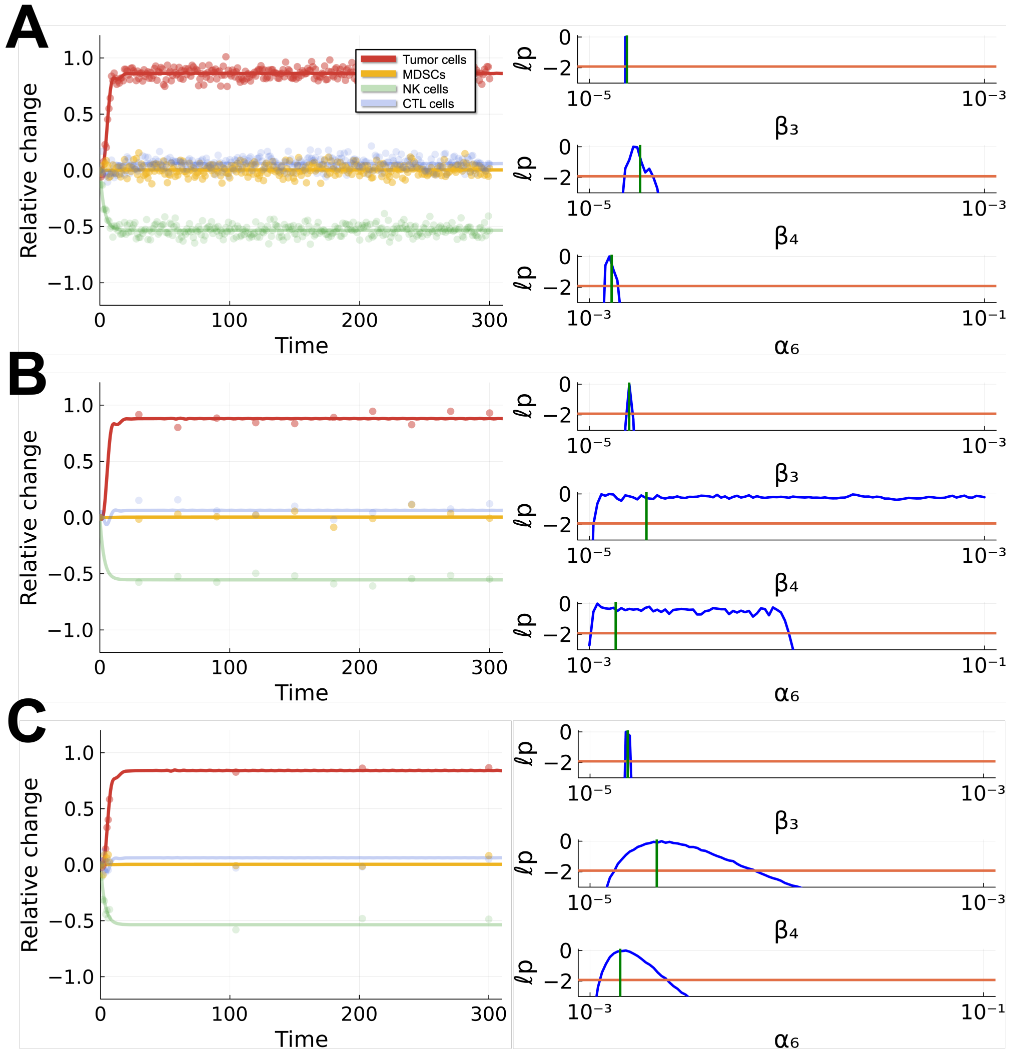


Figure S86: **Parameter identifiability for increasing tumors improves with data from all four model populations.** Same as Fig. S84 but data simulated from all four model populations. **A**. Simulated data every day. All three parameters are identifiable and well-constrained. **B**. Simulated data at 11 equally spaced points over the timespan. **C**. Simulated data every day for the first week after treatment initiation and then three equally spaced points after that, same number of data points as panel B.
